# More than just an Eagle Killer: The freshwater cyanobacterium *Aetokthonos hydrillicola* produces highly toxic dolastatin derivatives

**DOI:** 10.1101/2023.04.12.536103

**Authors:** Markus Schwark, José A. Martínez Yerena, Kristin Röhrborn, Pavel Hrouzek, Petra Divoká, Lenka Štenclová, Kateřina Delawská, Heike Enke, Christopher Vorreiter, Faith Wiley, Wolfgang Sippl, Roman Sobotka, Subhasish Saha, Susan B. Wilde, Jan Mareš, Timo H. J. Niedermeyer

## Abstract

Cyanobacteria are infamous producers of toxins. While the toxic potential of planktic cyanobacterial blooms is well documented, the ecosystem level effects of toxigenic benthic and epiphytic cyanobacteria are an understudied threat. The freshwater epiphytic cyanobacterium *Aetokthonos hydrillicola* has recently been shown to produce the “eagle killer” neurotoxin aetokthonotoxin causing the fatal neurological disease Vacuolar Myelinopathy. The disease affects a wide array of wildlife in the southeastern United States, most notably waterfowl and birds of prey, including the bald eagle. In an assay for cytotoxicity, we found the crude extract of the cyanobacterium to be much more potent than pure aetokthonotoxin, prompting further investigation. Here, we describe the isolation and structure elucidation of the aetokthonostatins, linear peptides belonging to the dolastatin compound family, featuring a unique modification of the C-terminal phenylalanine derived moiety. Using immunofluorescence microscopy and molecular modeling, we confirmed that aetokthonostatin acts as a potent tubulin binder. We also show that aetokthonostatin inhibits reproduction of the nematode *C. elegans*, resulting in increased population lethality of the combined action of the two toxins produced by *A. hydrillicola*. Bioinformatic analysis revealed the aetokthonostatin biosynthetic gene cluster encoding a non-ribosomal peptide synthetase/polyketide synthase accompanied by a unique tailoring machinery. The biosynthetic activity of a specific N-terminal methyltransferase was confirmed by *in vitro* biochemical studies, establishing a mechanistic link between the gene cluster and its product.

**Significance Statement:** Cyanotoxins have adverse effects on ecosystems. Our understanding of their potential risk has recently been expanded by the discovery of aetokthonotoxin, produced by the cyanobacterium *Aetokthonos hydrillicola* growing on invasive plants. Via trophic transfer, it acts as a neurotoxin causing mortality in animals including top predators like Bald Eagles. Closer examination of *A. hydrillicola* revealed that it also produces highly toxic dolastatin derivatives. *A. hydrillicola* is the first cultured cyanobacterium producing dolastatin derivatives, allowing us to uncover biosynthetic gene clusters of this compound family. In contrast to all other known dolastatin-producers, which are marine cyanobacteria, *A. hydrillicola* thrives in freshwater reservoirs, making it a potential threat also for human health. Monitoring of the cyanobacterium and its toxins is strongly recommended.

## Introduction

Cyanobacteria are a prolific source of biologically active specialized metabolites (1–3). They are infamous for their potent toxins, such as the neurotoxic alkaloids anatoxins or the hepatotoxic non-ribosomal peptides microcystins (4, 5). Harmful cyanobacterial blooms (mass proliferation of toxic cyanobacteria in stagnant waters) can pose a threat for ecosystem health (6). While planktic cyanobacteria have been studied intensely in the past, the effects of benthic or epiphytic cyanobacteria remain largely unknown, although their cyanotoxin repertoire is equally high, and they have been linked to numerous animal poisonings (7). Recently, a novel cyanobacterial toxin was discovered from the filamentous epiphytic cyanobacterium *Aetokthonos hydrillicola* (8). The biindole alkaloid aetokthonotoxin (AETX, Fig. 1A) has been found to be the cause of the disease Vacuolar Myelinopathy (VM), affecting wildlife in the southeastern United States. *A. hydrillicola* grows on the invasive submersed aquatic plants, especially *Hydrilla verticillata*, which is consumed by waterfowl, fish, and snails, which in turn are consumed by birds of prey. AETX consumption resulted in VM throughout every trophic level of this aquatic food chain (8).

**Figure 1.**
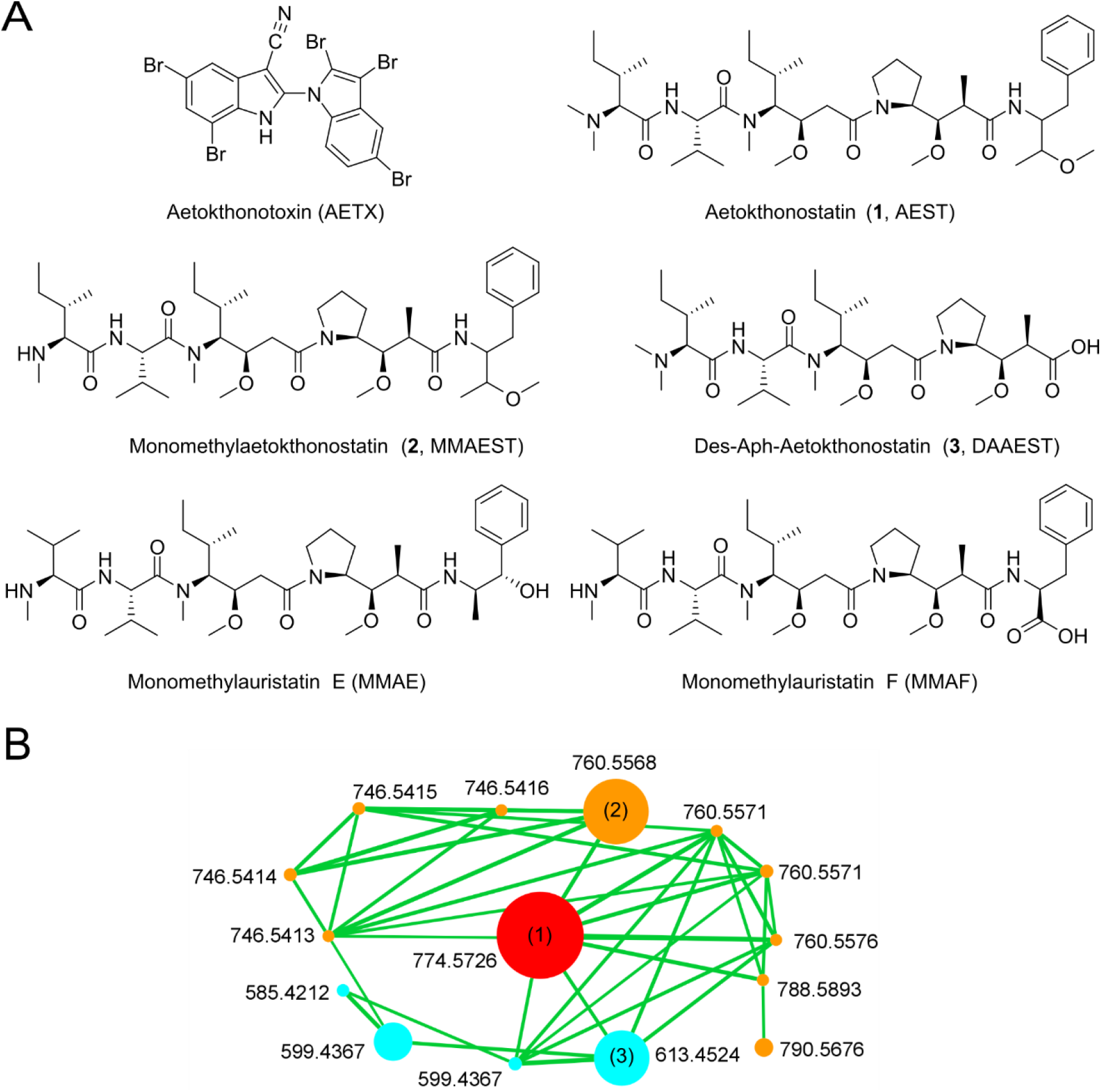
*Aetokthonos hydrillicola* produces dolastatin analogs. **(A)** Structures of aetokthonotoxin (AETX), aetokthonostatin (AEST, **1**), monomethylaetokthonostatin (MMAEST, **2**), and des-Aph-aetokthonostatin (DAAEST, **3**), as well as monomethylauristatins E and F (MMAE, MMAF). **(B)** Aetokthonostatin cluster in a GNPS feature-based MS/MS networking analysis of an HPLC-MS/MS analysis of an *A. hydrillicola* extract. Node size proportional to ion intensity, exact mass of [M+H]^+^ indicated next to the respective node. Red: AEST, orange: pentapeptides, cyan: tetrapeptides.

Biologically active cyanobacterial specialized metabolites provide opportunities for drug substance development (1, 9, 10). Prominent examples that have advanced into clinical studies are saxitoxins as long-lasting local anesthetics (11, 12), or cryptophycins as potential antineoplastic drugs (13, 14). However, the only cyanobacterial natural products of which derivatives have been clinically tested are the dolastatins (1, 9). These linear non-ribosomal pentapeptides are structurally characterized by two uncommon amino acids only present in the dolastatin compound family, dolaproine and dolaisoleuine (15). Dolastatins as well as structurally related peptides of cyanobacterial origin, e.g. the lyngbyastatins (16), symplostatins (17), or malevamide D (18), are highly cytotoxic (low to sub-nM EC_50_). Binding to the vinca binding site of the β1 subunit of tubulin, they interrupt microtubule assembly, resulting in cell cycle arrest in the G_2_/M phase (19–21). Synthetic derivatives of dolastatin 10, monomethylauristatin E (MMAE) and monomethylauristatin F (MMAF), have been introduced into the clinic as highly cytotoxic payloads of antineoplastic antibody-drug conjugates. The first of these, brentuximab vedotin, has been approved by the US FDA in 2011 (9).

The dolastatins were first described from the sea hare *Dolabella auricularia* by Petitt *et al*. in 1987 (22). However, later studies found that the sea hare acquires the dolastatins by their diet, which consists largely of cyanobacteria of the genus *Symploca*, the actual producers of these compounds (23, 24). Notably, all dolastatin derivatives known to date have been isolated from marine cyanobacteria such as *Symploca* or *Lyngbya*/*Moorena* (16). Marine benthic filamentous cyanobacteria have been intensively studied as an extremely rich source of bioactive compounds (25). However, no dolastatin-producing strain has been cultured yet, hampering the elucidation of the genetic basis for the compound’s biosynthesis as well as synthetic biology approaches for their modification. As well established total synthesis routes to the dolastatins are available, ongoing research on the medicinal chemistry of dolastatins focuses on synthetic analogs (26).

Profiling the bioactivity of purified AETX and crude extracts of the cyanobacterium *A. hydrillicola*, we observed that the cytotoxicity of a crude extract of *A. hydrillicola* biomass was much stronger than the cytotoxicity of a corresponding amount of pure AETX. Thus, there had to be another compound in the extract with higher cytotoxicity than AETX. Intrigued by these results, we set out to isolate and characterize the cytotoxic compound produced by *A. hydrillicola*. Here, we present the isolation and structure elucidation of aetokthonostatin (AEST) and AEST derivatives. The aetokthonostatins are linear peptides belonging to the dolastatin compound family, featuring a yet undescribed C-terminal phenylalanine-derived building block. Using immunofluorescence microscopy and molecular modeling, we confirmed that AEST exerts its strong cytotoxicity by binding to tubulin, comparable to the dolastatins. We show that AEST inhibits reproduction of the nematode *C. elegans*. Bioinformatic analysis of the *A. hydrillicola* genome revealed the biosynthetic gene cluster of AEST, which we confirmed by biochemical experiments. Our study highlights that the cyanobacterium *A. hydrillicola* might pose a serious threat to drinking-water supplies, as it is able to produce two specialized metabolites with synergistic pronounced toxicity.

## Results and Discussion

### Toxicity of *A. hydrillicola* Extract and AETX

Early investigation of a crude extract of the cyanobacterium *A. hydrillicola* for cytotoxicity showed a remarkably high activity of the extract (EC_50_ 0.12 µg/mL, HeLa, Fig. S2). Focusing on the isolation of the toxin causing VM, we assumed that the toxin causing VM would also be responsible for the extract’s cytotoxicity. After isolation of AETX and testing the pure compound for cytotoxicity, to our surprise, we found that AETX is only moderately cytotoxic (EC_50_ 1 µM = 0.65 µg/mL, HeLa). Quantification of the AETX content in a crude extract assayed for cytotoxicity showed that, based on the determined AETX content, the extract was about 5-fold more cytotoxic than expected if AETX was the sole cytotoxic compound in the extract. This suggested that another, more cytotoxic compound must be present in the extract. Indeed, microfractionation of the extract and subsequent cytotoxicity testing revealed that not AETX, but the main compound observed in the extract’s HPLC chromatogram was highly cytotoxic (Fig. S3).

### Isolation and Structure Elucidation of Aetokthonostatin

In contrast to AETX, which is only produced by *A. hydrillicola* when the strain’s cultivation medium is supplemented with bromide salts (8), we found the cytotoxic metabolite to be produced under all tested cultivation conditions. Thus, the cyanobacterium was cultivated in standard BG-11 medium to generate biomass for subsequent processing. After harvest and lyophilization, the biomass was extracted with methanol. The crude extract was fractionated using flash chromatography. Fractions were assayed for cytotoxicity, and the cytotoxic fractions were subjected to semi-preparative HPLC to isolate the active compounds. We found that the main peak observed in the chromatogram (Fig. S3) consisted of two individual compounds (**1** and **2**), which we subsequently separated using ammonium acetate buffer (pH 9) in the mobile phase.

Compound **1** was isolated as a white amorphous powder. High resolution mass spectrometry suggested the molecular formula C_43_H_76_O_7_N_5_ ([M+H]^+^ at *m/z* 774.5759. NMR analysis (Fig. S5-S10) was complicated by a considerable signal overlap in the ^1^H NMR spectrum, caused by the presence of two conformers in solution (ratio main to minor conformer about 3:2, Fig. S5). Analysis of the ^13^C and ^1^H NMR data suggested four amide or ester carbonyls and two NH protons (NMR data see Tab. S1). The presence of amide bonds was confirmed by four signals agreeing with α-carbonyl protons of amino acids, giving us confidence that **1** was a peptide. Furthermore, we observed signals of six N- or O-methyl groups. Evaluation of HSQC-DEPT, TOCSY and COSY spectra allowed us to assemble the spin systems of the amino acid building blocks in the peptide (for a more detailed discussion, see SI). In addition to the amino acids valine and isoleucine, we found that **1** contains the known non-proteinogenic amino acids dolaisoleuine (Dil) and dolaproine (Dap), indicating that the compound belongs to the dolastatin compound family. Presence of Dap explained the existence of two conformers in solution, which has also been described for other compounds in the dolastatin compound family (27). We could assemble the fifth monomer as 3-methoxy-1-phenylbutan-2-amine, a yet undescribed monomer in peptide natural products, that we give the trivial name aetophenine (Aph). The building block sequence as well as the position of the N- and O-methyl groups was deduced from an HMBC spectrum and confirmed by a ROESY spectrum. The planar structure of **1** was thus established to be *N,N*-dimethyl-Val-Ile-Dil-Dap-Aph. This sequence was supported by the fragmentation pattern in MS/MS experiments (Fig. S11). As this planar structure differs from the structure of symplostatin 1 only by the C-terminal monomer (23), we rerecorded a ^13^C-NMR spectrum of **1** in CD_2_Cl_2_ (Fig. S6b-c). Indeed, the NMR data of the four N-terminal monomers of **1** matched to those reported for symplostatin 1 (23), indicating that these four monomers have the same relative and absolute stereochemistries in both compounds (Tab. S4). To confirm this assumption, the absolute stereochemistries of the fourth monomer, Dap, was determined by Marfey’s analysis (Fig. S12). As no reference material for Aph is available, we first tried to hydrolyze **1** and isolate Aph for subsequent VCD spectroscopic analysis. However, we found that under the strong acidic or basic conditions needed for hydrolysis, the methyl ether was hydrolyzed as well, obfuscating the original configuration. Crystallization of **1** for X-ray crystallography was not successful. The absolute configuration of Aph could thus not be determined. The structure of **1** is shown in Fig. 1A. **1**, the main compound produced by *A. hydrillicola*, was named aetokthonostatin (AEST).

Compound **2** was isolated as a white amorphous powder. Its molecular formula was established as C_42_H_74_O_7_N_5_ ([M+H]^+^ at *m/z* 760.5571), differing from **1** by only by the absence of a single methyl group. Analysis of the MS/MS and NMR data of **2** confirmed that, compared to **1, 2** lacks one of the two N-methyl groups at the N-terminal valine (Figs. S13-S19, Tab. S2). Thus, this compound has been identified as monomethylaetokthonostatin (MMAEST).

Compound **3**, a white amorphous powder, eluted earlier than **1** and **2**, indicating higher polarity or smaller size. Indeed, its molecular formula, C_32_H_61_O_7_N_4_ ([M+H]^+^ at *m/z* 613.4535), indicated **3** is constituted of only four amino acids. The mass difference between **3** and **1** corresponds to the C-terminal monomer of **1**, Aph. MS/MS data as well as NMR data supported the hypothesis that this monomer was missing, as no signals were observed in the aromatic region in the ^1^H spectrum of **2** while the other signals agree with those in the spectra of **1** (Figs. S20-S23, Tab. S3), and the prominent fragment ion at 180.13 *m/z* could not be detected (Fig. S24). **3** was thus confirmed to be, *N,N*-dimethyl-Val-Ile-Dil-Dap, des-Aph-aetokthonostatin (DAAEST).

### Aetokthonostatin Derivatives in an *A. hydrillicola* Extract

In addition to these three isolated compounds, feature-based MS/MS networking analysis of HPLC-MS/MS data using GNPS (28) allowed us to detect additional aetokthonostatin derivatives in an *A. hydrillicola* extract (Fig. 1B). These derivatives differ in the methylation pattern and the number of the amino acids. Based on the ion intensity, AEST (**1**, *m/z* 774.5726) is the most abundant statin derivate in the extract, followed by its N-terminal desmethyl derivative **2** (*m/z* 760.5568), its des-Aph derivative **3** (*m/z* 613.4524), and the respective N-terminal desmethyl-des-Aph derivative (*m/z* 599.4367). Interestingly, four possible des-methyl derivatives of **1** as well as two of **3** could be detected as well as four derivatives of **1** and one derivative of **3** lacking two methyl groups. One derivative with an additional methyl group was observed (*m/z* 788.5893). While the pentapeptides represent the major part of the AEST derivatives, significant amounts of the respective tetrapeptides are present in the extract. No di- or tripeptides could be detected. The structures of most of these compounds could be deduced based on their MS/MS spectra (Fig. S25-S42).

### Bioactivity of Aetokthonostatin

Structure elucidation of AEST already explained the exquisite cytotoxicity of the compound: It belongs to the dolastatin compound family. Dolastatins and related compounds are known for their ability to arrest cells in G_2_/M phase due to binding to tubulin and disrupting its assembly to microtubules (19–21). The cytotoxicity of AEST was assessed against HeLa cells and the triple-negative breast cancer cell line MDA-MB 231, and compared to the cytotoxicity of monomethyl auristatin E (MMAE) and monomethyl auristatin F (MMAF), which are used as payloads in approved antibody-drug conjugates (29, 30). We found that AEST (EC_50_ 1 ± 0.2 nM) is equally potent as MMAE (EC_50_ 3 ± 0.4 nM) against HeLa cells (31). Both are about 170-fold more active than MMAF (EC_50_ 170 ± 30 nM; Fig. S4A) against MDA-MB 231 cells (31). In HeLa cells, this effect was less distinct, AEST and MMAE were 10-fold more active (Fig. 2A).

**Figure 2.**
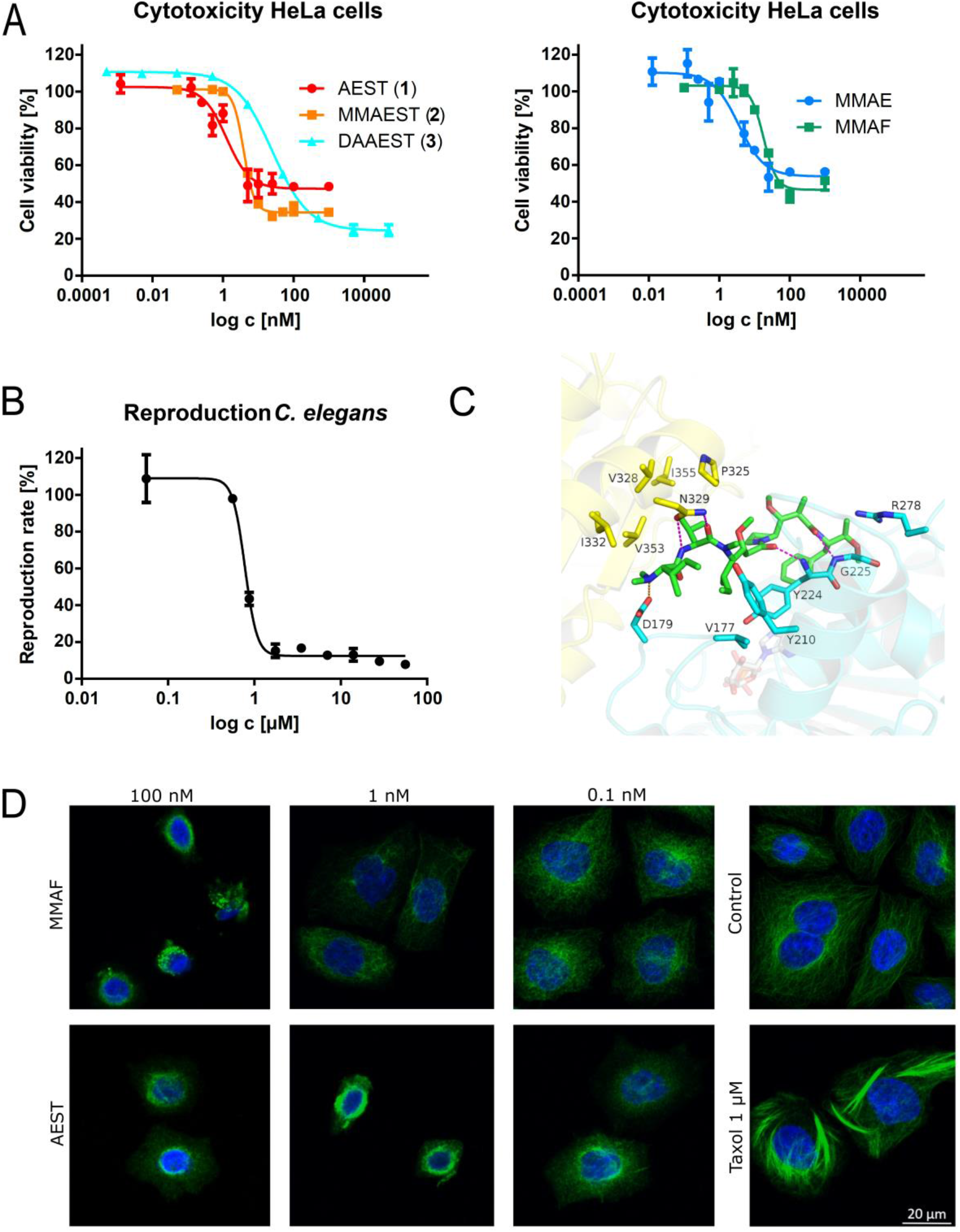
Aetokthonostatin is a cytotoxic tubulin binder. **(A)** Cytotoxicity of AEST (**1**, EC_50_ 1 ± 0.2 nM), AEST (**2**, EC_50_ 4 ± 0.01 nM), DAAEST (**3**, EC_50_ 25 ± 1.4 nM), MMAE (EC_50_ 3 ± 0.4 nM), and MMAF (EC_50_ 170 ± 30 nM) on HeLa cells (n = 3). Data are represented as average ± SEM. **(B)** Decrease of the reproduction rate of *C. elegans* treated with AEST (**1**, EC_50_ 0.8 ± 0.2 µM) (n = 3). Data are represented as average ± SEM. **(C)** Predicted binding of AEST (Aph as 2*S*,3*R*) to tubulin. Tubulin α-subunit visualized as yellow cartoon, β-subunit in cyan. Interacting binding site residues represented as sticks colored the same way. Magenta: hydrogen bonds, orange: salt-bridges, red: cation-π interactions. Cofactor GDP shown as white, transparent sticks. **(D)** Immunofluorescence microscopy of HeLa cells showing the effect of AEST and MMAF on tubulin (green); nuclei were stained with DAPI (blue); taxol (1 nM) was used as positive control.

Due to its negative charge under physiological pH conditions, auristatins possessing a carboxy group at the C-terminal monomer have lower cell permeability, explaining the lower *in vitro* potency of MMAF compared to MMAE (32). AEST, like MMAE, features an uncharged C-terminus. Therefore, it belongs to the MMAE-group of dolastatin analogs, which show higher membrane permeability and thus lower EC_50_ values. Lack of the methyl group at the N-terminal nitrogen in **2** does not have noticeable influence on potency against HeLa cells (EC_50_ 4 ± 0.01 nM; Fig. 2A). We found that the C-terminal monomer of **1** is not mandatory for cytotoxicity, as **3** is still strongly cytotoxic (EC_50_ 25 ± 1.4 nM; Fig. 2A). This finding was surprising, as it is assumed that a phenyl moiety at the C-terminus is mandatory for cytotoxicity (33, 34). However, as for example the clinically tested dolastatin analog tasidotin also does not contain an aromatic C-terminus but is still cytotoxic, the effect of the C-terminus might have been overrated (35). The reduced cytotoxicity of **3** compared to AEST is most likely caused by decreased cell penetration because of the free carboxylic acid. The residual survival rate was about 50% in the case of AEST even at high concentrations (Fig. 2A). This dose-response behavior is similar to that observed for MMAE (30). Interestingly, **2**, and even more pronounced **3**, had a lower residual cell viability (about 30% and 20%, resp.).

In early experiments with *Caenorhabditis elegans* and *Danio rerio*, we observed that the crude *A. hydrillicola* extract seemed to be more toxic than isolated AETX. Having identified AEST as a second toxin in the extract, we wondered whether this observed higher toxicity was due to an additive effect of the two toxins, or if they might act synergistically. However, in stark contrast to AETX, but in agreement with previous findings on dolastatin 15 (36), aetokthonostatin showed no acute toxic effect on *C. elegans* even at high concentrations of up to 56 µM. Instead, a decrease in the nematode’s reproduction rate could be observed at low concentration (EC_50_ 0.8 ± 0.2 µM, Fig. 2B). Thus, *A. hydrillicola* produces two toxins displaying different types of toxicity. In case of *C. elegans*, this results in a more effective inhibition: If a *C. elegans* population is treated with AETX, the surviving worms usually reproduce, and after some time, the population is reestablished. However, when in parallel reproduction is inhibited by AEST, the population dies out even if individual nematodes survive the AETX exposure.

As dolastatins are well-known tubulin inhibitors binding at the vinca domain, we performed a docking study to test whether AEST binds to tubulin in a similar fashion to known dolastatin derivatives. As the absolute configuration of the two stereocenters of Aph could not yet be determined, we used all four possible stereoisomers in this study. Using a crystal structure of a dolastatin analog cocrystallized with tubulin as template (Protein Data Bank ID 4X1K), we found that both AEST and MMAF can nicely be fit into the respective tubulin binding site (Fig. 2B; for a detailed discussion of the docking study see SI). The carbonyl oxygen atoms of Dil and Dap form hydrogen bonds to the backbone amide NH groups of Tyr β224 and Gly β225, respectively. The carbonyl oxygen as well as the amide NH of Val form a bifocal hydrogen bond network with the side chain of Asn α329 (Fig. 2B). The charged N-terminal dimethylamino group shows electrostatic interactions with Asp β179. The C-terminal benzyl group showed different orientations in the docking poses (either buried towards Gly α225 or cation-π interaction with Arg β278). The flexibility of the C-terminal group was also observed in the molecular dynamics simulations of AEST and structurally similar cocrystalized peptides, where the C-terminal benzyl moiety is moving significantly during the simulations (Fig. S46). This agrees with our finding that **3**, lacking the Aph unit, is still strongly cytotoxic. Also, it suggests that the absolute configuration of the stereocenters of Aph is not important for tubulin binding.

We subsequently confirmed inhibition of tubulin polymerization by AEST experimentally by immunofluorescence staining (Fig. 2D). Both AEST and MMAF were assessed in this assay. As expected, the tubulin network appeared affected by AEST at a concentration well below the EC_50_ value (observable effect at 0.1 nM) and proceeding towards higher concentrations, while for MMAF, the tubulin network morphology was comparable to untreated control cells at concentrations around 1 nM. Similar data were obtained with MDA-MB 231 cells, with the activity windows shifted towards higher concentrations (Fig. S4B).

### Biosynthesis of Aetokthonostatin

Surprisingly, although dolastatin derivatives have been the first and are still today the only approved drugs based on cyanobacterial specialized metabolites, their biosynthesis has not yet been studied. In the genome of *A. hydrillicola* Thurmond2011, we located a 36.94 kbp long non-ribosomal peptide synthetase/polyketide synthase (NRPS/PKS) gene cluster (GenBank accession number ON840103) consisting of twelve genes with deduced functions in AEST biosynthesis (Fig. 3A, Table S5). It was found in the middle of a 472 kbp long contig of clearly cyanobacterial origin. The putative *aes* biosynthetic gene cluster (BGC) included four NRPS genes (*aesA, C, F*, and *G*), harboring altogether five amino acid incorporating modules (Fig. 3B). The predicted substrate specificities of all the respective adenylation domains and the occurrence of two additional *N*-methyltransferase domains agreed with the observed amino acid sequence of AEST. Specifically, a starter unit of AesF was predicted to activate isoleucine and methylate it to form an *N*-terminal *N*-methylisoleucine residue, followed by the incorporation of valine and another *N*-methylisoleucine by two downstream NRPS modules of AesG. According to our prediction, the subsequent biosynthetic steps are not fully colinear with the arrangement of *aes* genes, analogously to other reported NRPS/PKS biosyntheses (38–40). We suggest that in the next step, the PKS enzyme AesJ, which contains a ketoreductase domain, accomplishes elongation of the nascent acyl chain by a reduced malonyl, as present in dolaisoleuine. Enzymes encoded in a cassette of genes *aesA-D* likely accomplish the remaining steps in the formation of the AEST backbone. AesA was predicted to incorporate proline, followed by another elongating and ketoreducing PKS (AesB) responsible for incorporation of the reduced malonyl residue of dolaproine. The last amino acid, phenylalanine, was predicted to be incorporated by AesC, and elongated by two additional carbons by the activity of the terminal PKS (AesD). AesD further included a thioesterase domain, which was predicted to catalyze the cleavage of the acyl intermediate from the NRPS/PKS megasynthetase. The remaining biosynthetic step(s) leading to formation of the aetophenine (Aph) residue of AEST were partly obscure. Such modification would likely involve a decarboxylation and reduction step (Fig. 3B); however, these enzymatic activities are not found in AesD. We hypothesize these reactions could be achieved by a putative short-chain dehydrogenase/reductase encoded by a gene (*orf1*) located at the 5’ terminus of the BGC.

**Figure 3.**
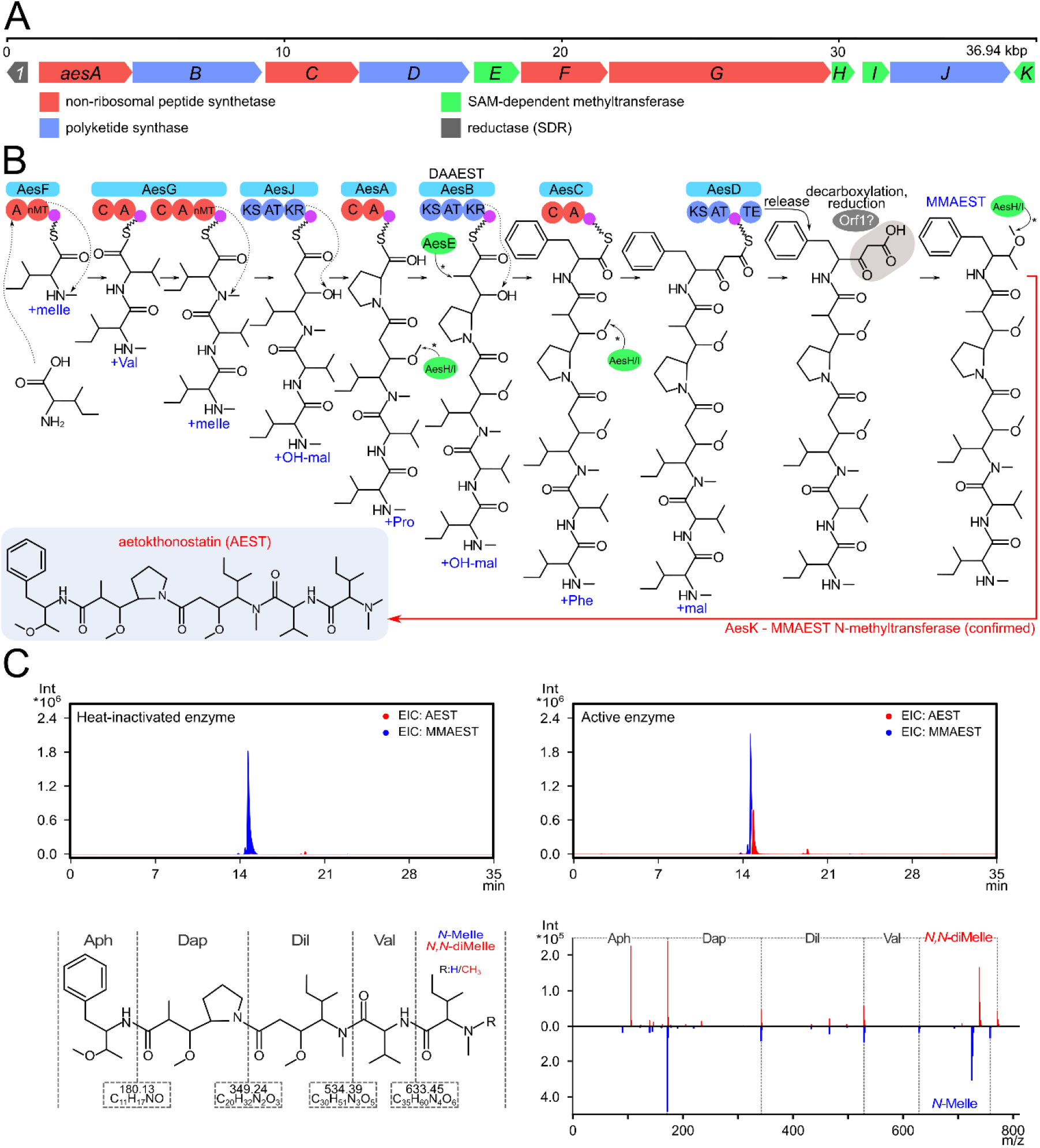
Aetokthonostatins are synthesized via a NRPS/PKS pathway. **(A)** Organization of the AEST biosynthetic gene cluster. **(B)** Scheme of the predicted biosynthetic assembly of AEST. The chronology of the tailoring methylation reactions catalyzed by AesE, H, and I (marked with asterisks) is unknown, the reactions are depicted at their first theoretically suitable substrate. The specific target moieties of AesH/I (the Dil/Dap/Aph OH groups) remain to be assigned experimentally. The proposed decarboxylation/reduction of the C-terminal AEST residue (grey box) was not fully explained by bioinformatic analysis. The N-terminal methylation (red arrow) of monomethylaetokthonostatin (MMAEST) was reconstituted by *in vitro* enzymatic activity of AesK. **(C)** HPLC-HRMS/MS analysis (extracted ion chromatograms) of AEST (red) in reaction mixtures of MMAEST (blue) incubated with heat-inactivated or fresh Strep-AesK (top left/right) show that the inactivated enzyme had no effect on the substrate while the fresh Strep-AesK methylated MMAEST. MMAEST and AEST differ only in the degree of methylation of the *N*-terminal Ile (bottom left). Comparison of the MS/MS spectra of MMAEST and the product of the fresh enzyme reaction (bottom right) shows that AesK methylated the *N*-terminus of MMAEST, producing AEST. Abbreviations: AEST, aetokthonostatin; Aph, aetophenine; Dap, Dolaproine; Dil, Dolaisoleuine; mal, malonyl-CoA; MMAEST, monomethylaetokthonostatin; *N,N*-di-Melle, *N,N*-dimethylisoleucine; OH-mal, hydroxymalonyl-CoA; SAM, *S*-adenosylmethionine; SDR, short-chain dehydrogenase/reductase.

In addition to the NRPS/PKS genes, the BGC contained four genes (*aesE, H, I*, and *K*) annotated as class I S-adenosylmethionine (SAM)-dependent methyltransferases. This observation was in line with multiple methylations found in AEST, specifically the second *N*-methylation of the *N*-terminal isoleucine, *O*-methylation of hydroxyl groups of the dolaisoleuine, dolaproine, and aetophenine residues, and an additional *C*-methyl group at C7 of AEST. As all four methyltransferases were coded by standalone genes, it was impossible to conclusively infer their target atoms in AEST and the precise chronology of their activity in AEST tailoring – the methylation could occur at the nascent acyl chain or *ex post* after cleavage of the intermediate from the NRPS/PKS assembly line. We hypothesized that AesE could more likely catalyze the introduction of C7 methyl of AEST, as its closest functionally annotated hit in MIBiG was the *C-C* methyltranserase GphF (Table S4) from the gephyronic acid biosynthetic pathway (41).

To provide a mechanistic link between the hypothetical *aes* BGC and its putative product, the three methyltransferases AesH, I, and K were heterologously expressed in *E. coli* (Fig. S51) and subsequently used in *in vitro* enzymatic assays to elucidate if any of them is involved in the biosynthesis of the *N,N*-dimethylamino group at the *N*-terminus of AEST. To test this hypothesis, aliquots containing the individual methyltransferases and SAM were incubated with the monomethylated, putative late-stage AEST biosynthetic intermediate **2** as well as with its synthetic analogue MMAF. MMAF was used as an appropriate alternative substrate to test the specificity of the enzymes, as it likewise possesses a single methylation on the *N*-terminal residue, in this case *N*-methylvaline.

We detected no modification of the two substrates in the assays with Strep-AesH and Strep-AesI (Fig. S52-S53). However, in the assay with Strep-AesK, we detected the formation of significant amounts of products at *m/z* 760.05, matching with the mass of AEST, in the assay containing **2** (Fig. 3C), and at *m/z* 746.05 (matching with auristatin F) in the assay containing MMAF (Fig. S54). In both cases, the detected product matched the expected molecular mass corresponding to a single methylation of the provided substrate. HRMS/MS analysis showed that this methylation as expected had occurred at the *N*-terminus of the respective substrates. Indeed, the MS/MS spectrum of the reaction product of **2** was identical to the MS/MS spectrum of AEST, proving that this compound was formed. (Fig. 3C, Fig. S55). We suggest that **2** is the true substrate of AesK, as we found AesK unable to methylate the N-terminal amino acids of **2**, *N*-methylisoleucine, or MMAF, *N*-methylvaline, when provided as monomeric substrates (Fig. S56).

As AesK only methylates the *N*-terminus of both late-stage biosynthesis analogs, we could confirm its predicted role as the last enzyme of the AEST biosynthetic pathway (Fig. 3B, C), demonstrating that the *aes* cluster indeed is responsible for the biosynthesis of AEST. This finding provides additional experimental proof for the cyanobacterial origin of dolastatins (42, 43), offering a cultured and genome-sequenced strain for further laboratory studies. A subsequent search in bioinformatic databases identified highly homologous BGCs in metagenomic assemblies of tropical marine cyanobacteria of the genus *Symploca* (Fig. S50, Tab. S6-S7), from which several dolastatin analogs (e.g. symplostatins) were isolated previously (17, 18, 23, 44). Interestingly, in contrast to the non-colinear arrangement of genes observed in the AEST BGC, the genes in these BGCs from *Symploca* are arranged colinearly with the statin biosynthesis (Fig. S50).

It was intriguing to see that AesK exhibited sufficient substrate specificity to avoid methylation of single *N*-methylisoleucine and *N*-methylvaline, while it was capable to efficiently methylate MMAF, a synthetic dolastatin analog used in antibody-drug conjugate anti-cancer therapy (45). Future in-depth studies of AEST biosynthesis, especially the formation of the unique Aph residue, may broaden our knowledge of the cyanobacterial biosynthetic capabilities and be valuable in synthetic biology and medical biotechnology.

## Materials and Methods

### Summary

All experimental details on the chemical, biological, and computational methods are provided in the Supplementary Information. This includes the isolation and structure elucidation of the natural products, cultivation of the cyanobacteria, information about the cell lines and the cell viability as well as *C. elegans* assays, immunofluorescence microscopy, the preparation of recombinant methyltransferases, the biochemical assays; GNPS Molecular Networking Analysis, genome sequencing and bioinformatic analysis, the docking studies and molecular dynamics simulations. For further details, refer to SI.

### SI

The SI contains a detailed discussion of the structure elucidation as well as of the docking studies and molecular dynamics simulation, all experimental details, and additional tables and figures (including NMR data and spectra of all isolated compounds, MS/MS spectra, crystallography data and refinement statistics). NMR and MS raw data will be available after publication of this manuscript at figshare (DOI 10.6084/m9.figshare.22578616).

## Acknowledgments

We acknowledge A. Dettmer for her support with the cytotoxicity and the *C. elegans* assay and A. Wodak for support with analytical sample preparation. NMR data were recorded by A. Porzel and G. Hahn. We thank R. Ghai for advice on the Oxford Nanopore sequencing and data assembly, and K. Saurav for supporting S.S. while contributing to the study. This work has been funded by the Deutsche Forschungsgemeinschaft (DFG, German Research Foundation – NI 1152/3-1; INST 271/388-1 for T.H.J.N.), the Czech Science Foundation (GAČR – 19-21649J for J.M.), and the Grant Agency of the University of South Bohemia (GAJU 112/2022/P for J.A.M.Y.).

