## Supplementary Information for "More than just an Eagle Killer: The freshwater cyanobacterium *Aetokthonos hydrillicola* produces highly toxic dolastatin derivatives"

### Table of Contents

| | Page |
|---|---|
| **SUPPLEMENTARY TEXT** | |
| Discussion of the NMR based structure elucidation of AEST | 2 |
| Docking study and Molecular dynamics simulation | 3 |
| **METHODS – Natural Product Chemistry** | 5 |
| **METHODS - Biology** | 7 |
| **METHODS - Computational and Bioinformatics** | 9 |
| **SUPPLEMENTARY REFERENCES** | 11 |
| **SUPPLEMENTARY TABLES** | 13 |
| **SUPPLEMENTARY FIGURES** | |
| Bioactivity characterization | 20 |
| Structure elucidation | 22 |
| Docking study and Molecular Dynamics simulation | 48 |
| Biosynthesis | 53 |
| In vitro enzyme assays | 55 |

### SUPPLEMENTARY TEXT

**Discussion of the NMR-based structure elucidation of aetokthonostatin (1).** NMR analysis (Fig. S4-S9) was complicated by a considerable signal overlap in the $^{1}H$ NMR spectrum, caused by the presence of two conformers in solution (ratio main to minor conformer about 3:1, Fig. S4). Analysis of the $^{13}C$ and $^{1}H$ NMR data suggested four amide or ester carbonyls (173.48, 168.84, 172.89, and 169.78 ppm) and two NH protons (7.90 and 8.02 ppm; NMR data see Tab. S1). The presence of amide bonds was confirmed by four signals agreeing with α-carbonyl protons of amino acids (2.25, 2.35, 4.64 and 2.79 ppm), giving us confidence that **1** was a peptide. Furthermore, we could observe signals of six N- or O-methyl groups. Evaluation of HSQC-DEPT, TOCSY and COSY spectra allowed us to assemble the spin systems of the amino acids in the peptide (Fig. S0). The amino acid sequence and the positions of the N- and O-methyl groups could be deduced from a HMBC spectrum. We could identify isoleucine and valine as the two N-terminal amino acids in the molecule. The third amino acid had a side chain like isoleucine, but featured three additional proton signals in its spin system (3.98 and 2.35/2.17 ppm), attached to two carbons. Evaluation of the HMBC and COSY spectra, we observed correlations indicating a $CH_2$ (2.35/2.17 ppm) group next to the carbonyl carbon (168.84 ppm), and an O-methyl group attached to a CH (3.20 ppm) that was connected to the carbon bearing the isoleucine side chain. This amino acid was thus identified as dolaisoleuine, indicating that our compound might belong to the dolastatin compound class. With this knowledge, the NMR data of the fourth amino acid could easily be assigned, as they fit to dolaproine, another amino acid characteristic for dolastatins. However, the NMR-data of the C-terminus did not fit to any known C-terminus in the dolastatin compound family. Analysis of the $^{1}H$, HMBC and HSQC spectra revealed the presence of an aromatic phenyl (7.12 – 7.20 ppm) and an O-methyl (3.32 ppm) group. With the help of HMBC correlations from CH (3.38 ppm) to the phenyl group and the carbonyl carbon, we were able to deduce the structure of a so far unknown monomer at the C-terminus, that we called aetophenine (Aph). ROESY data confirmed the monomer sequence of the compound as *N,N*-dimethyl-Val-Ile-Dil-Dap-Aph. This amino acid sequence was supported by the observed MS/MS fragmentation pattern (Figure S10). We named the main compound produced by *A. hydrillicola* "aetokthonostatin". Structure elucidation of **2** and **3** were conducted accordingly (NMR spectra Fig. S11-S16 and S18-S21; MS/MS spectra Fig. S17 and S22).

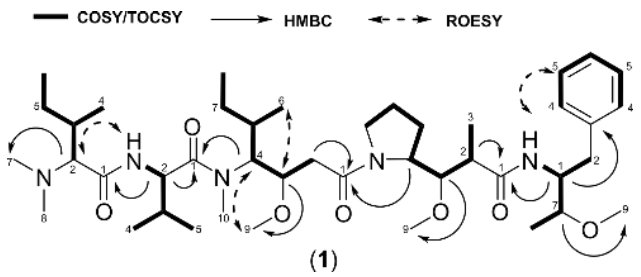


**Fig. S1.** COSY (bold bonds) and key HMBC correlations (arrows) of aetokthonostatin (1).

**Docking study.** Superposition of the available crystal structures revealed that the binding site residues adopt a similar conformation in all complexes. This observation and the high structural similarity of the cocrystallized inhibitors to MMAF and AEST led to the selection of 4X1K as PDB structure for the docking of both inhibitors. Docking protocol validation was carried out within the framework of a re- and cross-docking study: the binding modes of three inhibitors (4X1K, 4X20 and 4X1Y) could be reproduced using the selected protein structure (RMSD 0.321 Å, 0.869 Å, 0.579 Å, respectively), and only docking of inhibitor bound in the PDB structure 4X1I resulted in higher RMSD values (3.17 Å). Visualization of the docking poses of the latter compound revealed that the ligand‘s N-terminal and core parts were correctly docked. However, the aromatic groups at the C-terminal part of the peptide showed an inverted orientation as in the reference structure (Fig. S43).
Analyzing the available crystal structures of tubulin with different peptidic inhibitors revealed that all cocrystallized ligands show comparable ligand-receptor-interactions: the peptide backbone forms hydrogen bonds to the backbone of Tyr β224 and Gly β225. In addition, a bifocal hydrogen bond network with the side chain of Asn α329 can be observed (Fig. S44). The inhibitors charged N-terminal part shows electrostatic interactions with Asp β179. Except structure 4X1I, all inhibitors form salt-bridges with the side chain of Arg β278 involving the C-terminal carboxylic acid group. In the inhibitor structure found in 4X1I, the carboxylic acid group is replaced by an aromatic thiazole ring (1). Interestingly, in the resolved crystal structure, the benzyl moiety was observed to occupy the same position as the carboxylic acid group in 4X1K, 4X1Y or 4X20, and to undergo cation-$\pi$ interactions with Arg β278 (Fig. S. 44, panel A).
Compound MMAF has a structure similar to most of the cocrystallized ligands featuring a C-terminal carboxylic acid and an N-terminal amine moiety. As expected, the generated docking pose shows similar ligand-receptor-interactions. This is almost the same case for compound AEST (Fig. S44), although it features the more hydrophobic aetophenine as C-terminal group. Since the absolute sterochemical configuration of monomer Aph in AEST hast not yet been defined, all four possible isomers were included to the docking study. The top-ranked docking poses of the SR- and SS-configured peptides show a buried orientation of the C-terminal benzyl residue where the solvent-exposure of the hydrophobic group is minimized (Fig. S44 panels C, D). The remaining isomers lead to top-scored docking poses in which the benzyl residue is oriented towards Arg β278 (Fig. S44, panels E, F). Based on the docking scores, all four stereoisomers appear to bind similarly strong to tubulin. For MMAF and AEST, the same hydrogen bonds of the peptidic backbone were observed as in the X-ray structures of the cocrystallized inhibitors. Also, the ionic interaction between the N-terminal protonated amine and Asp179 is observed for all inhibitors.

**Molecular dynamics simulations.** Since the binding poses for MMAF and AEST were generated while applying a rather constrained docking protocol, we decided to confirm the binding pose stability by performing a series of MD simulations as described in the methods section.
At first, a MD simulation run was performed with crystal structure 4X1K, including the corresponding cocrystallized inhibitor. During the simulation, the protein remained stable, and no major deviations were observed for the inhibitor from its original pose. However, some peaks are visible in the ligand’s RMSD plot (Fig. S45, panel A). An analysis of the ligand’s RMSF values (Fig. S46, panel A) revealed that the C-terminal benzyl moiety is significantly moving during the simulations. This observation is in accordance with the given B-factor values, though, and might also be the reason for the aforementioned increased RMSD values regarding the crossdocking poses of ligand 4X1I.

Furthermore, MD simulations were performed for the docked ligands MMAF and AEST. The ligand RMSD values for MMAF and most of the AEST isomers remain stable and mostly below 3.5 Å during the MD simulation, confirming the accuracy of the generated docking poses (Fig. S45). In case of the four stereoisomers of AEST, the binding of the ligand was stable with higher flexibility of the C-terminal Aph residue. This is due to the solvent exposure of the Aph residue, which does not directly interact with the binding pocket (no hydrogen bond or salt bridges). For the stereoisomer of AEST with the R,R configuration of Aph resulted in larger deviation of the C-terminal residue (Fig. S45, panel F). It is noteworthy that again most fluctuations occur within the peptide's C-terminus (Fig. S46, S47), especially the carboxylic acid and benzyl moiety are affected.

**METHODS – NATURAL PRODUCT CHEMISTRY**

**Natural products analysis.** 1D and 2D NMR data for AEST (**1**) and its derivatives (**2**, **3**) were recorded on an Agilent Varian VNMRS 600 MHz spectrometer, operating at 600 MHz ($^{1}H$) and 150 ($^{13}C$), using DMSO-$d_6$ as solvent (referenced to $\delta_H$ 2.51, $\delta_C$ 39.54 for residual solvent). The instrument was equipped with a 5 mm cryoprobe. HRESIMS[2] data were acquired on a Q Exactive Plus mass spectrometer (Thermo Fisher Scientific) equipped with a heated ESI interface coupled to an UltiMate 3000 HPLC system (Thermo Fisher Scientific). The following parameters were used for the data acquisition: pos. mode, ESI spray voltage 4.5 kV; neg. mode, ESI spray voltage 2.5 kV, scan range 150 – 2000 m/z. Analytical and semipreparative HPLC were performed on an UltiMate 3000 HPLC system (Thermo Fisher Scientific). Flash chromatography was carried out on a GX-271 Liquid Handler system equipped with a 322 series pump and a 171 series diode array detector (Gilson).

**Extraction and Isolation.** Lyophilized biomass was mechanically homogenized with a spatula, resuspended in a methanol/$H_2O$ mixture 50/50 % (v/v, 1 mL per 10 mg biomass) and subsequently treated with an ultra sonotrode (Bandelin Sonificator 250, 5 min, 100 % duty cycle) to break up cell walls. To prevent degradation, the samples were maintained on ice during sonification. Samples were placed on an overhead shaker for 0.5 h, centrifuged (5100 rpm for 5 min), and the supernatant was collected. The samples were extracted with the same method, but the polarity of the solvent was decreased using a mixture of methanol/$H_2O$ 80/20 % (v/v) in the second step and 100 % methanol in the final third extraction step. Supernatants were collected and combined after each extraction step. Drying under reduced pressure resulted in a green, viscous extract. The extract was redissolved in a mixture of methanol/$H_2O$ 80/20% (v/v) and filtered (45 µM) to remove undissolvable residue. Celite (Celite® 545, Carl Roth, Germany) was added, and the solvent was removed under reduced pressure by rotary evaporation for dry loading flash chromatography, using an ACN-$H_2O$ gradient (5-100 % in 25 min followed by 100 % ACN for 5min). Fractions were collected every 1.5 min. Fractions were subsequently tested in the sulforhodamine B assay to confirm which fractions contained the cytotoxic compound of interest. Fractions 12-15 were confirmed as the fractions of interest and submitted to semipreparative reversed-phase HPLC (Phenomenex Luna C18 5µM, 250 x 10mm, 6.3 mL/min, UV detection at 210 nm), using an ACN-H2O (10 mM ammonium acetate, pH 9) linear gradient (60-85 % in 15 min) to afford compounds **1**, **2**, and **3**.

**Aetokthonostatin (AEST, 1).** Colorless, amorphous solid; $^{1}H$ and $^{13}C$ NMR data see Table S1; HRESI *m/z* $[M + H]^+$ 774.5729 (calculated for $C_{43}H_{76}O_7N_5$, 774.5739). Isolation yield: 0.15 % of dry biomass.

**Monomethyl-AEST (MMAEST, 2).** Colorless amorphous solid; $^{1}H$ and $^{13}C$ NMR data see Table S2; HRESI *m/z* $[M + H]^+$ 760.5571 (calculated for $C_{42}H_{74}O_7N_5$, 760.5569). Isolation yield: 0.03 % of dry biomass.

**Des-Aph-AEST (DAAEST, 3).** Colorless amorphous solid; $^{1}H$ and $^{13}C$ NMR data see Table S3; HRESI *m/z* $[M + H]^+$ 613.4520 (calculated for $C_{32}H_{61}O_7N_4$, 613.4535). Isolation yield: 0.01 % of dry biomass.

**HPLC microfractionation of an *A. hydrillicola* extract.** Microfractionation was carried out on a Luna® column (C18, 5 µm, 4.6 x 250 mm) using a gradient from 5-100 % (v/v) of ACN in water (0.1 % formic acid each) in 30 min with a flowrate of 1.3 mL/min. Initial mobile phase conditions were maintained for 5 min before starting the gradient and the final plateau was maintained 5 min before finishing the run. Fifty-three fractions were collected automatically, dried and resuspended in 10 % (v/v) DMSO in water for cytotoxicity testing.

**Determination of the absolute configuration of Dap**. The absolute configuration of Dolaproine was established using Marfey's method as described in (2). **1** and MMAE (500 µg each) were hydrolyzed in 6 N HCl (1 mL) in a sealed reaction vessel at 110 °C for 24 h. After drying under a stream of nitrogen, FDAA solution [(1-fluoro-2,4- dinitrophenyl)-5-L-alanine amide] in acetone (50 µL, 0.04 M) and NaHCO3 (100 µL, 1M) were added to each reaction vessel. The solution was kept at 80 °C for 3 min, cooled to room temperature and acidified with HCl (2 M, 50 µL). After the addition of ACN/water (200 µL, 1:1), the FDAA-amino acid derivatives from the hydrolysates were analyzed and compared by HPLC-MS analysis: Phenomenex Kinetex C18, 2.6 µm; 100 x 3 mm, flow rate 0.4 mL/min, 50 °C, gradient of 5 % (ACN + 0.1 % FA) to 100 % (ACN + 0.1 % FA) within 14 min. The absolute configuration was determined by comparing the retention time of the derivatized cis/trans dolaproine monomer from MMAE (rt 6.89 min and 5.64 min) with the retention time of the derivatized cis/trans dolaproine from AEST (rt 6.90 and 5.64 min min).

### METHODS – BIOLOGY

**Cyanobacteria Cultivation.** *Aetokthonos hydrillicola* was cultivated in 1 L glass bottles or in 20 L polycarbonate carboys in standard BG11 medium (3). Cultures were gently stirred using a magnetic stirrer (~100 rpm) and supplemented with 5 % CO2 (1 L h-1). Fluorescent tubes (Sylvania GroLux, F18W/GRO) were used as light source providing an average light intensity of 35 $\mu Em^{-2}s^{-1}$. Cultivation temperature was 28 °C.

**Cell Lines and Reagents.** Hela cells were purchased from the German Collection of Microorganisms and Cell Cultures GmbH (DSMZ). Cells were cultured in Dulbecco's modified Eagle medium low glucose (ROTI® CELL DMEM Low Glucose, Carl Roth, Germany) supplemented with 10% fetal bovine serum (FBS, Sigma, Germany) and 2 mM glutamine solution (ROTI®-CELL Glutamin-Lösung, Carl Roth, Germany) at 37 °C in a humidified incubator with 5 % $CO_2$. Sulforhodamine B was purchased from Sigma Aldrich, dissolved in 1% (v/v) acetic acid and stored at 8°C for further use.

**Sulforhodamine B Assay.** A Sulforhodamine B (SRB) assay was used to evaluate the cytotoxic potency of the extract, fractions, and pure compounds in HeLa cell line (4) . All experiments were run in triplicates and with three replications, except for the assay with DAAEST (**2**), which was run two times in triplicates. Cells were plated at a density of 10,000 cells per well in 96-well plates and allowed to adhere overnight. The cells were subsequently treated with the indicated concentrations for 48 h after which the cells were fixed (10% TCA in $H_2O$) and stained with SRB (0.057 % solution in 1 % (v/v) acetic acid). Cell viability was measured by the absorbance at 510 nm using a Tecan plate reader (INFINITE M PLEX, Switzerland). $IC_{50}$ was determined by non-linear regression analysis using GraphPad Prism 6.

**Immunofluorescence microscopy.** HeLa and MDA-MB231 cells were grown in 6-well plates on coated coverslips and treated with the compound of interest with 600 µL of exposure solution. The following exposure concentrations of MMAF and AEST were selected based on their potency against the cell lines: HeLa (100 nM, 1 nM, 0.1 nM) and MDA-MB 231 (100 and 10 nM). As a positive control, 1 µM taxol was used. The procedure of fixation and staining was as follows: Cells were washed twice with pre-warmed PBS, fixed with 4 % formaldehyde in PBS for 15 min at room temperature (RT), washed two times with PBS, permeabilized with 0.25 % Triton X-100 in PBS for 5 min, and again washed two times with PBS. Consequently, the cells were left in 1 % bovine serum albumin for 1 h at RT, incubated with alpha-tubulin mouse monoclonal antibody (A11126-Invitrogen™) at a concentration of 1ug/ml for 3 hours at RT, washed two times with PBS, and 5 µl Nuc Blue™ Fixed Cell Stain solution (Invitrogen™) per well for DAPI signal. Finally, 1 µL of 2mg/mL stock solution of Alexa Fluor 488 Rabbit Anti-Mouse IgG Secondary Antibody (A11001-Invitrogen™) was added to 200 µL in well and incubated for 30 min in darkness at RT. The coverslip with cells was attached to a slide using mounting gel ProLong™ Glass Antifade Mountant (Invitrogen™) and observed the next day. Images were acquired by a laser scanning confocal microscope (Zeiss LSM 880; Carl Zeiss Microscopy GmbH) equipped with a Plan-Apochromatic 20×/0.8 M27 objective. Cells were excited with 405 nm and 488 nm laser for Nuc Blue™ staining and for tubulin staining Alexa Fluor 488, respectively. Signal was detected at wavelengths 419 - 544 nm for NucBlue Dapi staining and 499 - 544 nm for tubulin staining Alexa fluor 488 in a frame mode. The signal was detected by a GaAsP photomultiplier in 8-bit mode and visualised by Software Zen Blue 3.3.

***C. elegans* – Cultivation.** *C. elegans* were cultured at 22 °C in 90 mm petri dishes on NGM agar seeded with *E. coli* OP50 as a food source. Synchronous populations of the worms were obtained by treating gravid worms for 5 min with 5M NaOH and 5 % NaOCl for lysis and decontamination. The lysate was pelleted by centrifugation (1200 rpm, 2 min). The eggs were separated from the pellet by density gradient centrifugation (300 rpm, 1 min) using a sucrose solution (60 % w/v) and water in the ratio 1:1. The upper layer, in which the eggs float, was collected and transferred to a fresh tube with sterile water (1:3) and then submitted to a last centrifugation step (350 rpm, 5 min) to pellet the eggs and wash out the sucrose. The eggs were allowed to hatch in M9 buffer.

***C. elegans* – Reproduction assay.** A 24-well plate was prepared with NGM agar and 20 µL of an *E. coli* OP50 solution was pipetted in every well. The plate was incubated at 37 °C for 24 h and subsequently treated with either 10 % DMSO (negative control) or different concentrations of AEST in DMSO. After adding the test compound solutions, the plate was incubated for another hour. Three age synchronized L4 larvae were placed in every well. After 72 h, the total number of healthy adult worms were counted. $IC_{50}$ was determined by non-linear regression analysis using GraphPad Prism 6.

**Preparation of recombinant methyltransferases.** The *aesH*, *aesI* and *aesK* genes, all predicted to code for the three SAM-dependent methyltransferases at the 5´ end of the AEST BGC, were codon-optimized (Fig. S46) for expression in *E. coli* and synthesized by GenScript (USA). The obtained synthetic genes were sub-cloned using *NdeI* and *NheI* restriction sites into the in-house pET-28a-derived vector containing an N-terminal StrepII-tag and improved translation initiation region and T7 promoter according to Shilling et al. (5). Plasmids were transformed into the electro-competent *E. coli* BL21(DE3) cells. To produce the recombinant methyltransferases, *E. coli* cells were grown in 300 mL of an auto-inducible LB broth medium (FORMEDIUM, United Kingdom) supplemented with 50 µg/mL kanamycin in an orbital shaker at 200 rpm and 37 °C. After reaching an OD at 600 nm of ~0.5, the temperature was decreased to 18 °C and, under these conditions, the cells were incubated for additional 18 h. The cell cultures were harvested by centrifugation (10 min, 4 °C, 4,000 g), re-suspended in a lysis buffer containing 100 mM Tris-HCl (pH = 8.0), 150 mM NaCl, 5 mM $MgCl_2$, 1 mM EDTA, 1 mM DTT, 5 µg/mL DNase I, 1 mg/mL lysozyme and complete protease inhibitor cocktail (Roche). After incubation at 10 °C for 45 min, the cells were disrupted by sonication, and the obtained lysate was clarified by centrifugation (12,000 rpm at 8 °C for 30 min). The soluble strep-tagged proteins were purified by affinity chromatography using the Strep-Tactin Superflow high-capacity resin (IBA Lifesciences, Germany) following the manufacturer's instructions, and the purity of the proteins was confirmed by SDS-PAGE gel (Fig. S48).

**SAM methyltransferase *in vitro* assays.** The methyltransferase activity assay was conducted as follows: 100 µM of the substrate and 3 µM of the enzyme were incubated in 100 µL of the assay buffer containing 100 mM Tris-HCl, 150 mM NaCl, 5 mM $MgCl_2$, 1 mM EDTA, 1 mM DTT and 1 mM SAM (Sigma-Aldrich). The reaction tubes were gently rotated at 28 °C for 3 h and then the reaction was quenched with an equal volume of methanol. The supernatant, obtained by centrifugation at 12,000 g for 10 min, was analyzed by HPLC-MS. All assays were conducted in triplicates. As negative controls, we assayed the equivalent concentration of the analyzed enzyme denatured at 100 °C for 10 min and additionally an enzyme-less reaction mixture.

### METHODS – COMPUTIONAL METHODS AND BIOINFORMATICS

**GNPS Molecular Networking Analysis.** The raw data was converted to the .mzXML format using the software MSConvert (version 3.0.21026-c1c9c6b09) (6). The converted data was processed with the software MZmine (version 2.53) following the online workflow (https://ccms-ucsd.github.io/GNPSDocumentation/featurebasedmolecularnetworking-with-mzmine2/) (7–11). Data analysis used default parameters, except for the noise level (1.E04), group intensity threshold (1.E05), min. highest intensity (1.E05), *m/z* tolerance (0.005 *m/z*), RT range for MS2 scan pairing (0.2 min). The mass spectral network was assembled and visualized using Cytoscape (version 3.8.2) (12).

**Genome sequencing and bioinformatic analysis.** The previously published genome assembly of *A. hydrillicola* Thurmond2011 was improved in terms of completeness and contiguity using additional sequencing and genome assembly (13). Two types of DNA templates were prepared: (i) total genomic DNA isolated from the non-axenic uni-cyanobacterial strain using NucleoSpin Soil Kit (Macherey-Nagel), (ii) a pooled MDA product obtained by whole genome amplification from separated single filaments of *A. hydrillicola* as described previously (13). Each of the two DNA templates was sequenced using *de novo* Illumina MiSeq (Illumina, San Diego, CA, USA) with a 400 bp average length of insert pair-end library and 250 bp reads (sequencing yield of ~2.5 Gbp) by SeqMe (Dobříš, Czech Republic). In parallel, each of the templates was sequenced using Oxford Nanopore MinION device with Rapid Sequencing kit and the R9 version FlowCell (Oxford Nanopore) in our laboratory (sequencing yield of ~6.3 Gbp) to obtain longer sequencing reads and improve the genome draft contiguity. Raw sequencing reads in each of the Illumina data sets were trimmed and assembled separately using BBTools and SPAdes 3.11 with the single-cell option enabled (14). The resulting assemblies were automatically annotated using Prokka v 1.12 (15), and the deduced CDS were subjected to BLASTp search against prokaryotic RefSeq genomes. All scaffolds retrieving top BLAST hits from cyanobacteria shared a similar GC content and thus were further assumed to be part of the *A. hydrillicola* genome. The previously available genomic scaffolds of *A. hydrillicola* Thurmond2011 (GenBank accession JAALHA010000000) and the scaffolds originating from Illumina sequencing in this study were pooled into a single FASTA file. Subsequently, all raw sequencing reads ever obtained for the strain by Illumina and Oxford Nanopore sequencing reads (in FASTQ format) were mapped to the pool of genomic scaffolds created in the previous step, using the Map to Reference tool in Geneious Prime v. 2020.1.2 (https://www.geneious.com). The successfully mapped reads were saved to a new file. The resulting sets of mapped Illumina and Nanopore reads were then co-assembled *de-novo* using SPAdes to accomplish the final genome assembly published in this study (GenBank accession JAALHA020000000). As it was assumed that AEST could be synthesized by NRPS/PKS enzymatic machinery, the genome assembly was analyzed using antiSMASH v 6.0 (16). Selected gene clusters were investigated in detail using BLASTp against the nr database, and searched in the MIBiG database (17). Substrate specificity of individual amino acid adenylation domains encoded in NRPS genes was predicted using the antiSMASH v. 6.0 pipeline.

**Docking study.** To evaluate the accuracy of molecular docking for tubulin, four crystal structures of tubulin cocrystallized with inhibitors were considered (PDB IDs 4X1I, 4X1K, 4X1Y, 4X20, structures of cocrystallized compounds see Fig. S41). 3D structures were downloaded from the Protein Data Bank (PDB, http://www.rcsb.org). Metal ions present in the structures as well as protein chains not being involved in forming the binding pocket (chains A, D and E) were deleted for each structure. Subsequently, Schrödinger's Protein Preparation Wizard was used to prepare the protein structures for ligand docking by adding hydrogen atoms, filling in missing side chains and loops, capping the chains' termini and optimizing the hydrogen bond network (at pH 7.4) (18, 19). Finally, an energy minimization step was executed using the default settings. The receptor grid was generated by assigning the cocrystallized ligand as the centre of the grid. The four inhibitor structures were extracted and prepared while retaining specified chiral centres and keeping charges using Schrödinger's Ligprep (20). Confgen was used afterwards to generate 10 diverse conformers per ligand (21, 22). Docking of the highly flexible AEST was performed using Schrödinger's Glide in Standard Precision-Peptide (SP-Peptide) mode (23–26). In re- and cross-docking trials, the highest accuracy was observed when using a docking protocol in Glide including a core constraint. In this mode, a PDB crystal structure (here PDB ID 4X1K) is used as a template restricting the docking poses to the position of a reference ligand's core defined via a consensus pattern of the backbone peptide groups. While restricting the docking procedure to defined reference position, the pre-set tolerance value of 0.1 Å was kept. Additionally, the option to retry docking with less tight core constraints (1.0 Å) in case poses are rejected was enabled. Input ring conformations were also considered during the docking process. Resulting binding poses were visualized using PyMOL (27).

**Molecular dynamics simulations.** MD simulations were carried out for obtained protein-docking pose complexes using the Amber18 software (28). AM1-BCC atomic charges were used for ligands (29, 30). The protein chains were prepared for tLeap input using pdb4amber. Afterwards, tLeap executed parameterization according to the ff14SB force field for protein residues and the General Amber Force Field 2 (GAFF2) for ligand structures (31–33). The resulting system was solvated in an octahedral periodic box of TIP3P water molecules at a margin of 10 Å and neutralized using $Na^+$ ions (34). Prior to the production step, the built system underwent two minimization steps, a heating and a pressure equilibration step. The first minimization step included 1000 iterations of steepest descent and 2000 iterations of conjugate gradient minimization affecting only solvent molecules. In the second minimization step, the whole system underwent 2000 iterations of steepest descent and 2000 iterations of conjugate gradient minimization. Afterwards, the system was subsequently heated to production temperature (300 K) through 100 ps of MD simulation, while keeping protein chains and ligands restrained again with a force constant of 10 $kcal*mol^{-1}*Å^{-2}$. Constant volume periodic boundary was set to equilibrate the temperature of the system by Langevin thermostat using a collision frequency of 2 $ps^{-1}$. Final pressure equilibration of the system was performed for 100 ps at the target temperature (300 K) while applying a constant pressure of 1 bar. The actual production run of 50 ns with a time step of 2 fs was simulated at a constant temperature of 300 K using Langevin thermostat with a collision frequency of 2 $ps^{-1}$. Two individual MD simulations (each 50 ns) were performed for each protein-inhibitor complex (MD1 and MD2). A non-bonded cut-off distance of 10.0 Å for long-range electrostatic interactions was used by applying the Particle Mesh Ewald (PME) method during the temperature equilibration and MD routines (35). The SHAKE algorithm was applied to constrain all bonds involving hydrogen (36). Marvin Sketch was used for 2D structure depiction and atom number assignment (37).

### SUPPLEMENTARY TABLES

**Table S1.** NMR spectroscopic data of aetokthonostatin (**1**; major and minor conformer). All spectra were recorded in DMSO-$d_6$ ($^1$H 600 MHz, $^{13}$C 150 MHz). n.o. not observed.

| | | conformer 1 | | conformer 2 | |
|---|---|---|---|---|---|
| unit | C/H no. | $\delta_H$ ($J$ in Hz) | $\delta_C$, mult | $\delta_H$ ($J$ in Hz) | $\delta_C$, mult |
| Aph | 1 | 4.23, m | 51.5, CH | 4.13, m | 52.5, CH |
| | 2a | 2.84, m | 35.0, $CH_2$ | 2.80, m | 34.8, $CH_2$ |
| | 2b | 2.69, m | | 2.66, m | |
| | 3 | | 139.0, qC | | 139.2, qC |
| | 4a/b | 7.20, m | 128.6, CH | 7.19, m | 128.8, CH |
| | 5a/b | 7.19, m | 127.9, CH | 7.19, m | 128.0, CH |
| | 6 | 7.12, m | 125.8, CH | 7.11, m | 125.7, CH |
| | 7 | 3.38, m | 77.1, CH | 3.32, m | 76.9, CH |
| | 8 | 1.07, m | 14.9, $CH_3$ | 1.07, m | 14.6, $CH_3$ |
| | 9 | 3.32, s | 56.0, $CH_3$ | 3.30, s | 55.9, $CH_3$ |
| | NH | 7.90, d (9.13) | | 7.64, d (8.84) | |
| Dap | 1 | | 173.4, qC | | 173.0, qC |
| | 2 | 2.25, m | 42.9, CH | 2.21, m | 43.5, CH |
| | 3 | 1.07, m | 15.9, $CH_3$ | 1.06, m | 15.9, $CH_3$ |
| | 4 | 3.30, m | 85.3, CH | 3.79, m | 81.0, CH |
| | 5 | 3.15, m | 58.1, CH | n.o | |
| | 6a | 1.69, m | 25.1, $CH_2$ | 1.66, m | 25.2, $CH_2$ |
| | 6b | 1.22, m | | 1.28, m | |
| | 7a | 1.79, m | 24.2, $CH_2$ | 1.72, m | 23.1, $CH_2$ |
| | 7b | 1.48, m | | 1.45, m | |
| | 8a | 3.54, m | 46.1, $CH_2$ | 3.45, m | 47.1, $CH_2$ |
| | 8b | 3.02, m | | 3.20, m | |
| | 9 (O-Me) | 3.24, s | 60.7, $CH_3$ | 3.21, s | 60.0, $CH_3$ |
| Dil | 1 | | 168.8, qC | | 168.6, qC |
| | 2a | 2.35, m | 34.8, $CH_2$ | 2.43, m | 37.1, $CH_2$ |
| | 2b | 2.17, m | | 2.25, m | |
| | 3 | 3.98, m | 77.0, CH | n.o | |
| | 4 | 4.73, m | 54.7, CH | 4.64, m | 55.5, CH |
| | 5 | 1.83, m | 31.3, CH | 1.76, m | 32.3, CH |
| | 6 | 0.87, m | 15.1, $CH_3$ | 0.89, m | 15.3, $CH_3$ |
| | 7a | 1.30, m | 25.0, $CH_2$ | n.o | |
| | 7b | 0.91, m | | n.o | |
| | 8 | 0.76, m | 10.2, $CH_3$ | n.o | |
| | 9 (O-Me) | 3.20, s | 56.9, $CH_3$ | 3.17, s | 56.8, $CH_3$ |
| | 10 (N-Me) | 3.09, s | 31.0, $CH_3$ | 3.00, s | 31.3, $CH_3$ |
| Val | 1 | | 172.9, qC | | 173.0, qC |
| | 2 | 4.64, m | 53.5, CH | 4.54, m | 53.7, CH |
| | 3 | 2.01, m | 29.6, CH | 1.95, m | 29.6, CH |
| | 4 | 0.96, m | 18.8, $CH_3$ | 0.92, m | 18.7, $CH_3$ |
| | 5 | 0.96, m | 18.8, $CH_3$ | 0.92, m | 18.7, $CH_3$ |
| | NH | 8.02, br | | 8.03, br | |
| N-diMe-Ile | 1 | | 169.7, qC | | 169.8, qC |
| | 2 | 2.79, m | 70.4, CH | n.o | |
| | 3 | 1.76, m | 32.2, CH | n.o | |
| | 4 | 0.71, m | 15.1, CH3 | n.o | |
| | 5a | 1.57, m | 24.6, $CH_2$ | n.o | |
| | 5b | 1.08, m | | n.o | |
| | 6 | 0.83, m | 10.2, $CH_3$ | n.o | |
| | 7 (N-Me) | 2.21, m | 41.6, $CH_3$ | n.o | |
| | 8 (N-Me) | 2.21, m | 41.6, $CH_3$ | n.o | |

**Table. S2.** NMR spectroscopic data of monomethyl-aetokthonostatin (**2**; major and minor conformer). All spectra were recorded in DMSO-$d_6$ ($^1$H 600 MHz, $^{13}$C 150 MHz). n.o. not observed.

| | | conformer 1 | | conformer 2 | |
|---|---|---|---|---|---|
| unit | C/H no. | $\delta_H$ ($J$ in Hz) | $\delta_C$, mult | $\delta_H$ ($J$ in Hz) | $\delta_C$, mult |
| Aph | 1 | 4.22, m | 51.5, CH | 4.11, m | 52.3, CH |
| | 2a | 2.84, m | 35.0, $CH_2$ | 2.80, m | 34.7, $CH_2$ |
| | 2b | 2.70, m | | 2.65, m | |
| | 3 | | 139.0, qC | | 139.3 qC |
| | 4a/b | 7.20, m | 128.6, CH | 7.19, m | 128.9, CH |
| | 5a/b | 7.19, m | 127.8, CH | 7.20, m | 128.0, CH |
| | 6 | 7.11, m | 125.8, CH | 7.11, m | 125.7, CH |
| | 7 | 3.37, m | 77.5, CH | 3.32, m | 76.9, CH |
| | 8 | 1.06, m | 15.0, $CH_3$ | 1.06, m | 14.7, $CH_3$ |
| | 9 (O-Me) | 3.32, s | 56.0, $CH_3$ | 3.30, s | 56.0, $CH_3$ |
| | NH | 7.88, d (9.23) | | 7.64, d (8.78) | |
| Dap | 1 | | 173.6, qC | | 173.2, qC |
| | 2 | 2.24, m | 43.0, CH | 2.22, m | 43.6, CH |
| | 3 | 1.06, m | 15.9, CH | 1.06, m | 15.5, $CH_3$ |
| | 4 | 3.30, m | 85.3, CH | 3.79, m | 81.3, CH |
| | 5 | 3.15, m | 58.2, CH | 3.20, m | 55.0, CH |
| | 6a | 1.68, m | 25.1, $CH_2$ | 1.66, m | 25.0, $CH_2$ |
| | 6b | 1.22, m | | 1.28, m | |
| | 7a | 1.78, m | 24.0, $CH_2$ | 1.72, m | 22.9, $CH_2$ |
| | 7b | 1.48, m | | 1.41, m | |
| | 8a | 3.53, m | 46.2, $CH_2$ | 3.44, m | 47.2, $CH_2$ |
| | 8b | 3.02, m | | 3.20, m | |
| | 9 (O-Me) | 3.23, s | 60.9, $CH_3$ | 3.20, s | 60.3, $CH_3$ |
| Dil | 1 | | 168.9, qC | | 168.6, qC |
| | 2a | 2.35, m | 34.8, $CH_2$ | 2.43, m | 36.9, $CH_2$ |
| | 2b | 2.17, m | | 2.25, m | |
| | 3 | 3.98, m | 77.0, CH | n.o | |
| | 4 | 4.73, m | 54.7, CH | 4.62, m | 55.5, CH |
| | 5 | 1.83, m | 31.3, CH | 1.77, m | 31.8, CH |
| | 6 | 0.86, m | 15.2, $CH_3$ | 0.88, m | 15.3, $CH_3$ |
| | 7a | 1.29, m | 25.1, $CH_2$ | n.o | |
| | 7b | 0.91, m | | n.o | |
| | 8 | 0.77, m | 10.0, $CH_3$ | n.o | |
| | 9 (O-Me) | 3.20, s | 57.0, $CH_3$ | 3.17, s | 56.8, $CH_3$ |
| | 10 (N-Me) | 3.05, s | 31.1, $CH_3$ | 2.97, s | 31.3, $CH_3$ |
| Val | 1 | | 172.6, qC | | 172.8, qC |
| | 2 | 4.67, m | 53.3, CH | 4.54, m | 53.5, CH |
| | 3 | 2.04, m | 29.9, CH | 1.95, m | 29.9, CH |
| | 4 | 0.94, m | 18.9, $CH_3$ | n.o | |
| | 5 | 0.94, m | 18.9, $CH_3$ | n.o | |
| | NH | 8.03, br | | 8.02, br | |
| N-diMe-Ile | 1 | | 169.8, qC | | 169.8, qC |
| | 2 | 2.77, m | 68.1, CH | n.o | |
| | 3 | 1.50, m | 37.4, CH | n.o | |
| | 4 | 0.80, m | 15.2, $CH_3$ | n.o | |
| | 5a | 1.54, m | 24.8, $CH_2$ | n.o | |
| | 5b | 1.10, m | | n.o | |
| | 6 | 0.82, m | 11.1, $CH_3$ | n.o | |
| | 7 (N-Me) | 2.17, s | 34.5, $CH_3$ | 2.15, s | 34.4 |

**Table. S3.** NMR spectroscopic data of des-Aph-aetokthonostatin (**3**; major and minor conformer). All spectra were recorded in DMSO-*d6* ($^{1}$H 600 MHz, $^{13}$C 150 MHz). n.o. not observed.

| | conformer 1 | conformer 2 |
|---|---|---|

| unit | C/H no. | $\delta_H$ (*J* in Hz) | $\delta_C$, mult | $\delta_H$ (*J* in Hz) | $\delta_C$, mult |
|---|---|---|---|---|---|
| Dap | 1 | | 175.8, qC | n.o | |
| | 2 | 2.29, m | 42.6, CH | n.o | |
| | 3 | 1.09, m | 13.0, $CH_3$ | 1.12, m | 14.6, $CH_3$ |
| | 4 | 3.83, m | 81.1, CH | n.o | |
| | 5 | 4.00, m | 58.6, CH | 4.05, m | 58.8, CH |
| | 6a | 1.91, m | 24.5, $CH_2$ | n.o | |
| | 6b | 1.77, m | | n.o | |
| | 7a | 1.93, m | 23.9, $CH_2$ | n.o | |
| | 7b | 1.74, m | | n.o | |
| | 8a | 3.52, m | 46.7, $CH_2$ | 3.61, m | 45.8, $CH_2$ |
| | 8b | 3.35, m | | 3.15, m | |
| | 9 (O-Me) | 3.28, s | 59.7, $CH_3$ | 3.30, s | 60.4, $CH_3$ |
| Dil | 1 | | 169.2, qC | n.o | |
| | 2a | 2.45, m | 36.8, $CH_2$ | 2.43, m | 36.7, $CH_2$ |
| | 2b | 2.34, m | | 2.31, m | |
| | 3 | 3.98, m | 77.5, CH | n.o | |
| | 4 | 4.63, m | 55.5, CH | 4.67, m | 55.2, CH |
| | 5 | 1.76, m | 31.9, CH | 1.81, m | 31.7, CH |
| | 6 | 0.90, m | 14.9, $CH_3$ | n.o | |
| | 7a | 1.30, m | 25.0, $CH_2$ | n.o | |
| | 7b | 0.90, m | | n.o | |
| | 8 | 0.76, m | 10.1, $CH_3$ | n.o | |
| | 9 (O-Me) | 3.19, s | 56.9, $CH_3$ | 3.18, s | 56.9, $CH_3$ |
| | 10 (N-Me) | 3.00, s | 31.3, $CH_3$ | 3.05, s | 31.3, $CH_3$ |
| Val | 1 | | 173.0, qC | | 172.9, qC |
| | 2 | 4.54, m | 53.5, CH | n.o | |
| | 3 | 1.95, m | 29.6, CH | n.o | |
| | 4 | 0.92, m | 18.6, $CH_3$ | n.o | |
| | 5 | 0.92, m | 18.6, $CH_3$ | n.o | |
| | NH | 8.02, d (8.30) | | n.o | |
| N-diMe-Ile | 1 | | 169.8, qC | | 169.8, qC |
| | 2 | 2.77, m | 70.4, CH | n.o | |
| | 3 | 1.74, m | 32.2, CH | n.o | |
| | 4 | 0.69, m | 14.9, $CH_3$ | n.o | |
| | 5a | 1.57, m | 24.5, $CH_2$ | n.o | |
| | 5b | 1.07, m | | n.o | |
| | 6 | 0.83, m | 10.1, $CH_3$ | n.o | |
| | 7 (N-Me) | 2.21, m | 41.3, $CH_3$ | n.o | |
| | 8 (N-Me) | 2.21, m | 41.3, $CH_3$ | n.o | |

**Table. S4.** Comparison of the $^{13}C$ NMR spectroscopic data of AEST (major conformer, spectra recorded in $CD_2Cl_2$,150 MHz) and symplostatin 1 (literature data, ref. 23 main manuscript). n.o. not observable due to solvent signal. Spectra see Fig. S6b and S6c.

| | | **AEST** | **Symplostatin 1** |
|---|---|---|---|
| **unit** | **C no.** | **$\delta_C$** | **$\delta_C$** |
| Aph | 1 | n.o. | |
| | 2 | 38.7 | |
| | 3 | 139.4 | |
| | 4 | 129.9 | |
| | 5 | 128.8 | |
| | 6 | 126.6 | |
| | 7 | 76.3 | |
| | 8 | 16.1 | |
| | 9 | 62.0 | |
| Dap | 1 | 174.1 | 174.0 |
| | 2 | 45.4 | 44.7 |
| | 3 | 15.0 | 14.5 |
| | 4 | 82.4 | 81.9 |
| | 5 | 59.9 | 59.7 |
| | 6 | 25.3 | 24.9 |
| | 7 | 25.5 | 25.4 |
| | 8 | 48.2 | 48.0 |
| | 9 (O-Me) | 61.1 | 60.9 |
| Dil | 1 | 170.1 | 170.4 |
| | 2 | 38.1 | 37.9 |
| | 3 | 78.7 | 78.7 |
| | 4 | 57.1 | 57.2 |
| | 5 | 33.8 | 33.4 |
| | 6 | 15.8 | 15.9 |
| | 7 | 26.2 | 26.1 |
| | 8 | 10.8 | 10.8 |
| | 9 (O-Me) | 58.3 | 58.1 |
| | 10 (N-Me) | 32.2 | 32.3 |
| Val | 1 | 173.7 | 173.4 |
| | 2 | 54.7 | 54.5 |
| | 3 | 31.5 | 31.2 |
| | 4 | 18.2 | 18.3 |
| | 5 | 20.0 | 19.6 |
| N-diMe-Ile | 1 | 174.2 | 174.0 |
| | 2 | 75.4 | 74.5 |
| | 3 | 34.9 | 34.5 |
| | 4 | 15.1 | 15.0 |
| | 5 | 26.4 | 26.8 |
| | 6 | 12.1 | 11.8 |
| | 7 (N-Me) | 43.3 | 42.6 |

**Table. S5.** Summary of encoded proteins in the aetokthonostatin biosynthetic pathway and their closest homologues. NRPS, non-ribosomal peptide synthetase; PKS, polyketide synthase; SAM, S-adenosylmethionine; SDR, short-chain dehydrogenase/reductase; TE, thioesterase.

| **Aetokthonostatin pathway** | | | **Top BLASTp hit** | | | | **Top MIBiG hit with known function** | | | |
|---|---|---|---|---|---|---|---|---|---|---|
| **Protein** | **Size [aa]** | **Predicted Function** | **Accession** | **Organism** | **Identity [%]** | **Annotation** | **Accession** | **Organism** | **Identity [%]** | **Function** |
| ORF1 | 252 | SDR reductase | MBW4632174.1 | *Iphinoe* sp. HA4291-MV1 | 63.9 | SDR family oxidoreductase | AFN69430.1 | *Staphylococcus epidermidis* | 41.0 | ElxO (epilancin 15x); SDR - reduction of pyruvyl to lactyl |
| AesA | 1120 | NRPS | NER93225.1 | *Symploca* sp. SIO1B1 | 64.3 | amino acid adenylation domain-containing protein | AEU11003.1 | *Nostoc* sp. 152 | 49.0 | NpnC (nostophycin); NRPS |
| AesB | 1552 | PKS | NES18617.1 | *Caldora* sp. (= *Symploca* sp.) SIO3E6 | 65.1 | acyltransferase domain-containing protein | WP_035122279.1 | *Fischerella* sp. PCC 9431 | 51.0 | HapE (hapalosin); type I PKS |
| AesC | 1121 | NRPS | NER93223.1 | *Symploca* sp. SIO1B1 | 52.4 | amino acid adenylation domain-containing protein | AAO23334.1 | *Nostoc* sp. ATCC 53789 | 49.0 | NcpB (nostocyclopeptide A2); NRPS |
| AesD | 1320 | PKS + TE | NEQ64994.1 | *Symploca* sp. SIO2D2 | 55.3 | acyltransferase domain-containing protein | AAS98787.1 | *Lyngbya majuscula* | 66.0 | JamP (jamaicamides); PKS + TE |
| AesE | 547 | SAM-dependent *C*-methyltransferase | NES20410.1 | *Caldora* sp. (= *Symploca* sp.) SIO3E6 | 60.1 | class I SAM-dependent methyltransferase | AHA38199.1 | *Archangium violaceum* Cb vi76 | 27.0 | GphF (gephyronic acid); *C*-methyltransferase |
| AesF | 1039 | NRPS (starter unit) | NEP61017.1 | *Symploca* sp. SIO2G7 | 59.7 | amino acid adenylation domain-containing protein | AWI62629.1 | *Cystobacter* sp. | 44.0 | VioD (vioprolides); NRPS |
| AesG | 2657 | NRPS | NER93229.1 | *Symploca* sp. SIO1B1 | 60.3 | amino acid adenylation domain-containing protein | ABI26078.1 | *Planktothrix agardhii* NIVA-CYA 116 | 46.0 | OciB (cyanopeptin); NRPS |
| AesH | 281 | SAM-dependent *O*-methyltransferase | NEP58941.1 | *Symploca* sp. SIO2G7 | 72.0 | class I SAM-dependent methyltransferase | CAL69886.1 | *Paraburkholderia rhizoxinica* | 43.0 | RhiI (rhizoxin A); *O*-methyltransfearse |
| AesI | 325 | SAM-dependent *O*-methyltransferase | NEP58940.1 | *Symploca* sp. SIO2G7 | 61.4 | class I SAM-dependent methyltransferase | CAJ77713.1 | *Mycobacteroides abscessus* | 35.0 | Fmt (glycopeptidolipid); *O*-methyltransferase |
| AesJ | 1438 | PKS | WP_229547965.1 | *Nostoc* sp. CHAB 5836 | 47.2 | SDR family NAD(P)-dependent oxidoreductase | WP_035122279.1 | *Fischerella* sp. PCC 9431 | 46.0 | HapE (hapalosin); type I PKS |
| AesK | 253 | SAM-dependent *N*-methyltransferase (monomethylaetokthonostatin *N*-methyltransferase) | WP_017308077.1 | *Fischerella* sp. PCC 9339 | 65.5 | class I SAM-dependent methyltransferase | DAB41913.1 | *Fischerella* sp. PCC 9339 | 66.0 | ArzK (aranazoles); *O*-methyltransferase |

**Table. S6.** Summary of encoded proteins in the putative dolastatin analogue biosynthetic pathway and their closest homologues in *Symploca* sp. SIO1B1 (JAAHFO010000022.1). NRPS, non-ribosomal peptide synthetase; PKS, polyketide synthase; SAM, S-adenosylmethionine.

| **Putative dolastatin family-compound pathway** | | | **Top BLASTp hit (except the ortholog from *Symploca* sp. SIO1C2)** | | | | **Top MIBiG hit with known function** | | | |
|---|---|---|---|---|---|---|---|---|---|---|
| **Protein** | **Size [aa]** | **Predicted Function** | **Accession** | **Organism** | **Identity [%]** | **Annotation** | **Accession** | **Organism** | **Identity [%]** | **Function** |
| SymE | 552 | SAM-dependent *C*-methyltransferase | NES20410.1 | *Caldora* sp. (= *Symploca* sp.) SIO3E6 | 83.5 | class I SAM-dependent methyltransferase | ABI75094.1 | *Cylindrospermopsis raciborskii* T3 | 31.0 | SxtA (saxitoxin); PKS (including methyltransferase) |
| SymF | 1047 | NRPS (starter unit) | NES20409.1 | *Caldora* sp. (= *Symploca* sp.) SIO3E6 | 85.4 | amino acid adenylation domain-containing protein | AZH23822.1 | *Okeania hirsuta* | 41.0 | MgiJ (malyngamide); NRPS |
| SymG | 2701 | NRPS | NES19078.1 | *Caldora* sp. (= *Symploca* sp.) SIO3E6 | 90.0 | amino acid adenylation domain-containing protein | ABI26078.1 | *Planktothrix agardhii* NIVA-CYA 116 | 44.0 | OciB (cyanopeptin); NRPS |
| SymH | 277 | SAM-dependent *O*-methyltransferase | NEP58941.1 | *Symploca* sp. SIO2G7 | 92.8 | class I SAM-dependent methyltransferase | ADZ25003.1 | *Sorangium cellulosum* | 38.0 | Leu14 (leupyrrins); *O*-methyltransfearse |
| SymI | 289 | SAM-dependent *O*-methyltransferase | NEP58940.1 | *Symploca* sp. SIO2G7 | 96.5 | class I SAM-dependent methyltransferase | CAD15506.1 | *Ralstonia solanacearum* GMI1000 | 38.0 | RSc1804 (micacocidin); NRPS/PKS (inlcuding SAM-dependent methyltransferase) |
| SymJ | 1559 | PKS | NEP58939.1 | *Symploca* sp. SIO2G7 | 92.9 | SDR family NAD(P)-dependent oxidoreductase | WP_035122279.1 | *Fischerella* sp. PCC 9431 | 51.0 | HapE (hapalosin); type I PKS |
| SymA | 1118 | NRPS | NES24143.1 | *Caldora* sp. (= *Symploca* sp.) SIO3E6 | 93.9 | AMP-binding protein | AEU11003.1 | *Nostoc* sp. 152 | 50.0 | NpnC (nostophycin); NRPS |
| SymB | 1556 | PKS | NEP59557.1 | *Symploca* sp. SIO2G7 | 94.9 | acyltransferase domain-containing protein | WP_035122279.1 | *Fischerella* sp. PCC 9431 | 52.0 | HapE (hapalosin); type I PKS |
| SymC | 2487 | NRPS | NEO91324.1 | *Moorena* sp. SIO3G5 | 76.1 | amino acid adenylation domain-containing protein | AAN32981.1 | *Lyngbya majuscula* | 56.0 | BarG (barbamide); NRPS |
| ORF1 | 361 | hydrolase | NER51533.1 | *Symploca* sp. SIO1A3 | 97.8 | amidohydrolase | AAN32982.1 | *Lyngbya majuscula* | 79.0 | BarH (barbamide); hydrolase |
| ORF2 | 77 | unknown | NES19872.1 | *Caldora* sp. (= *Symploca* sp.) SIO3E6 | 93.3 | cyclase family protein | NA | NA | NA | NA |
| SymK | 270 | SAM-dependent *N*-methyltransferase | NEO41470.1 | *Moorena* sp. SIOASIH | 84.8 | class I SAM-dependent methyltransferase | AAA67510.1 | *Streptomyces glaucescens* | 28.0 | TcmP (tetracenomycin C); *O*-methyltransferase |
| ORF3 | 277 | unknown | NEQ71097.1 | *Symploca* sp. SIO2D2 | 97.8 | cyclase family protein | ABX24504.1 | *Streptomyces cacaoi* subsp. *asoensis* | 49.0 | Orf1 (polyoxins); putative cyclase |

**Table. S7.** Summary of encoded proteins in the putative dolastatin analogue biosynthetic pathway and their closest homologues in *Symploca* sp. SIO1C2 (JAAHFP010000047.1). NRPS, non-ribosomal peptide synthetase; PKS, polyketide synthase; SAM, S-adenosylmethionine.

| Putative dolastatin family-compound pathway | | | Top BLASTp hit (except the ortholog from *Symploca* sp. SIO1B1) | | | | Top MIBiG hit with known function | | | |
|---|---|---|---|---|---|---|---|---|---|---|
| **Protein** | **Size [aa]** | **Predicted Function** | **Accession** | **Organism** | **Identity [%]** | **Annotation** | **Accession** | **Organism** | **Identity [%]** | **Function** |
| SymE | 552 | SAM-dependent *C*-methyltransferase | NES20410.1 | *Caldora* sp. (= *Symploca* sp.) SIO3E6 | 83.5 | class I SAM-dependent methyltransferase | ABI75094.1 | *Cylindrospermopsis raciborskii* T3 | 31.0 | SxtA (saxitoxin); PKS (including methyltransferase) |
| SymF | 1047 | NRPS (starter unit) | NES20409.1 | *Caldora* sp. (= *Symploca* sp.) SIO3E6 | 85.4 | amino acid adenylation domain-containing protein | AAO62586.1 | *Anabaena* sp. 90 | 43.0 | McyA (microcystin); NRPS |
| SymG | 2709 | NRPS | NES19078.1 | *Caldora* sp. (= *Symploca* sp.) SIO3E6 | 90.0 | amino acid adenylation domain-containing protein | ABI26078.1 | *Planktothrix agardhii* NIVA-CYA 116 | 44.0 | OciB (cyanopeptin); NRPS |
| SymH | 277 | SAM-dependent *O*-methyltransferase | NEP58941.1 | *Symploca* sp. SIO2G7 | 92.8 | class I SAM-dependent methyltransferase | CAJ77713.1 | *Mycobacteroides abscessus* | 37.0 | Fmt (glycopeptidolipid); *O*-methyltransferase |
| SymI | 289 | SAM-dependent *O*-methyltransferase | NEP58940.1 | *Symploca* sp. SIO2G7 | 96.5 | class I SAM-dependent methyltransferase | CAD15506.1 | *Ralstonia solanacearum* GMI1000 | 39.0 | RSc1804 (micacocidin); NRPS/PKS (inlcuding SAM-dependent methyltransferase) |
| SymJ | 1559 | PKS | NEP58939.1 | *Symploca* sp. SIO2G7 | 92.9 | SDR family NAD(P)-dependent oxidoreductase | WP_035122279.1 | *Fischerella* sp. PCC 9431 | 50.0 | HapE (hapalosin); type I PKS |
| SymA | 1118 | NRPS | NES24143.1 | *Caldora* sp. (= *Symploca* sp.) SIO3E6 | 93.9 | AMP-binding protein | AEU11003.1 | *Nostoc* sp. 152 | 50.0 | NpnC (nostophycin); NRPS |
| SymB | 1556 | PKS | NEP59557.1 | *Symploca* sp. SIO2G7 | 94.9 | acyltransferase domain-containing protein | WP_035122279.1 | *Fischerella* sp. PCC 9431 | 52.0 | HapE (hapalosin); type I PKS |
| SymC/D | 2545 | NRPS/PKS | NES18616.1 | *Caldora* sp. (= *Symploca* sp.) SIO3E6 | 90.3 | SDR family NAD(P)-dependent oxidoreductase | AAY42398.1 | *Lyngbya majuscula* | 47.0 | HctF (hectochlorin); NRPS |
| SymK | 270 | SAM-dependent *N*-methyltransferase | WP_075896359.1 | *Moorena bouillonii* | 79.6 | class I SAM-dependent methyltransferase | AAA67510.1 | *Streptomyces glaucescens* | 29.0 | TcmP (tetracenomycin C); *O*-methyltransferase |
| ORF1 | 277 | unknown | NEQ71335.1 | *Symploca* sp. SIO2D2 | 99.3 | cyclase family protein | ABX24504.1 | *Streptomyces cacaoi* subsp. *asoensis* | 49.0 | Orf1 (polyoxins); putative cyclase |

### SUPPLEMENTARY FIGURES – Bioactivity characterization

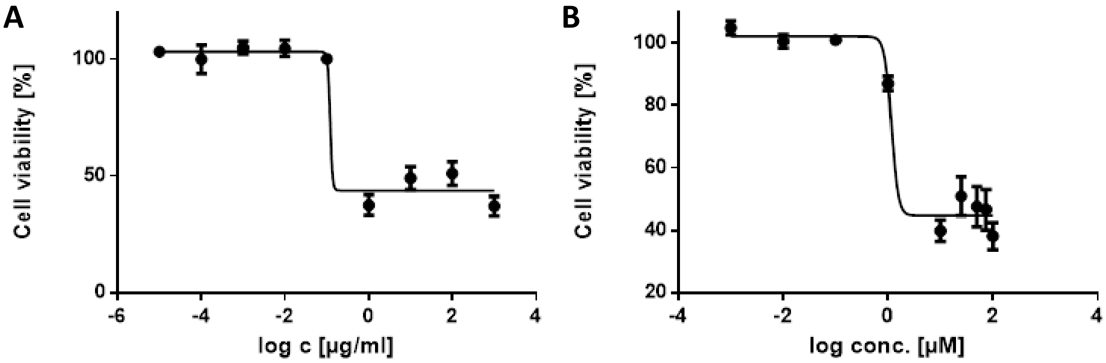


**Fig. S2.** Cytotoxicity of *A. hydrillicola* extract (**A**, $EC_{50}$ 0.12 µg/mL) and pure AETX (**B**, $EC_{50}$ 1 µM; corresponds to about 0.65 µg/mL). HeLa cells, SRB assay.

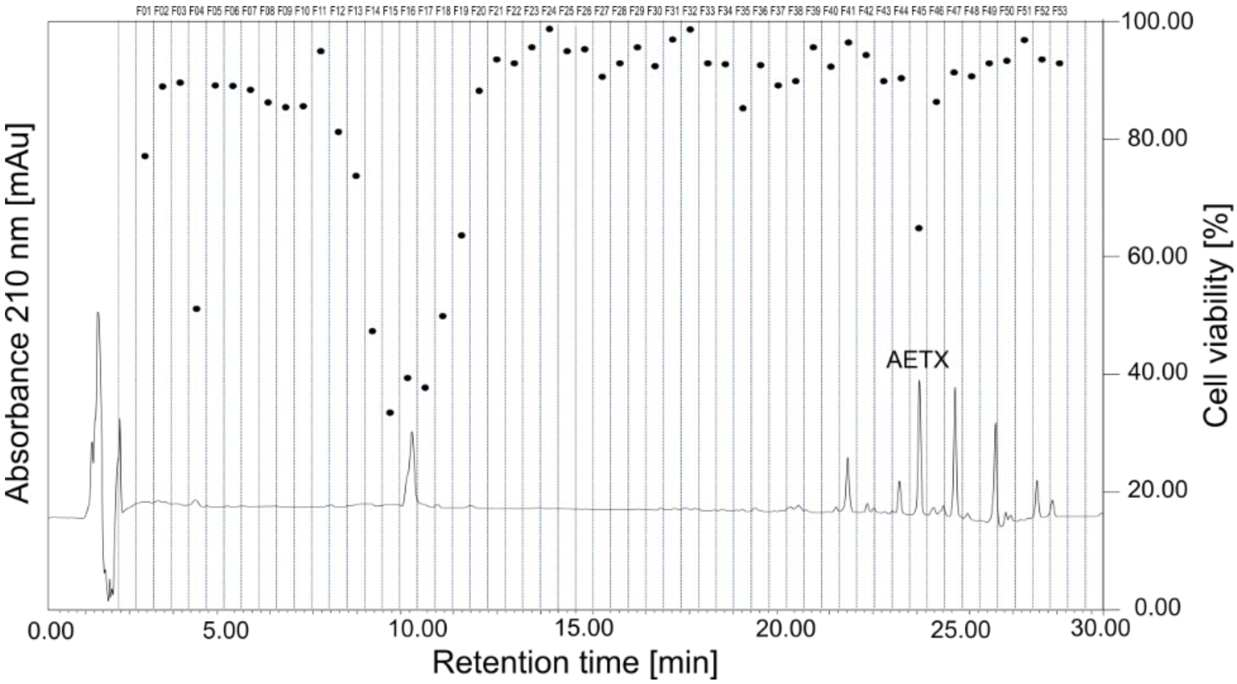


**Fig. S3.** Microfractionation and results from the cytotoxicity assay of an *A. hydrillicola* extract.

A

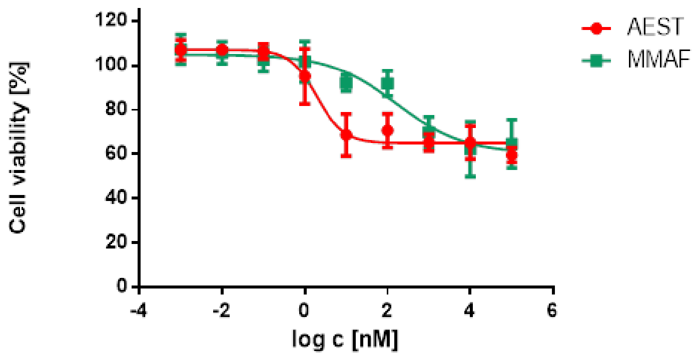


B

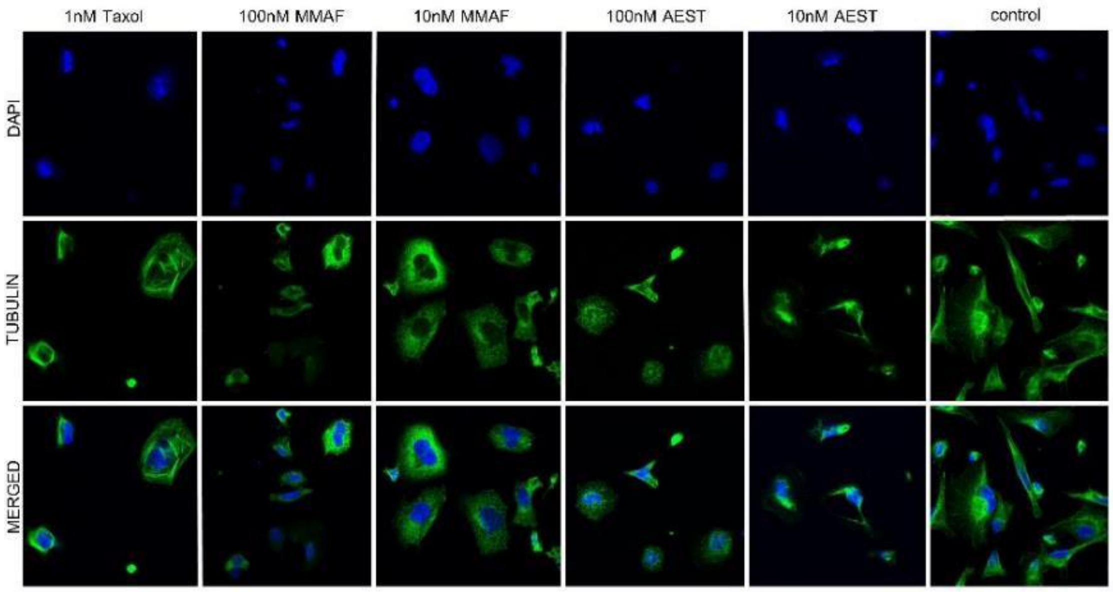


**Fig. S4. (A)** Cytotoxicity data of AEST and MMAF against MDA-MB 231 breast cancer cells. (B) Immuno-fluorescence microscopy of MDA-MB 231 cells showing the effect of AEST and MMAF on tubulin (in green); nuclei were stained with DAPI (in blue), taxol (1 nM) was used as positive control.

### SUPPLEMENTARY FIGURES – Structure elucidation

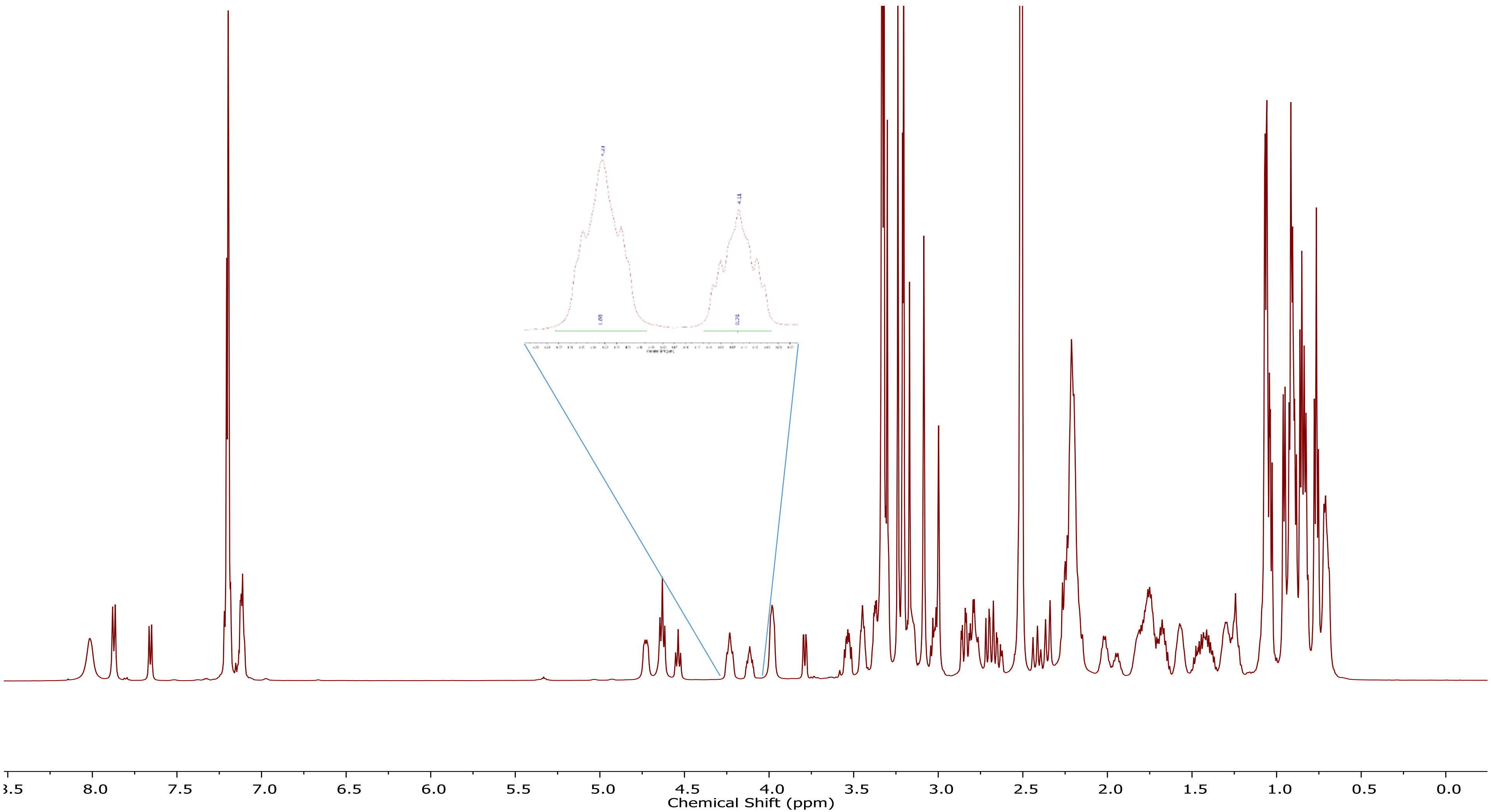


**Fig. S5.** $^{1}$H NMR spectrum of aetokthonostatin (**1**) in DMSO-$d_6$ (600 MHz). Inset: Comparison of the integrals of a proton of main and minor conformer. See table S1 for a list of the observed chemical shifts.

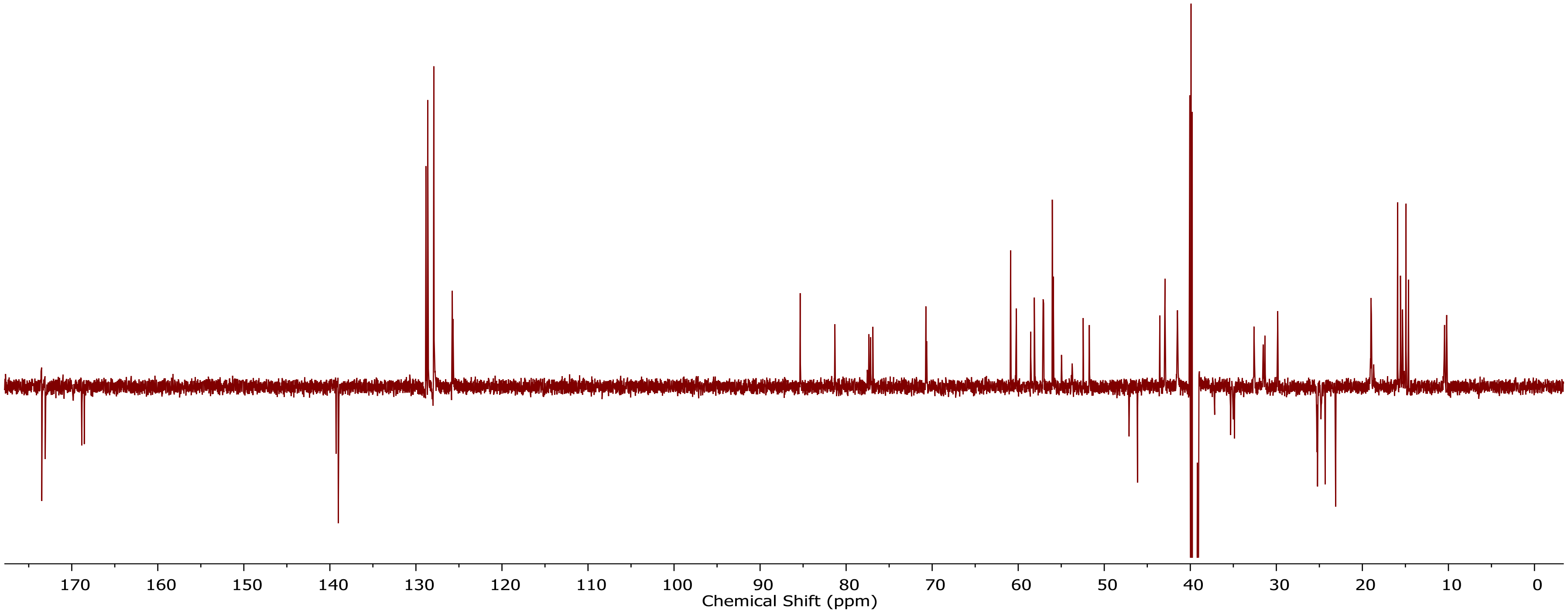


**Fig. S6a.** $^{13}C$-APT NMR spectrum of aetokthonostatin (**1**) in DMSO-$d_6$ (150 MHz). See table S1 for a list of the observed chemical shifts.

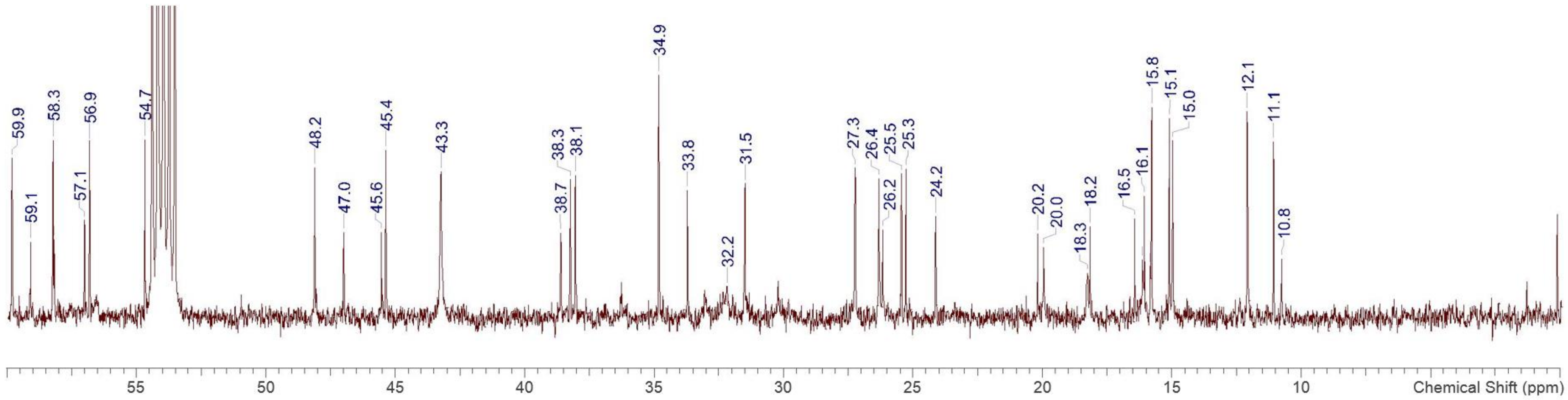


**Fig. S6b.** $^{13}C$ NMR spectrum of aetokthonostatin (**1**) in $CD_2Cl_2$ (150 MHz) from 0 to 60 ppm.

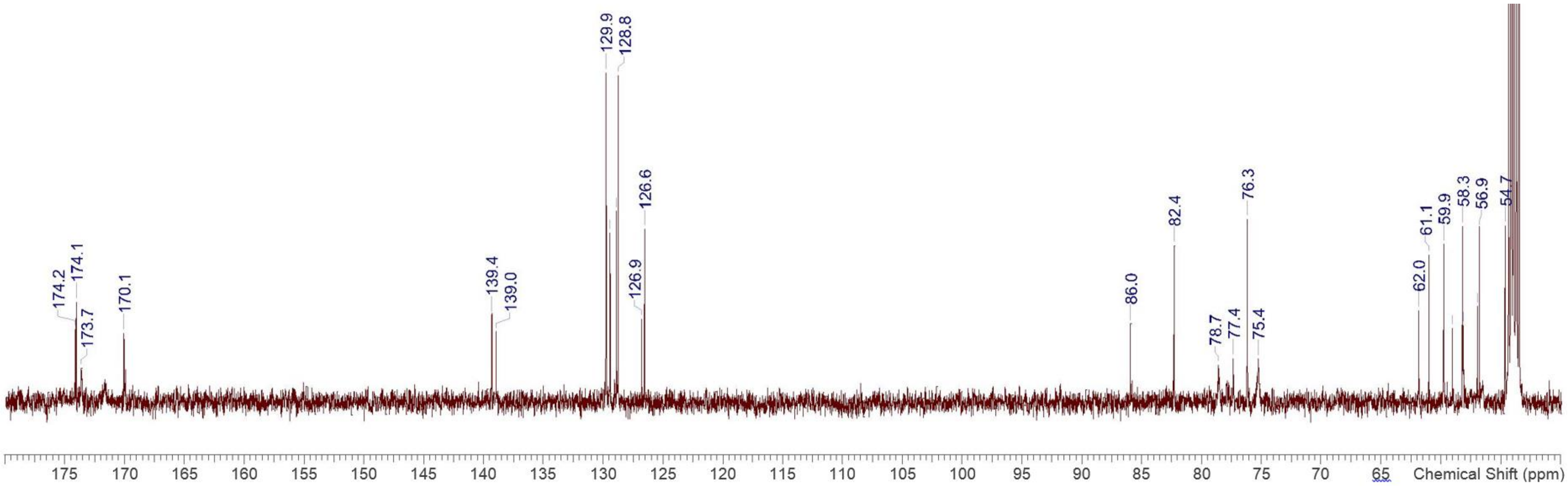


**Fig. S6c.** $^{13}C$ NMR spectrum of aetokthonostatin (**1**) in $CD_2Cl_2$ (150 MHz) from 50 to 180 ppm.

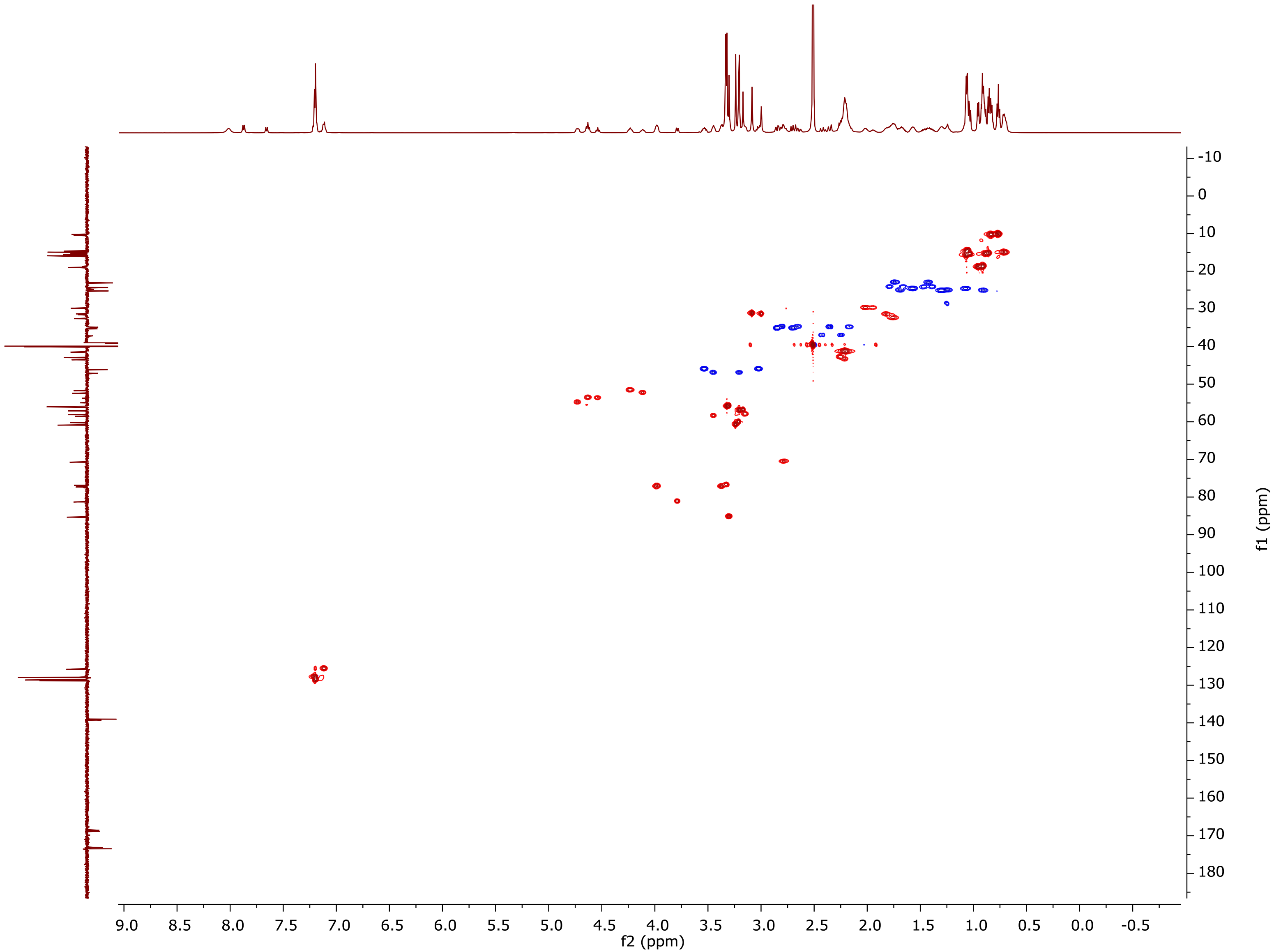


**Fig. S7.** $^{13}$C-HSQC NMR spectrum of aetokthonostatin (**1**) in DMSO-$d_6$ (600 MHz). See table S1 for a list of the observed chemical shifts.

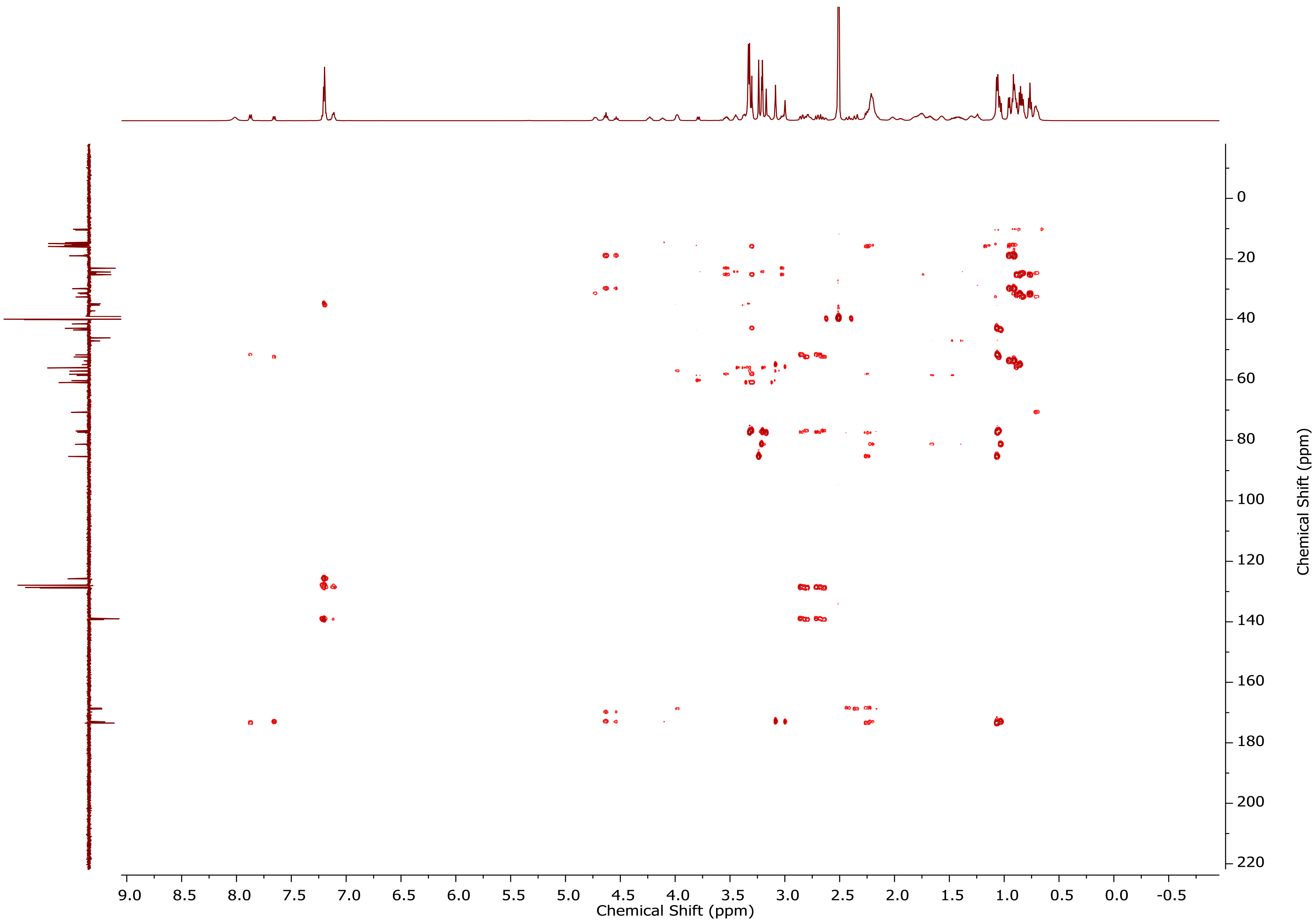


**Fig. S8.** $^{13}$C-HMBC NMR spectrum of aetokthonostatin (**1**) in DMSO-$d_6$ (600 MHz). See table S1 for a list of the observed chemical shifts.

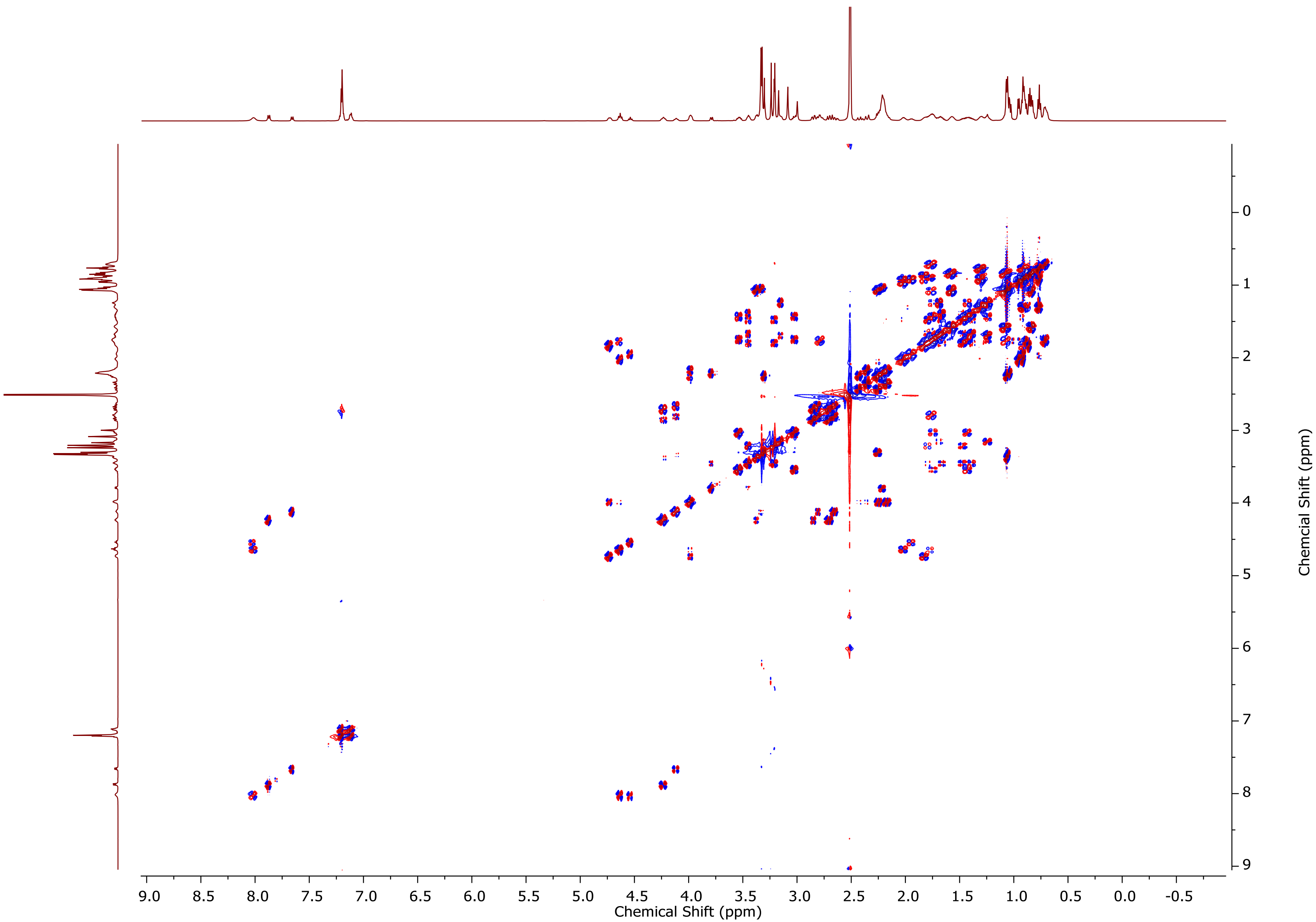


**Fig. S9.** DQF-COSY NMR spectrum of aetokthonostatin (**1**) in DMSO-$d_6$ (600 MHz).

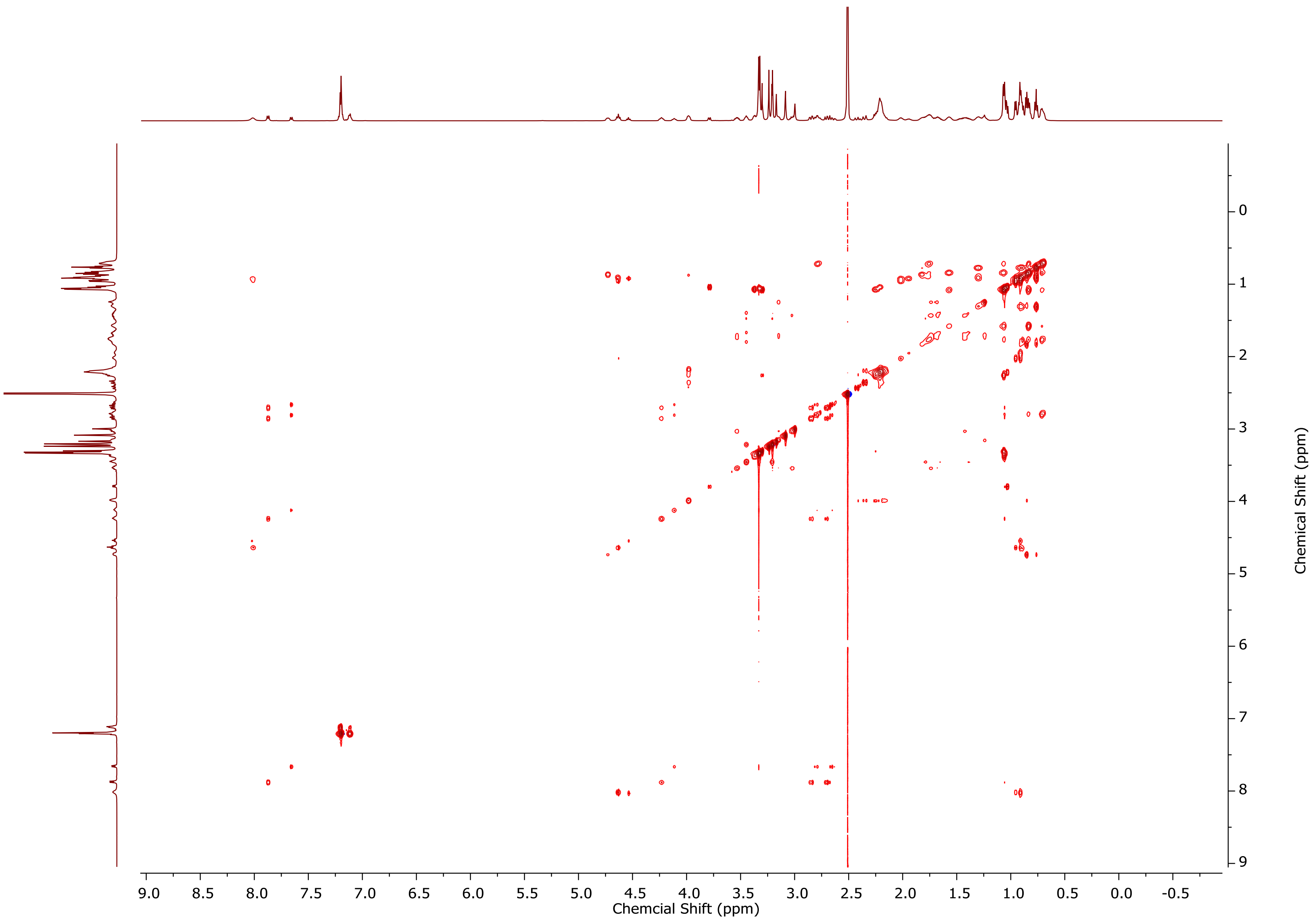


**Fig. S10.** TOCSY NMR spectrum of aetokthonostatin (**1**) in DMSO-$d_6$ (600 MHz).

A

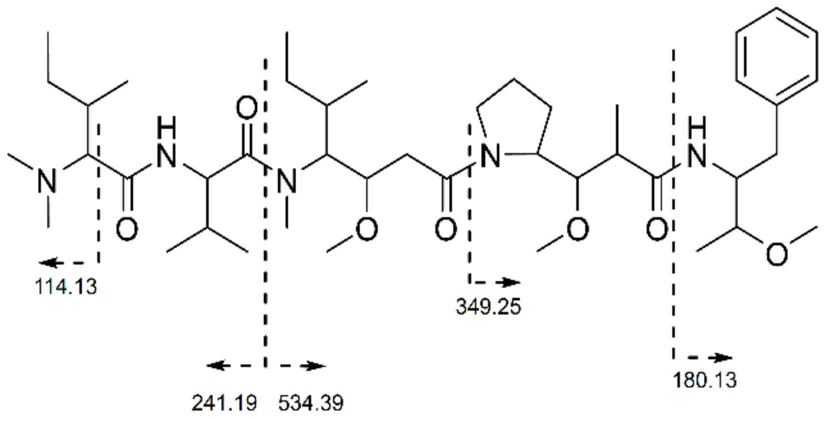


B

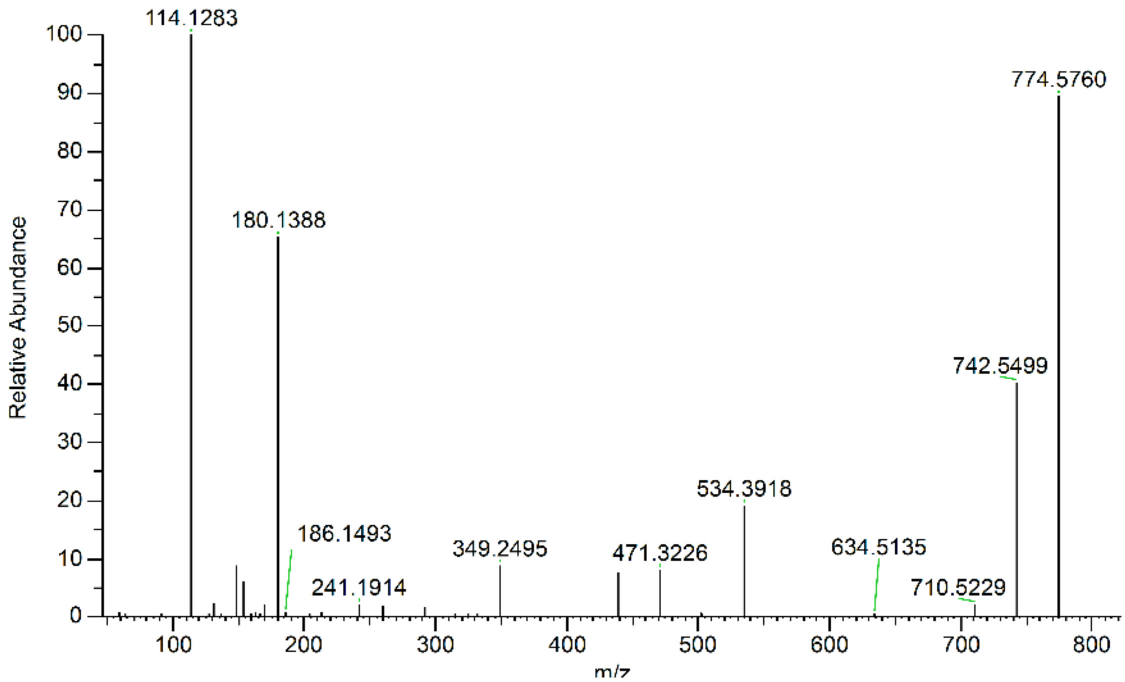


**Fig. S11. (A)** Key MS/MS fragments and **(B)** MS/MS spectrum of aetokthonostatin (**1**).

A

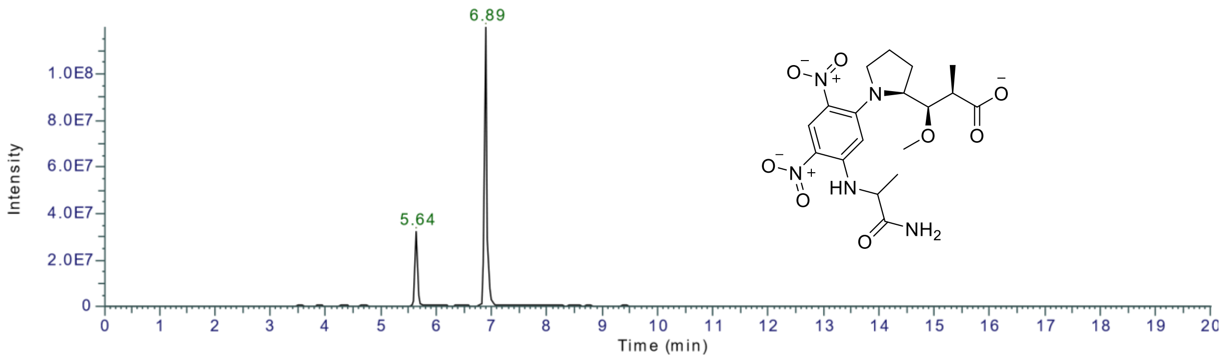


B

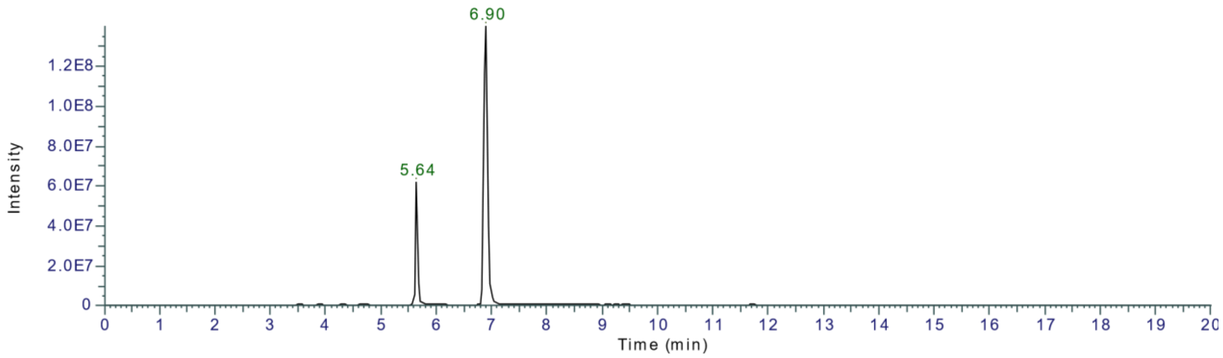


**Fig. S12.** Marfey's analysis of dolaproine. EICs of Dap derivatized with Marfey's reagent (*m/z* 438.1630, structure shown in the chromatogram) in **(A)** AEST (**1**) and **(B)** MMAF.

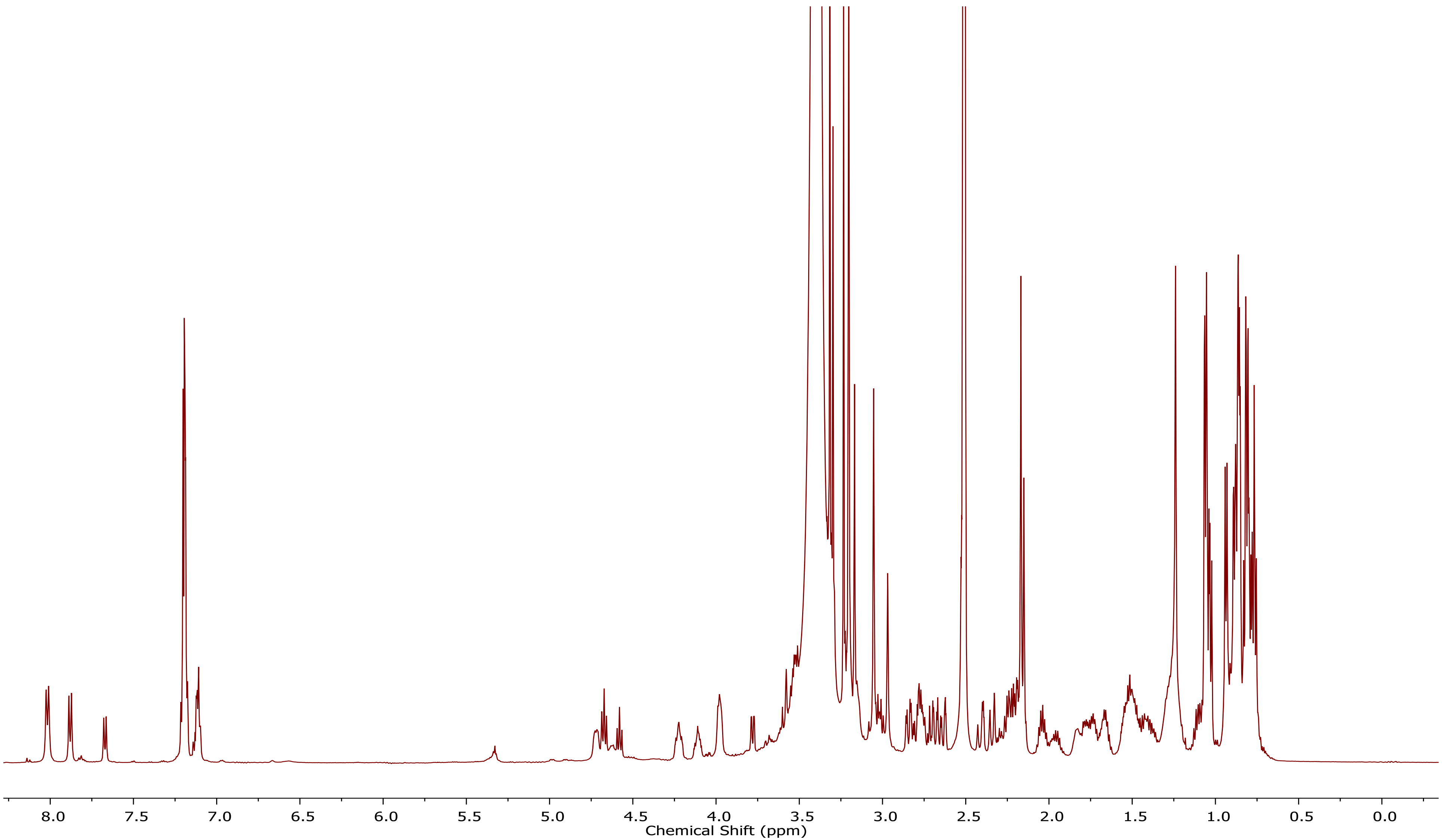


**Fig. S13.** $^1$H NMR spectrum of monomethyl-aetokthonostatin (**2**) in DMSO-$d_6$ (600 MHz). See table S2 for a list of the observed chemical shifts.

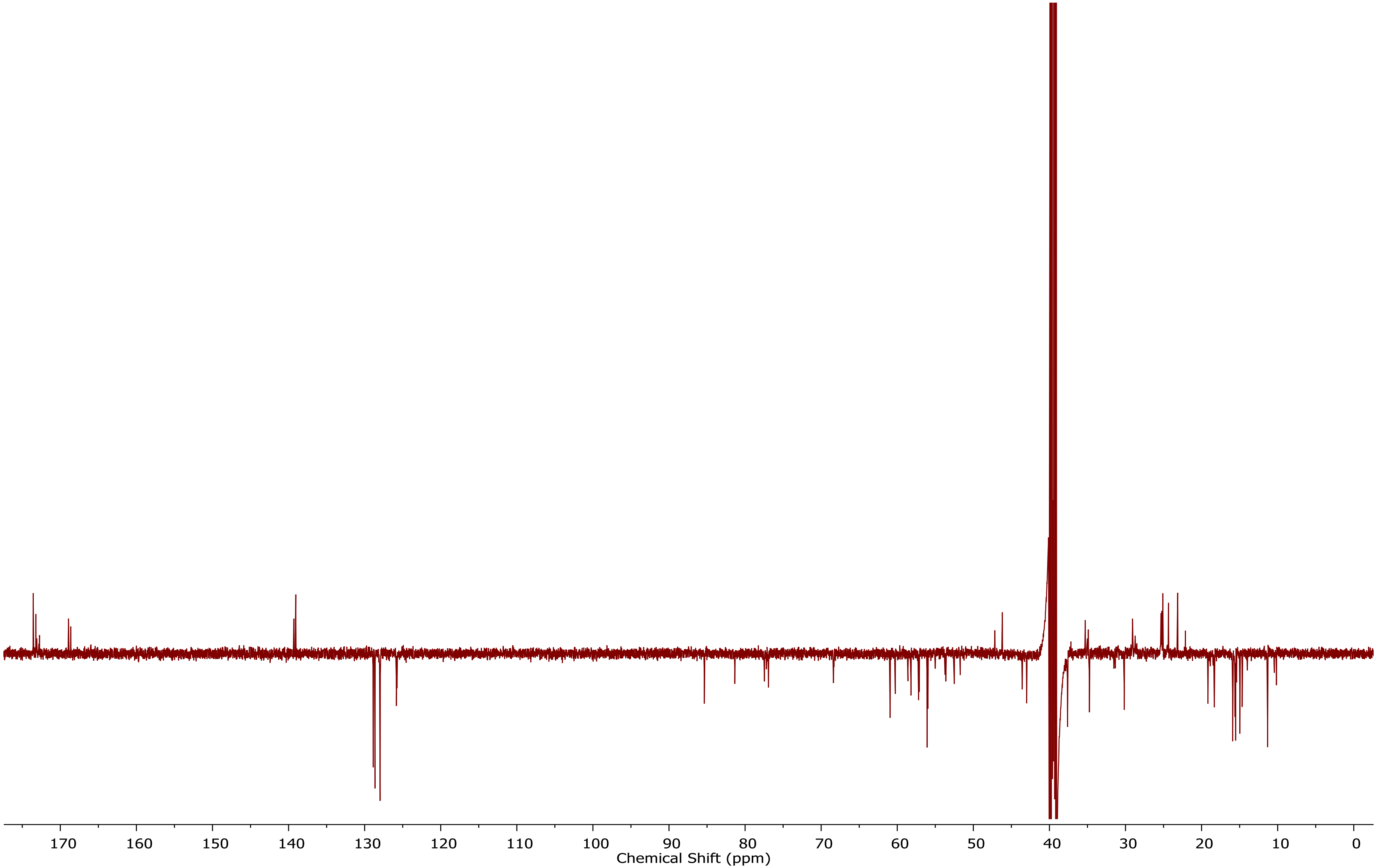


**Fig. S14.** $^{13}$C-ATP NMR spectrum of monomethyl-aetokthonostatin (**2**) in DMSO-$d_6$ (150 MHz). See table S2 for a list of the observed chemical shifts.

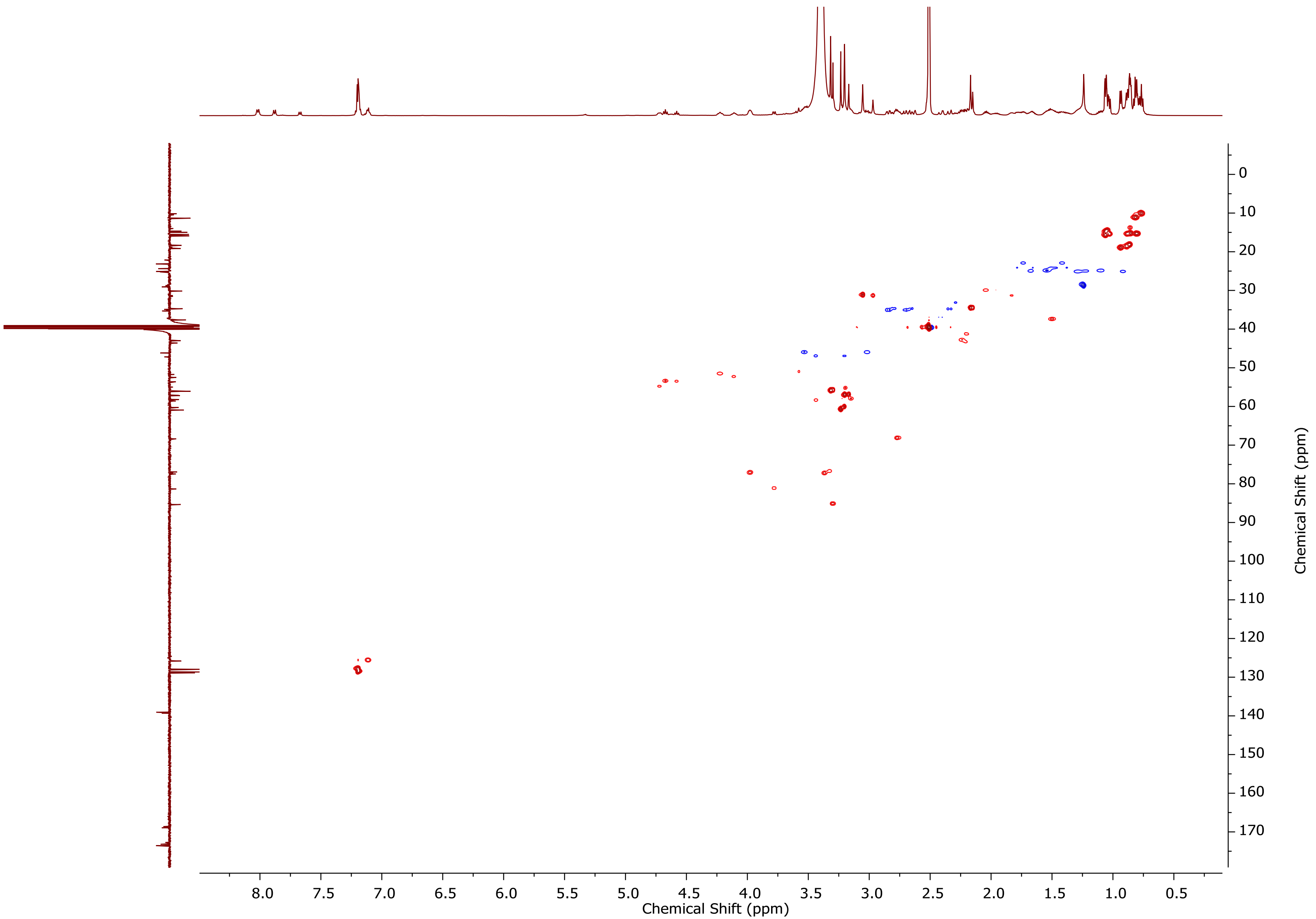


**Fig. S15.** $^{13}C$-HSQC NMR spectrum of monomethyl-aetokthonostatin (**2**) in DMSO-$d_6$ (600 MHz). See table S2 for a list of the observed chemical shifts.

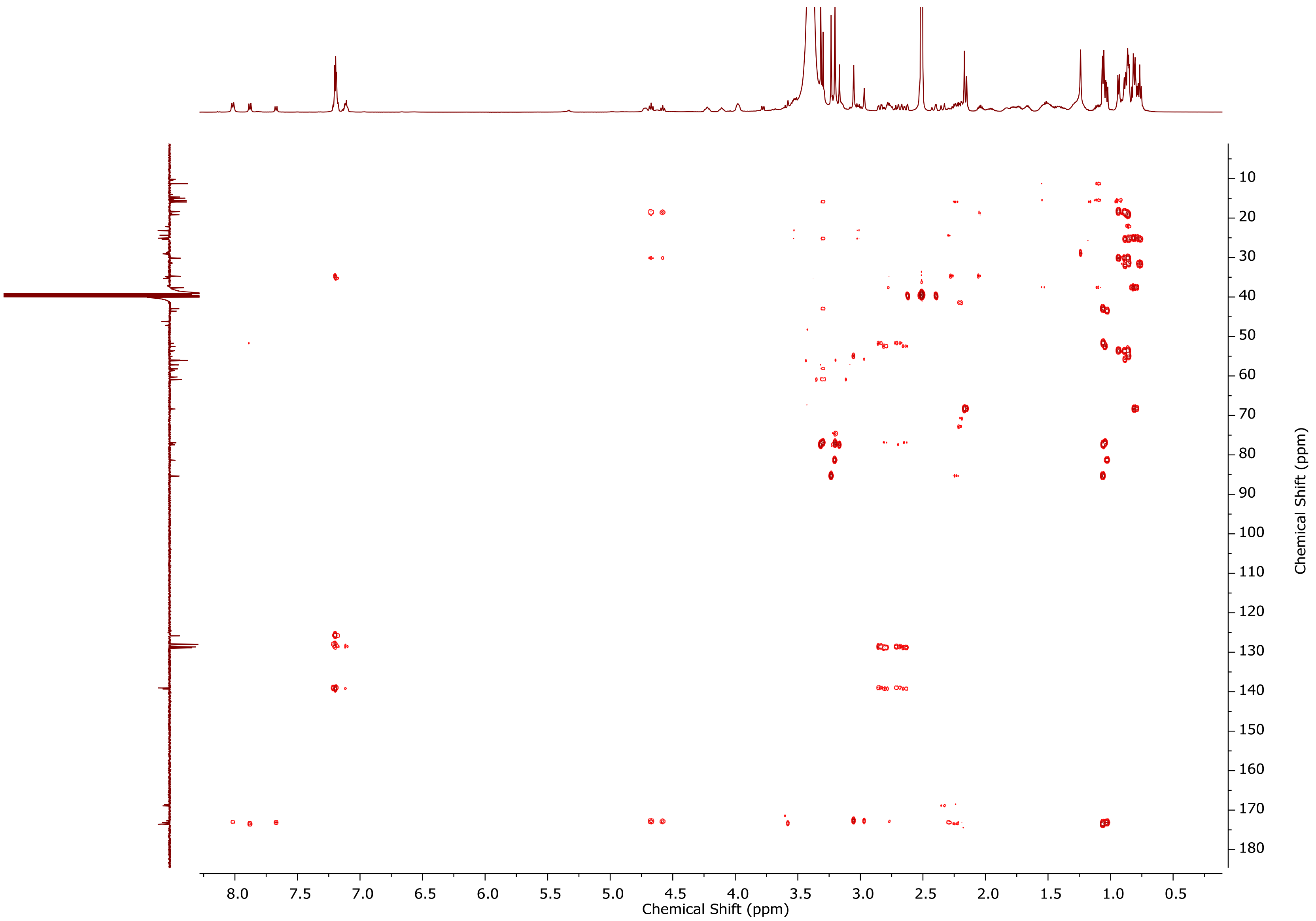


**Fig. S16.** $^{13}$C-HMBC NMR spectrum of monomethyl-aetokthonostatin (**2**) in DMSO-$d_6$ (600 MHz). See table S2 for a list of the observed chemical shifts.

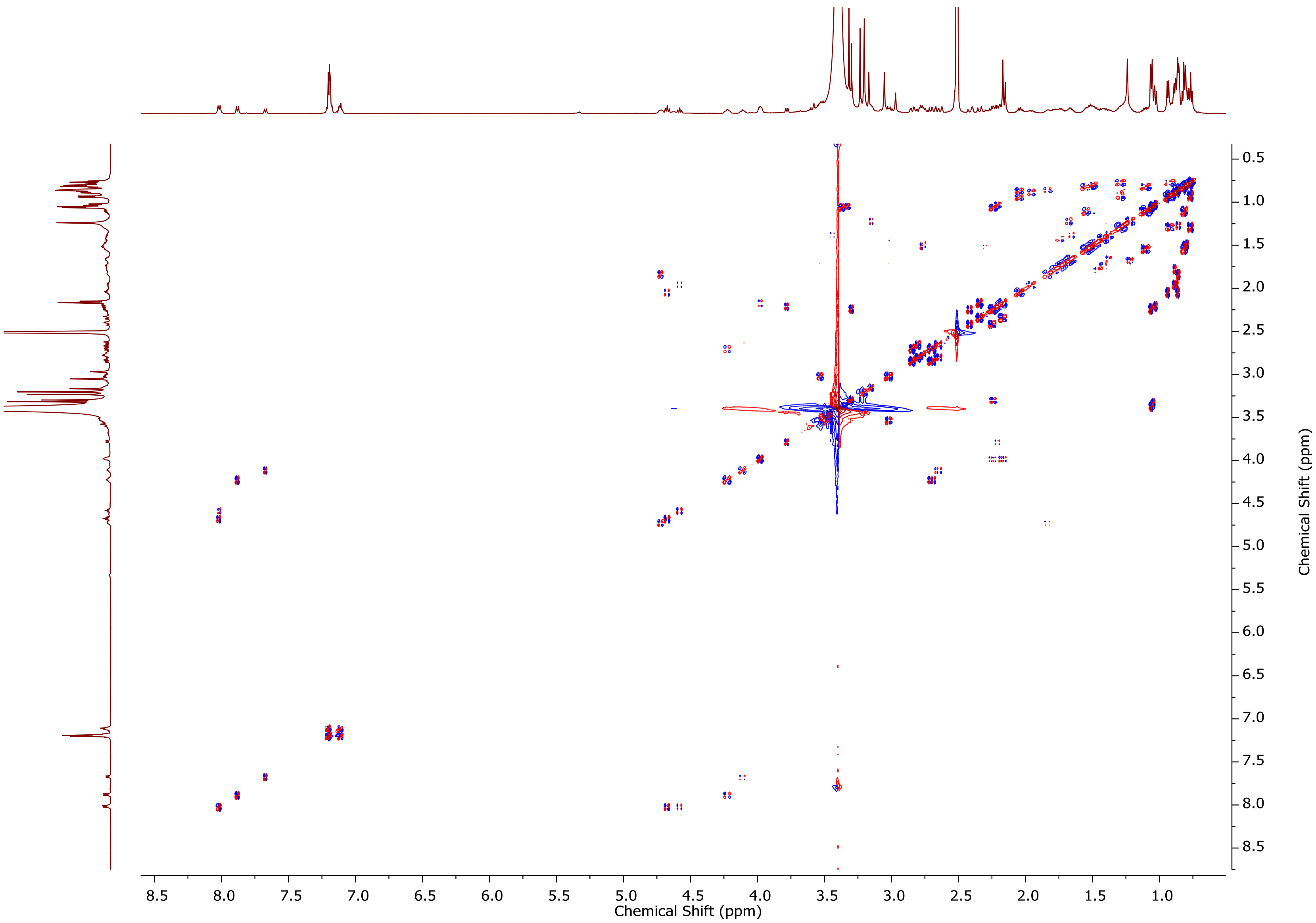


**Fig. S17.** DQF-COSY NMR spectrum of monomethyl-aetokthonostatin (**2**) in DMSO-$d_6$ (600 MHz).

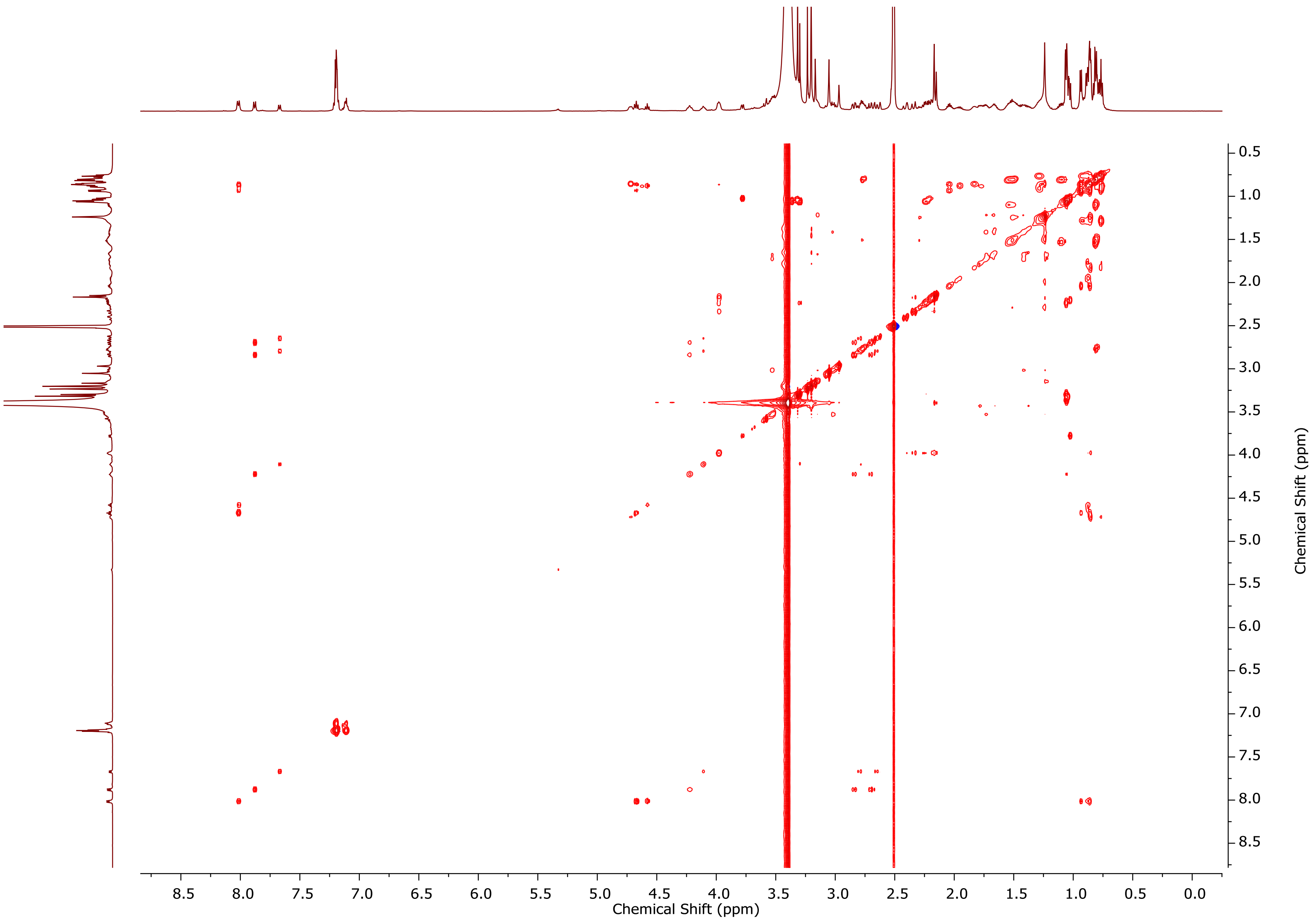


**Fig. S18.** TOCSY NMR spectrum of monomethyl-aetokthonostatin (**2**) in DMSO-$d_6$ (600 MHz).

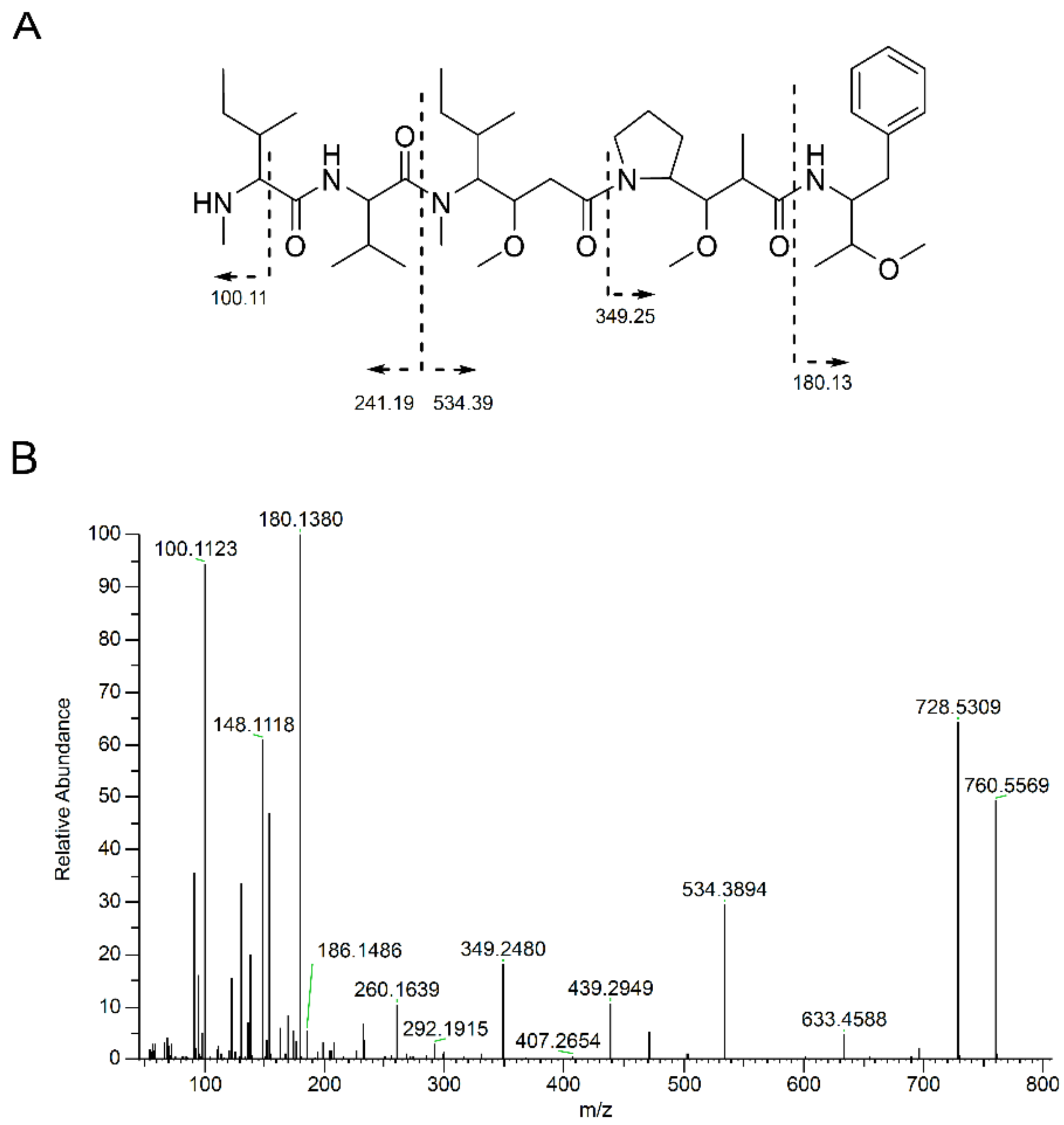


**Fig. S19. (A)** Key fragments and **(B)** MS/MS spectrum of monomethyl-aetokthonostatin (**2**).

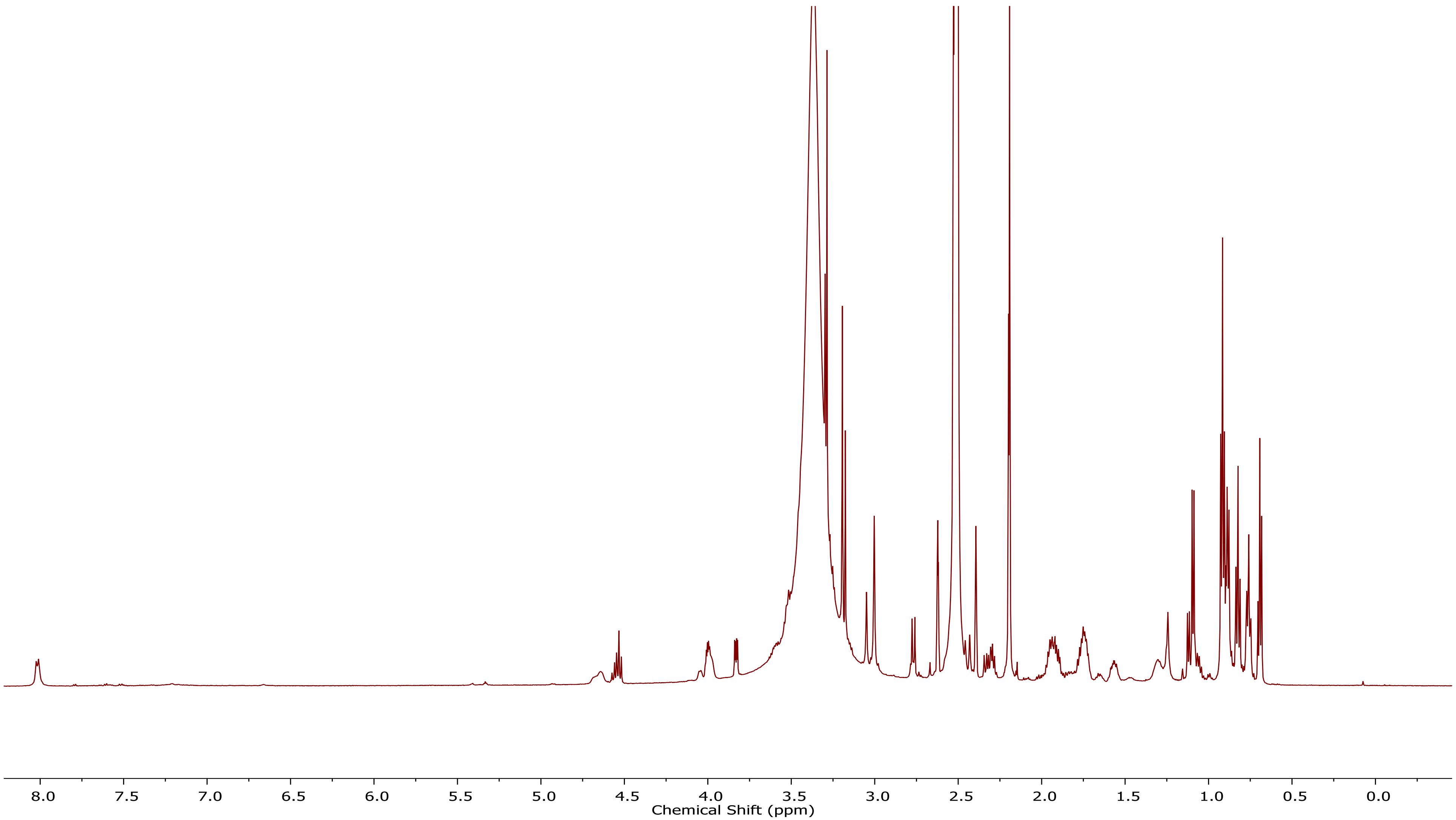


**Fig. S20.** $^1$H NMR spectrum of Des-Aph-aetokthonostatin (**3**) in DMSO-$d_6$ (600 MHz). See table S3 for a list of the observed chemical shifts.

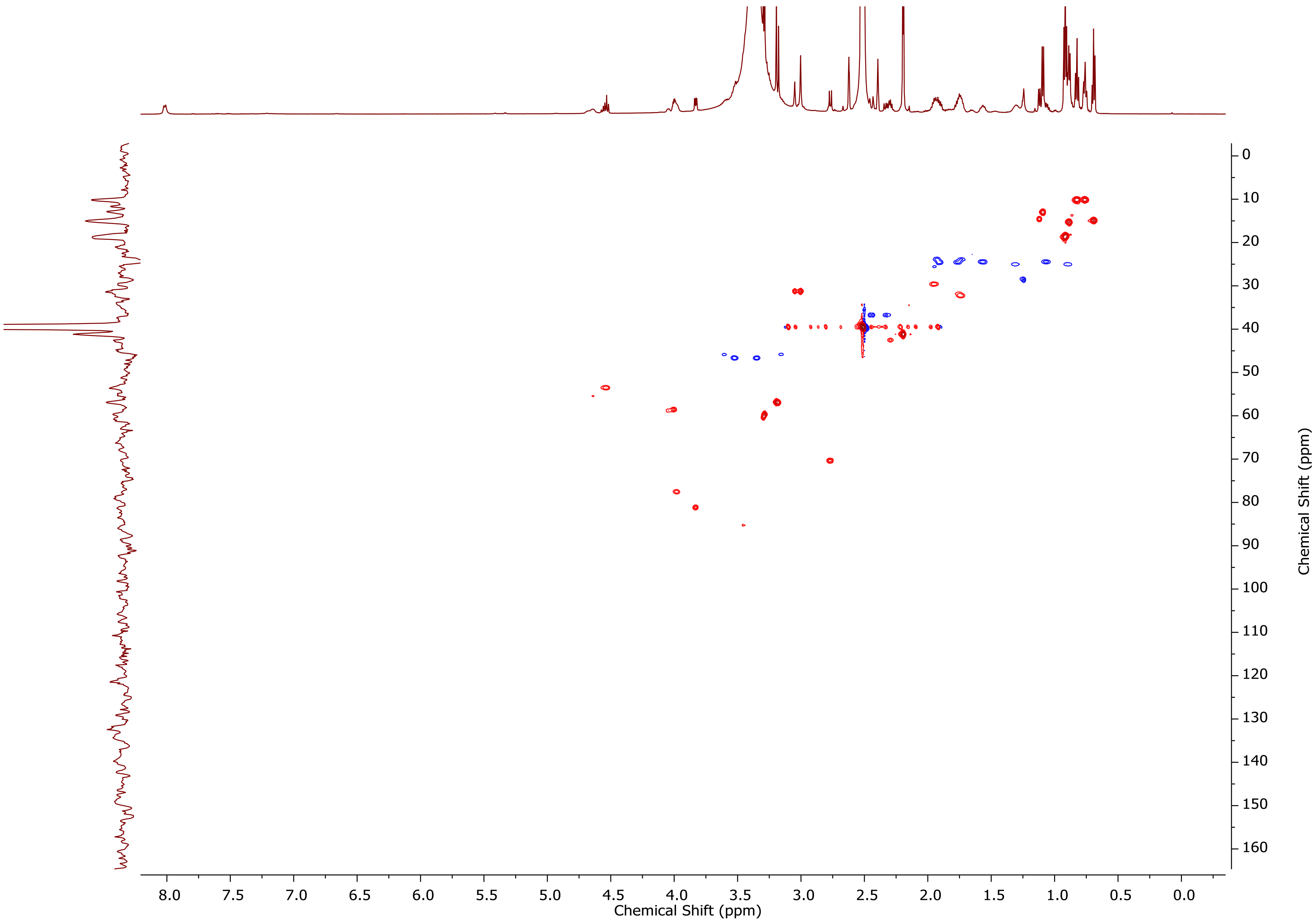


**Fig. S21.** $^{13}$C-HSQC NMR spectrum of Des-Aph-aetokthonostatin (**3**) in DMSO-$d_6$ (600 MHz). See table S3 for a list of the observed chemical shifts.

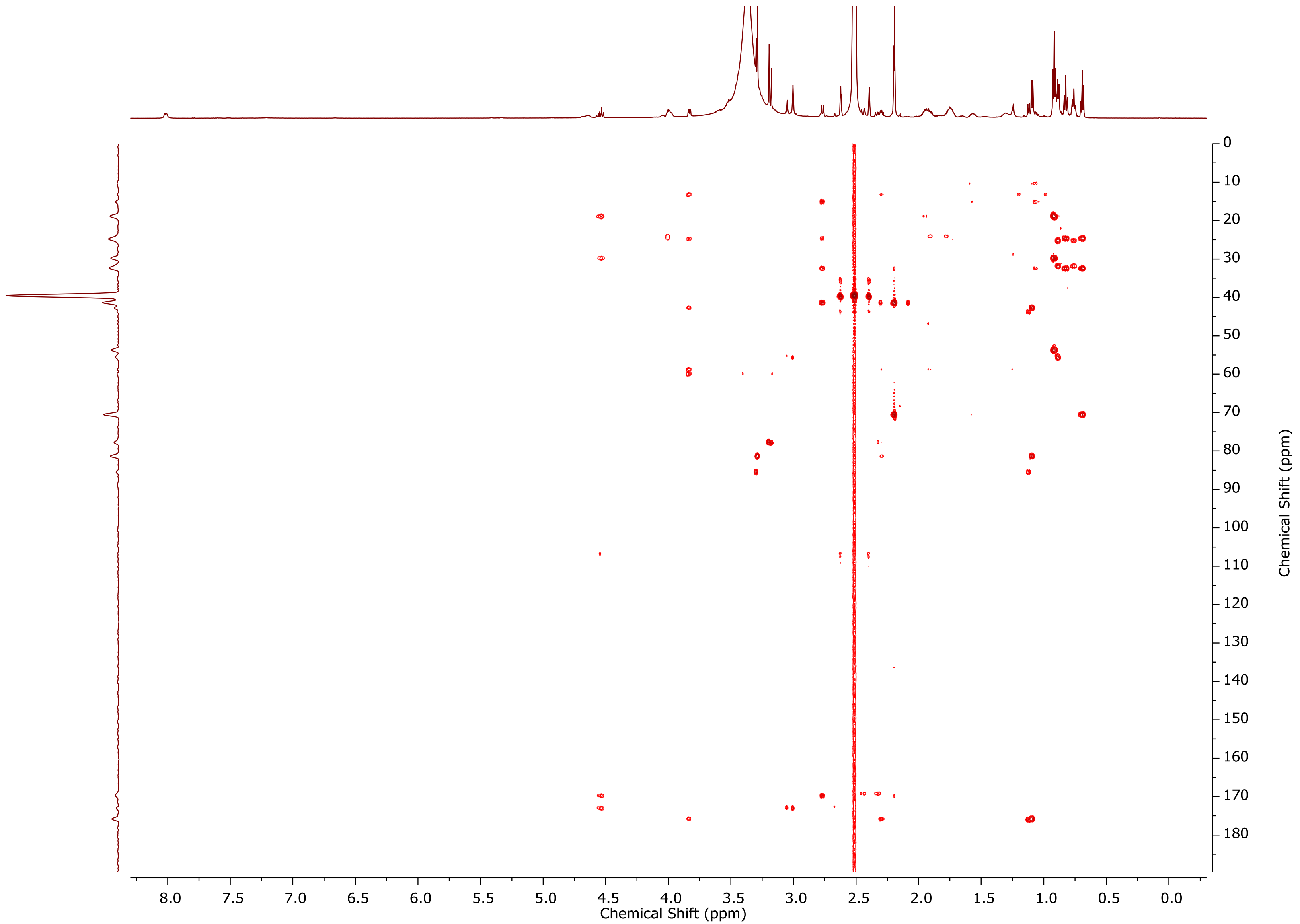


**Fig. S22.** $^{13}$C-HMBC NMR spectrum of Des-Aph-aetokthonostatin (**3**) in DMSO-$d_6$ (600 MHz). See table S3 for a list of the observed chemical shifts.

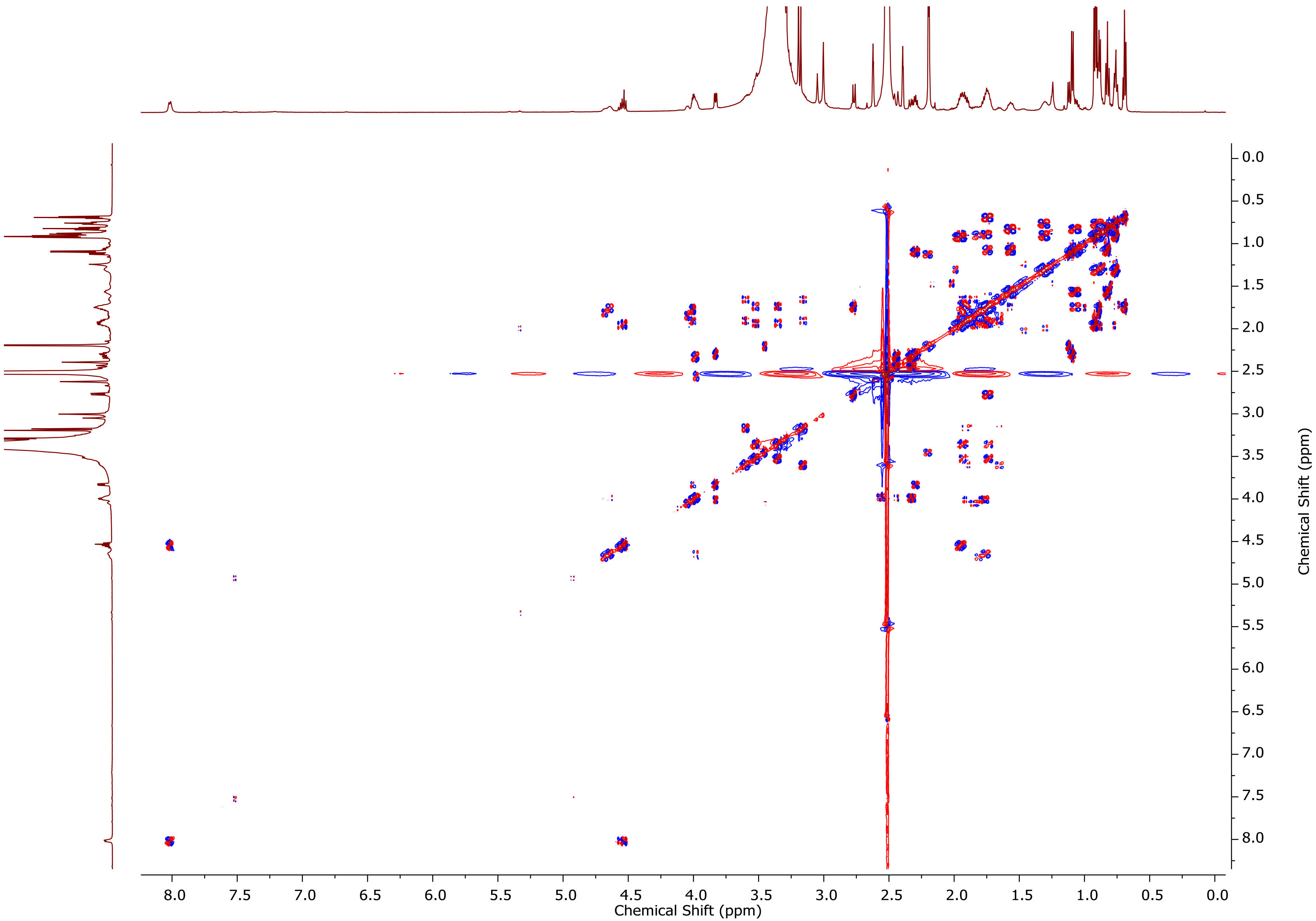


**Fig. S23.** DQF-COSY NMR spectrum of Des-Aph-aetokthonostatin (**3**) in DMSO-$d_6$ (600 MHz).

A

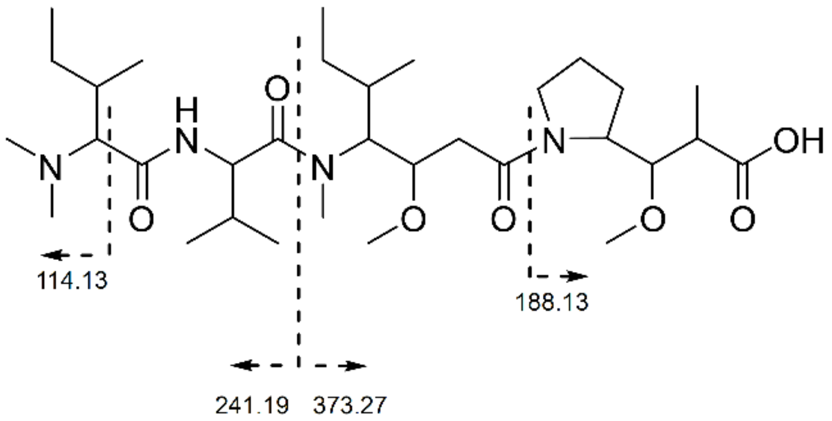


B

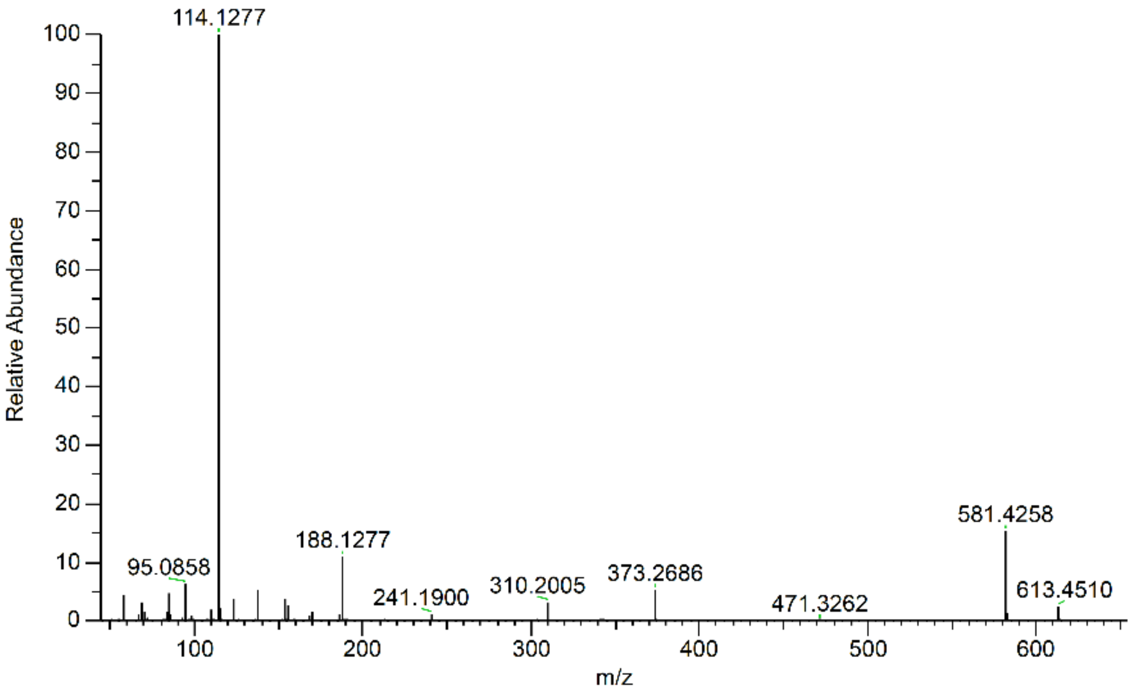


**Fig. S24. (A)** Key fragments and **(B)** MS/MS spectrum of Des-Aph-aetokthonostatin (**3**).

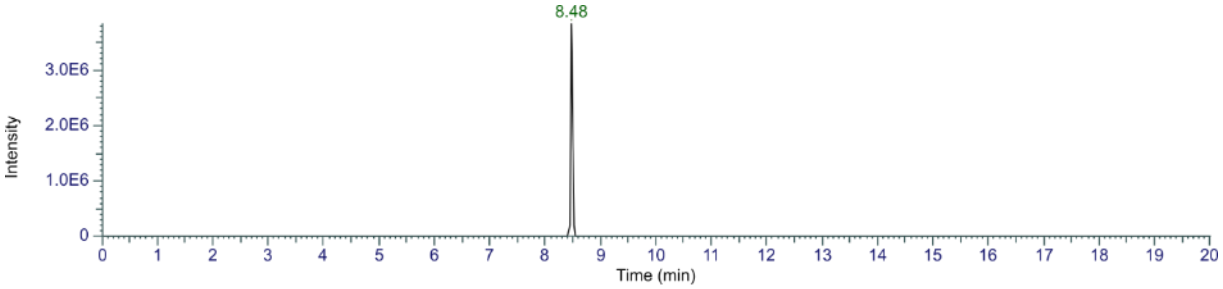


**Fig. S25.** Extracted ion chromatogram (EIC) of AEST (*m/z* 788.5) in the crude extract.

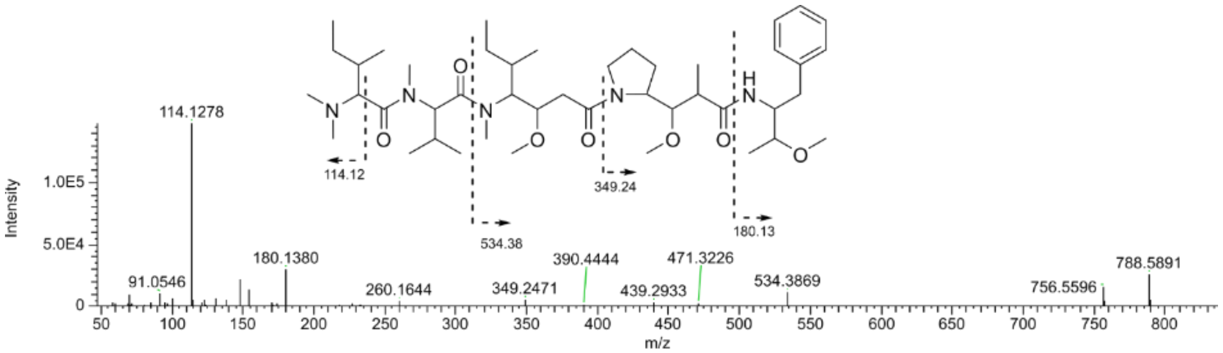


**Fig. S26.** Predicted structure based on MS/MS data of AEST at retention time 8.48 min.

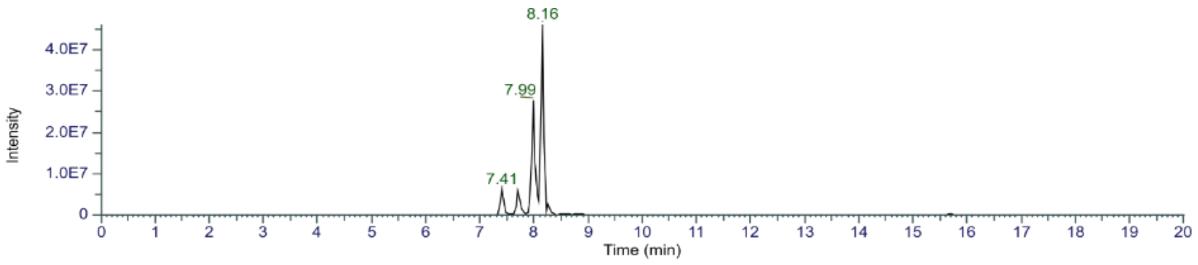

**Fig. S27.** Extracted ion chromatogram (EIC) of desmethyl AEST derivatives (*m/z* 760.5) in the crude extract.

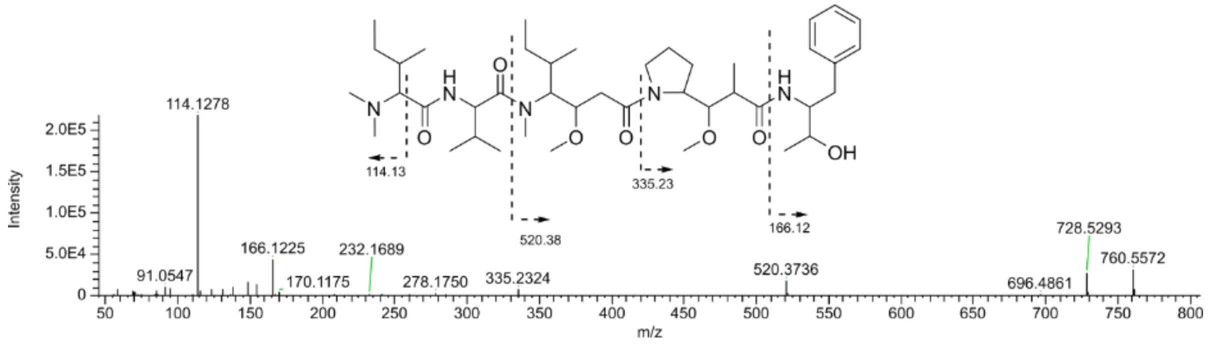

**Fig. S28.** Predicted structure based on MS/MS data of desmethyl AEST derivative at retention time 7.41 min.

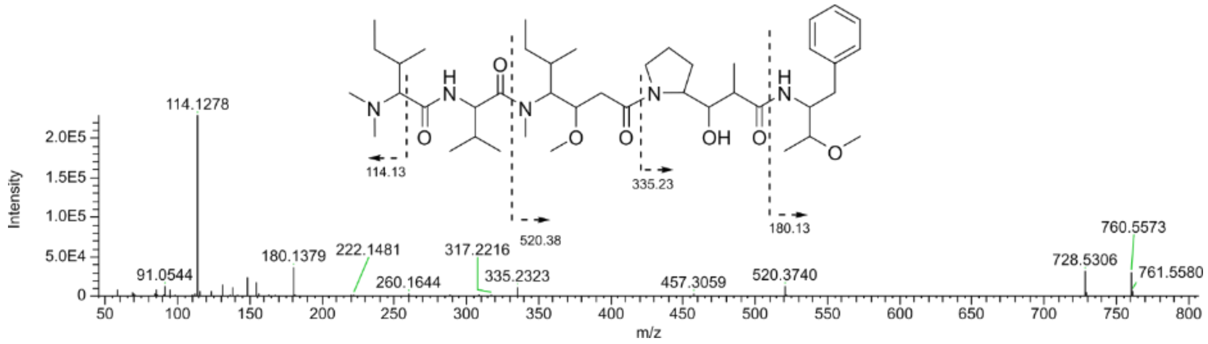

**Fig. S29.** Predicted structure based on MS/MS data of desmethyl AEST derivative at retention time 7.70 min.

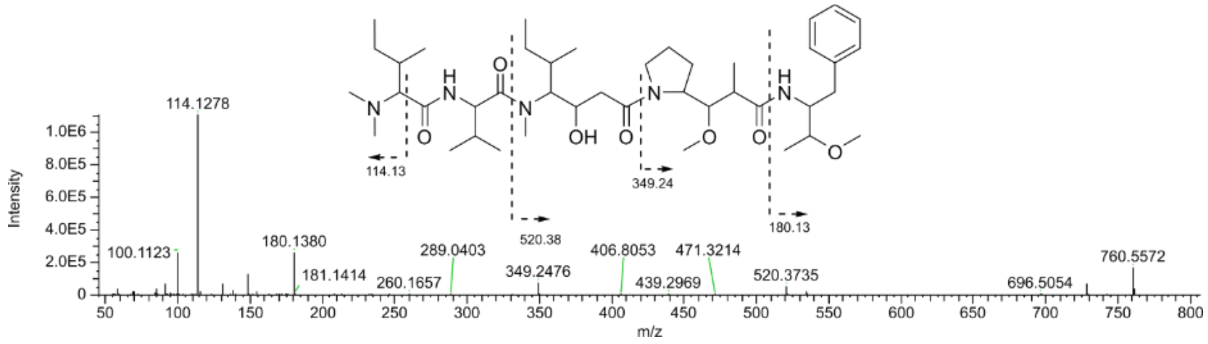

**Fig. S30.** Predicted structure based on MS/MS data of desmethyl AEST derivative at retention time 7.99.

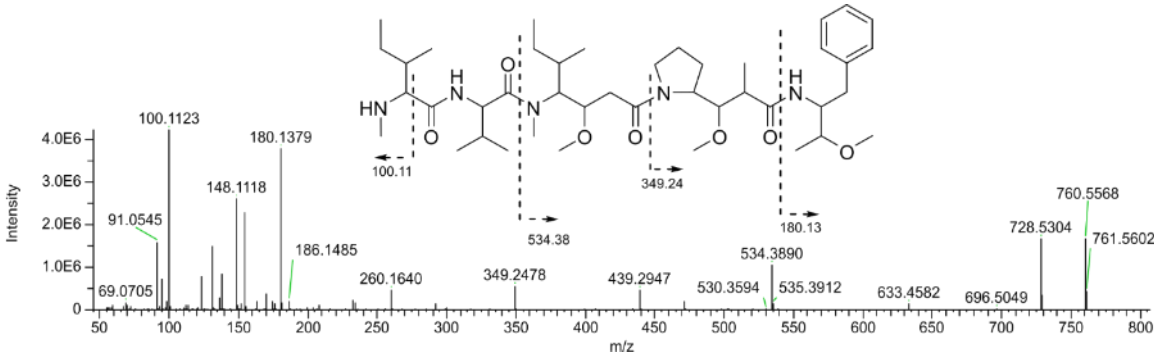

**Fig. S31.** Structure of MMAEST (**2**) and its MS/MS spectrum at retention time 8.16 min.

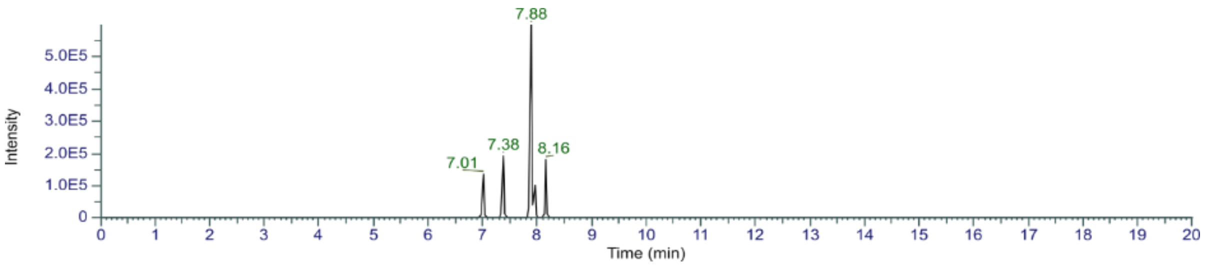


**Fig. S32.** Extracted ion chromatogram (EIC) of di-desmethyl AEST (DD-AEST) derivatives (*m/z* 746.5).

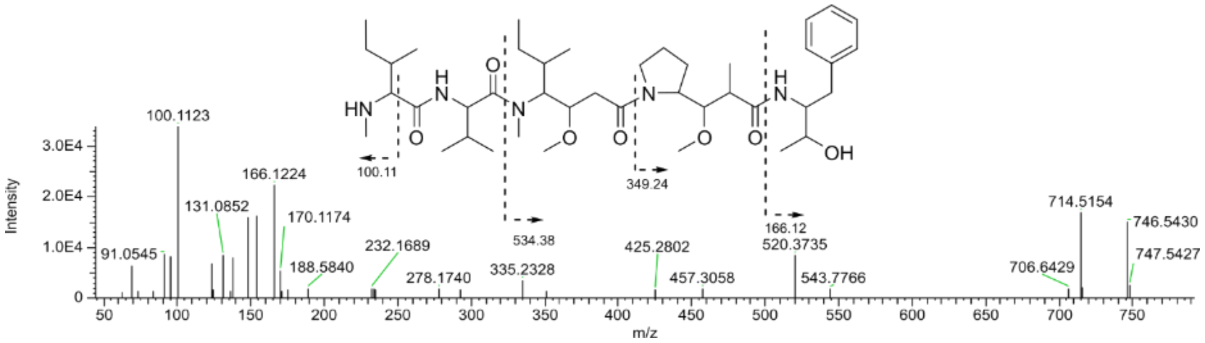


**Fig. S33.** Predicted structure based on MS/MS data of DD-AEST derivative at retention time 7.01 min.

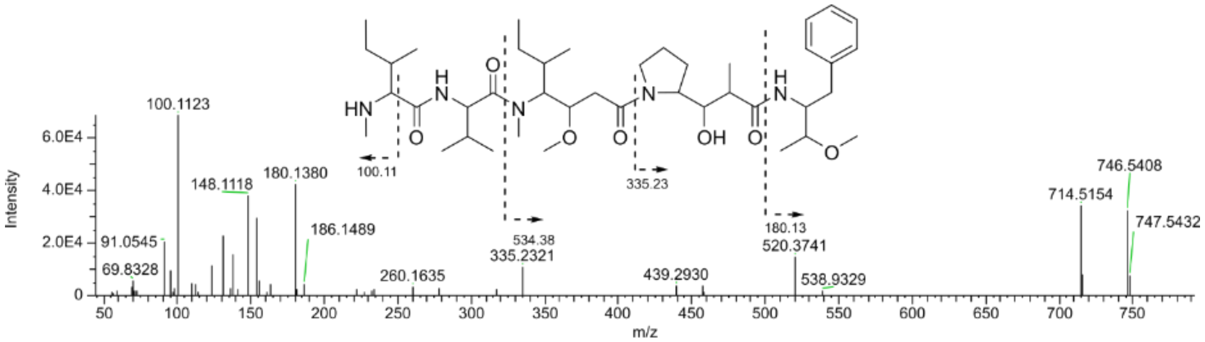


**Fig. S34.** Predicted structure based on MS/MS data of DD-AEST derivative at retention time 7.38 min.

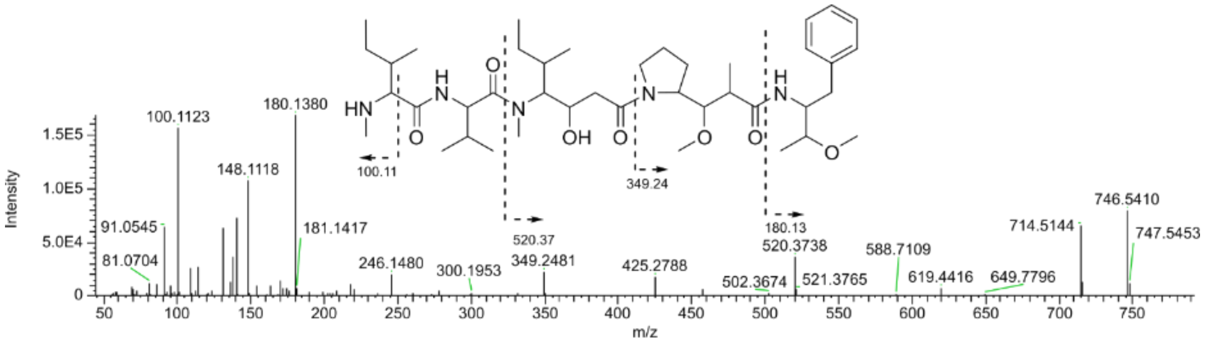


**Fig. S35.** Predicted structure based on MS/MS data of DD-AEST derivative at retention time 7.78 min.

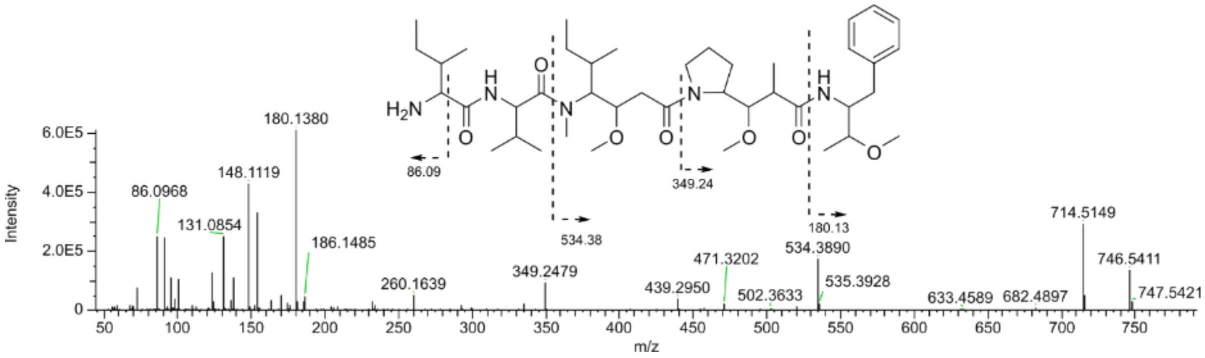


**Fig. S36.** Predicted structure based on MS/MS data of DD-AEST derivative at retention time 8.16 min.

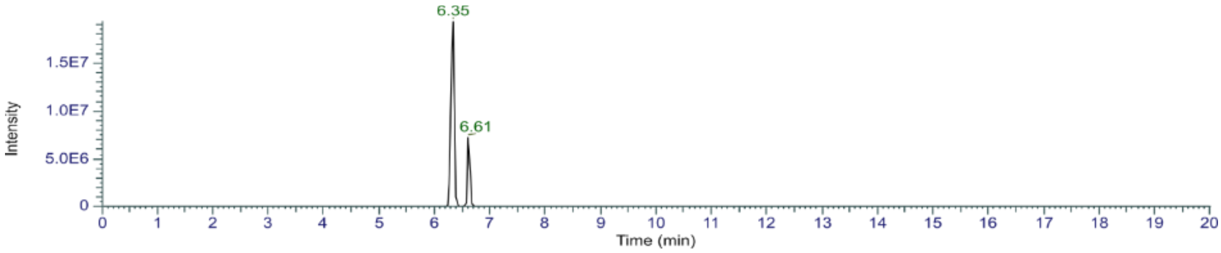


**Fig. S37.** Extracted ion chromatogram (EIC) of MM-tetrapeptide derivatives (*m/z* 599.4) in the crude extract.

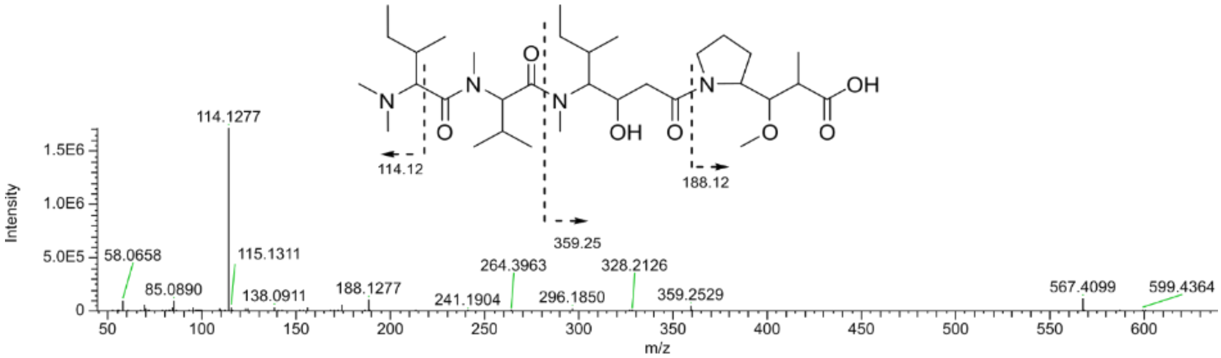


**Fig. S38.** Predicted structure based on MS/MS data of the MM-tetrapeptide derivative at retention time 6.35 min.

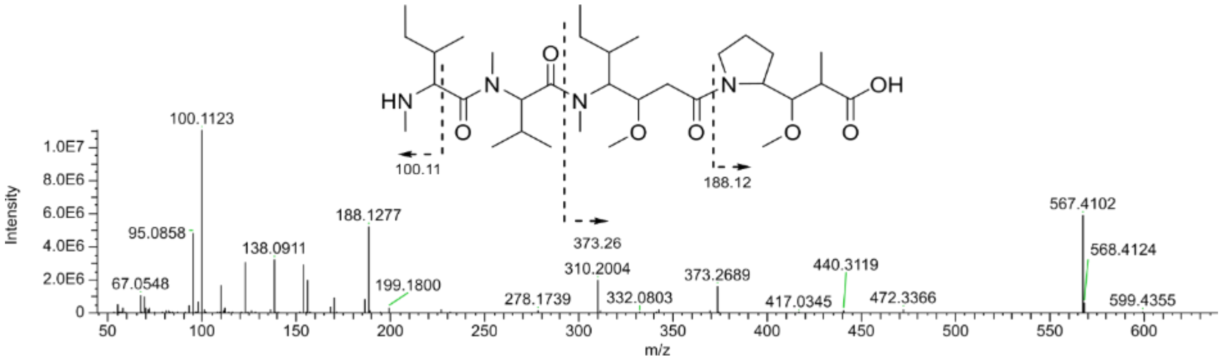


**Fig. S39.** Predicted structure based on MS/MS data of the MM-tetrapeptide derivative at retention time 6.6 min.

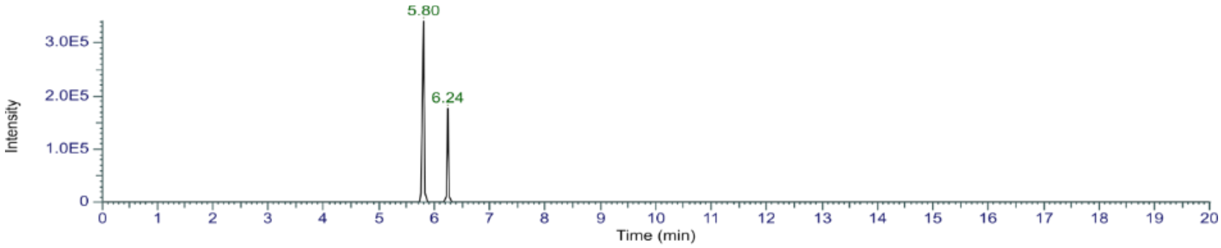


**Fig. S40.** Extracted ion chromatogram (EIC) of DD-tetrapeptide derivatives (*m/z* 585.4) in the crude extract

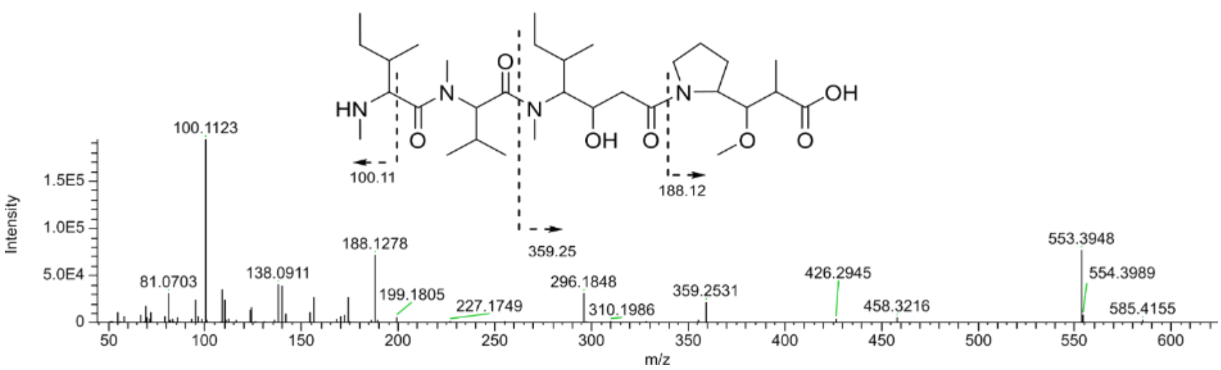


**Fig. S41.** Predicted structure based on MS/MS data of DD-tetrapeptide derivative at retention time 5.80 min.

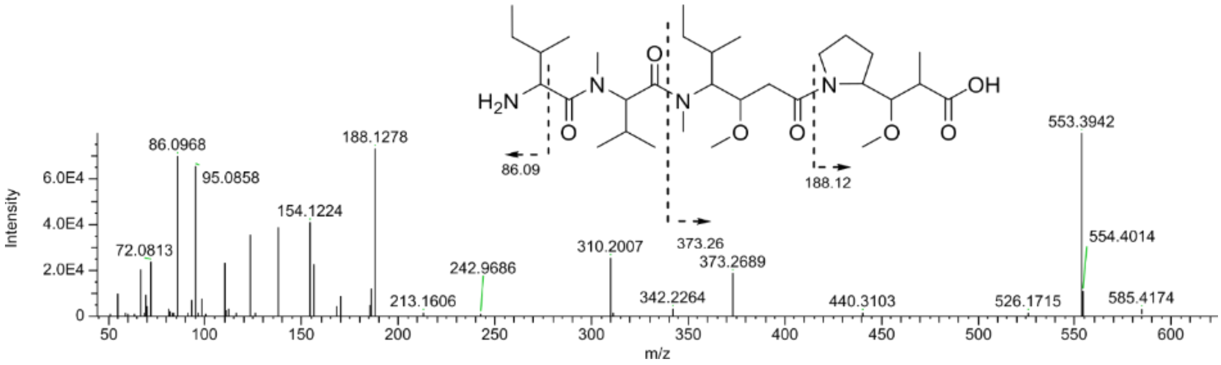


**Fig. S42.** Predicted structure based on MS/MS data of DD-tetrapeptide derivative at retention time 6.24 min.

### SUPPLEMENTARY FIGURES – Docking study and Molecular Dynamics simulation

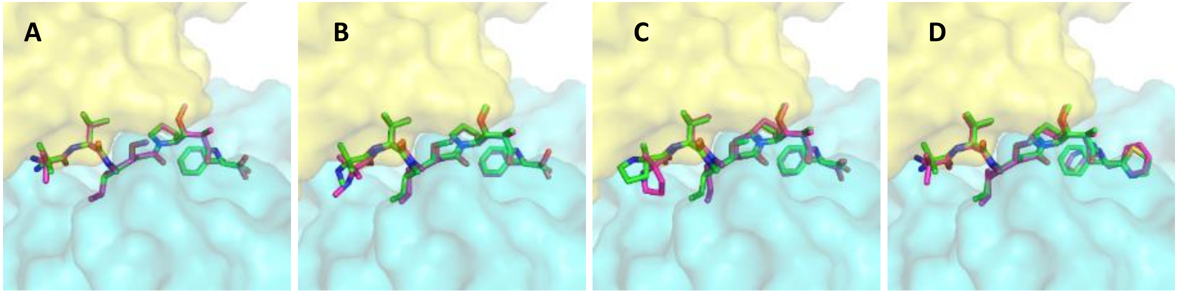


**Fig. S43.** Top-scored re- and cross-docking results in tubulin (PDB ID 4X1K) for inhibitor structures obtained from PDB IDs 4X1K (**A**), 4X1Y (**B**), 4X20 (**C**) and 4X1I (**D**). Reference binding poses are shown as magenta sticks, docking poses as green sticks. α-subunit of tubulin is shown as yellow surface, β-subunit appears as cyan surface. The cofactor GDP and the amino acid residues are not shown for the sake of clarity.

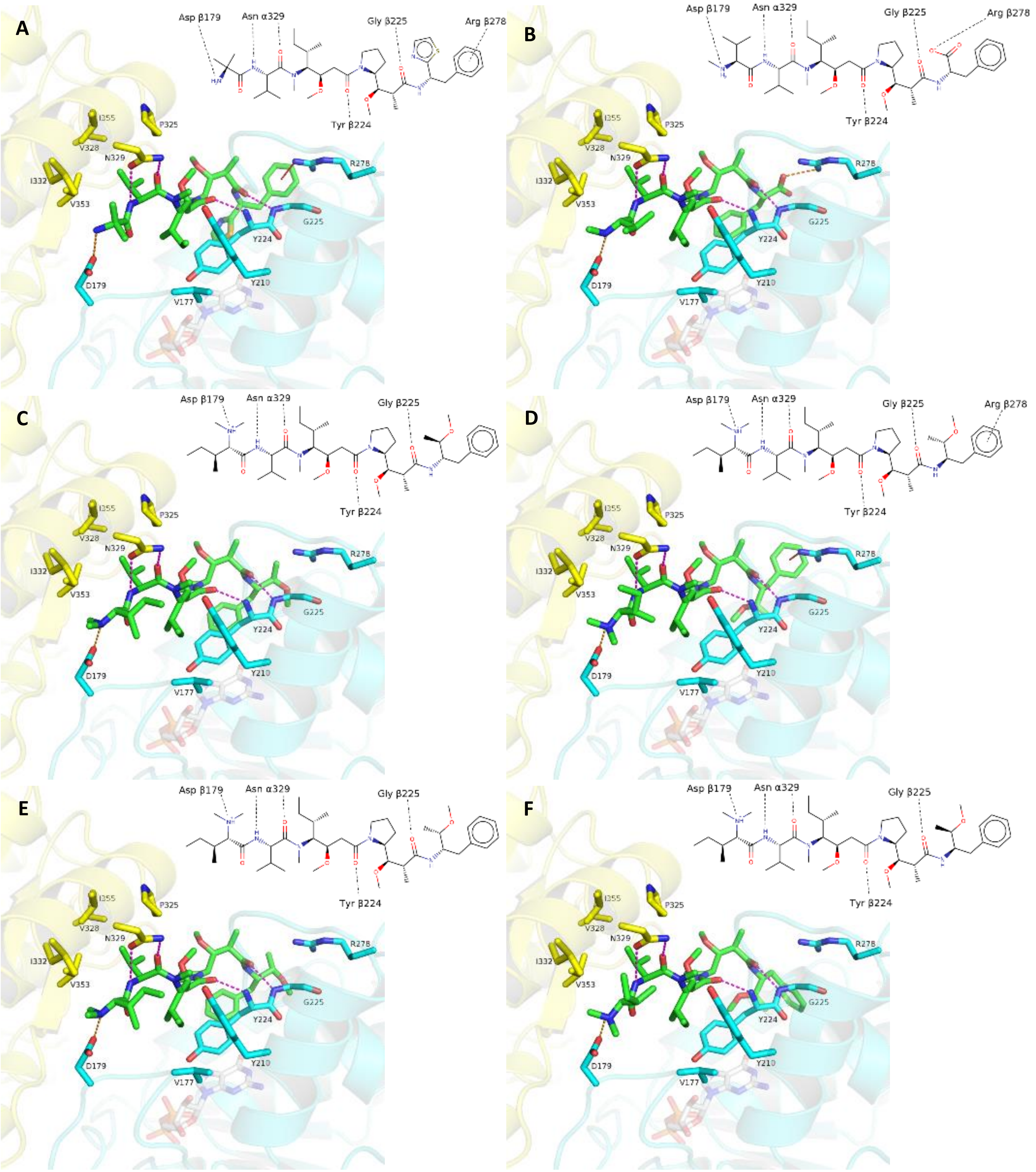


**Fig. S44.** Binding mode of the cocrystallized inhibitor found in 4X1I (**A**). Predicted binding modes of MMAF (**B**) and AEST (varied stereochemical configurations; **C** *(SR)*, **D** *(RS)*, **E** *(SS)*, **F** *(RR))*. in tubulin (PDB ID 4X1K). Tubulin α-subunit visualized as yellow cartoon, β-subunit in cyan. Interacting binding site residues represented as sticks, coloured the same way. Magenta: hydrogen bonds, orange: salt-bridges, red: cation-π interactions. Cofactor GDP: white, transparent sticks. Inhibitor structures with ligand-receptor-interactions depicted as 2D structures.

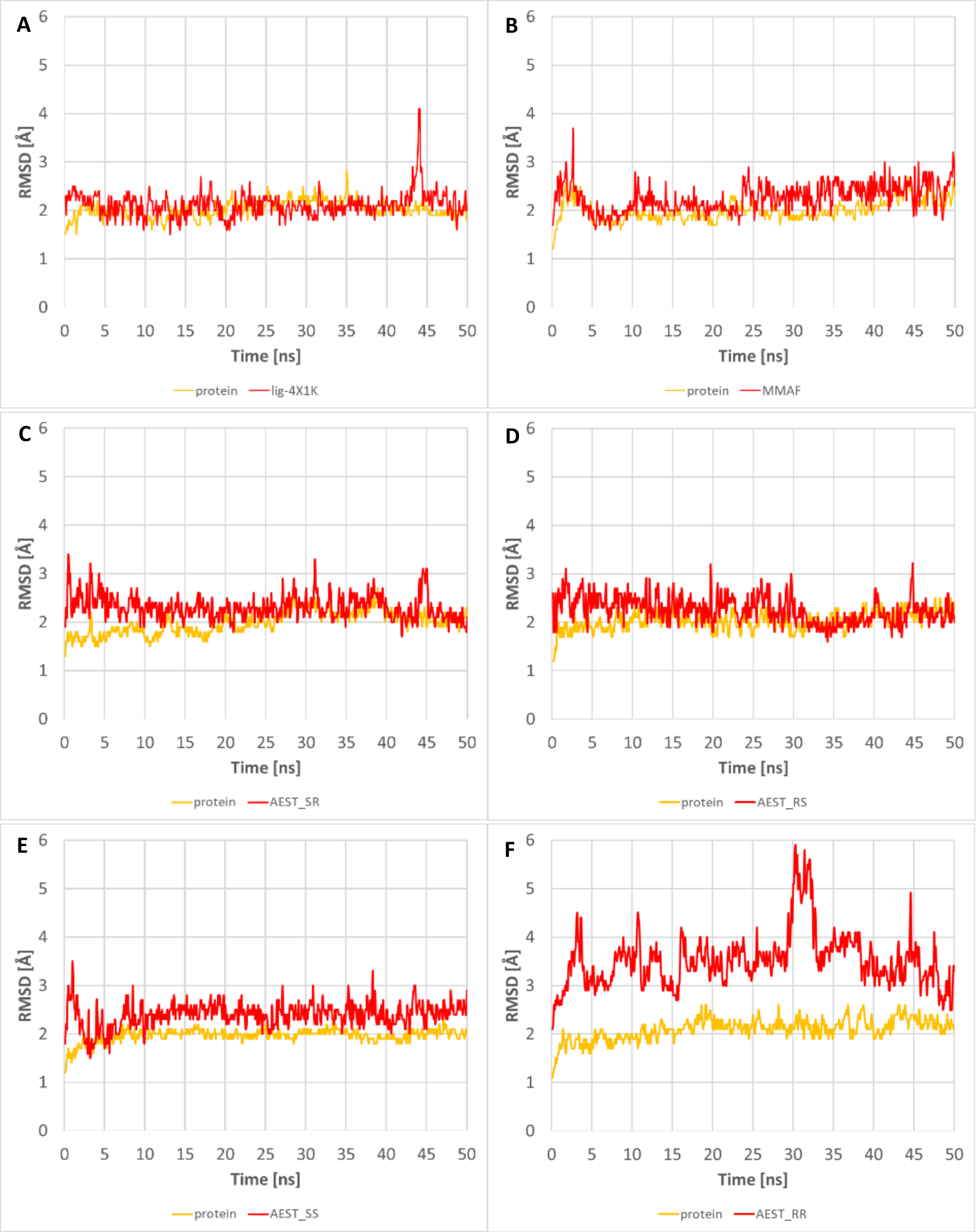


**Fig. S45.** RMSD plots of MD simulations performed using PDB ID 4X1K together with corresponding cocrystallized ligand (**A**) and generated docking poses for MMAF (**B**) and AEST (varied stereochemical configurations; **C** *(SR),* **D** *(RS),* **E** *(SS),* **F** *(RR)*). Protein RMSD graphs are colored in orange while inhibitor RMSD graphs appear in red.

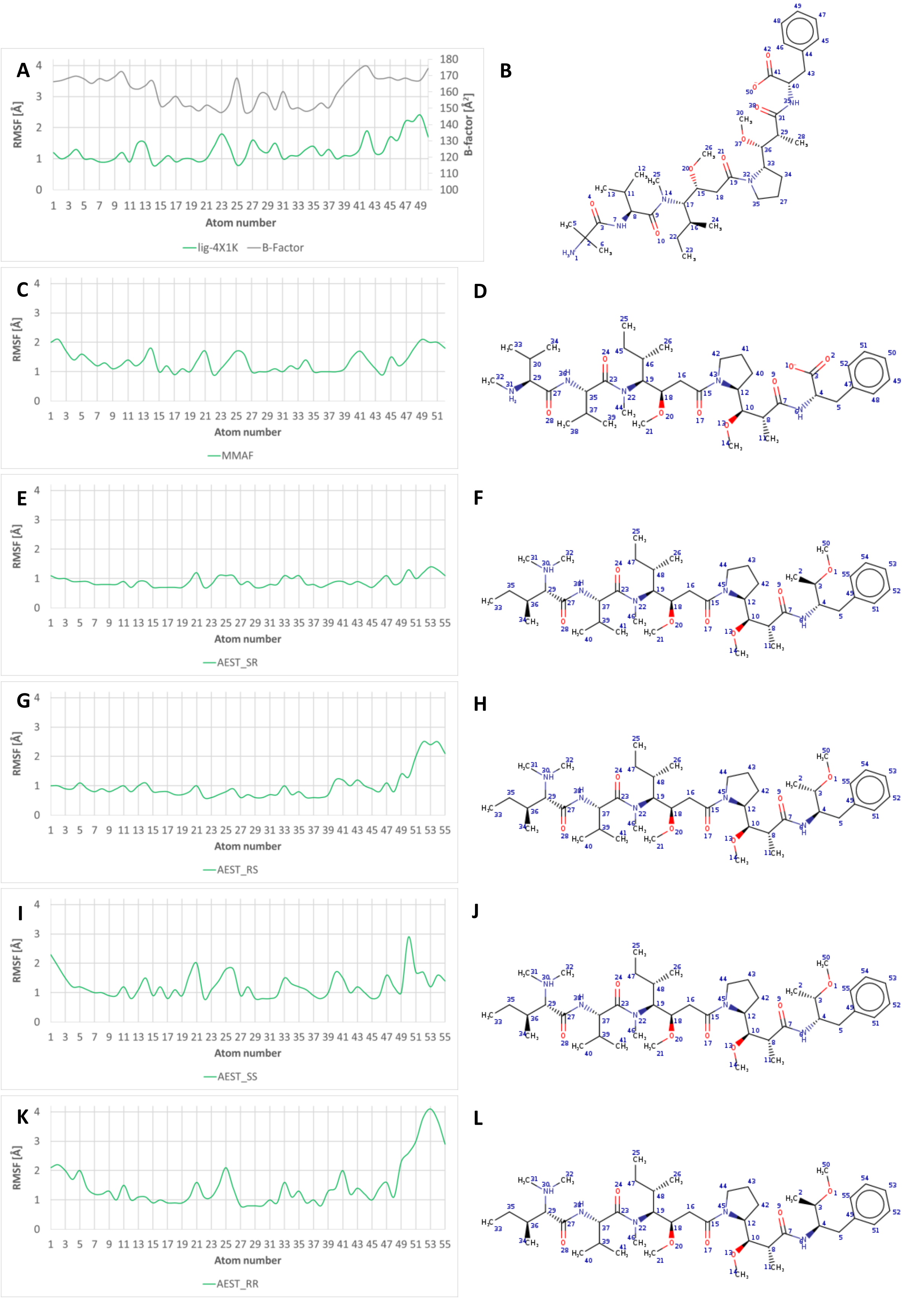


**Fig. S46.** RMSF plots of MD simulations performed using PDB ID 4X1K together with corresponding cocrystallized inhibitor (**A**) and generated docking poses for MMAF (**C**) and AEST (varied stereochemical configurations; **E** (*SR*), **G** (*RS*)**, I** (*SS*), **K** (*RR*). Plot A also depicts B-factor values in grey for comparison. The chemical structures of mentioned compounds next to the graphs allow an overview of atom number assignments (**B**, **D**, **F, H, J, L**).

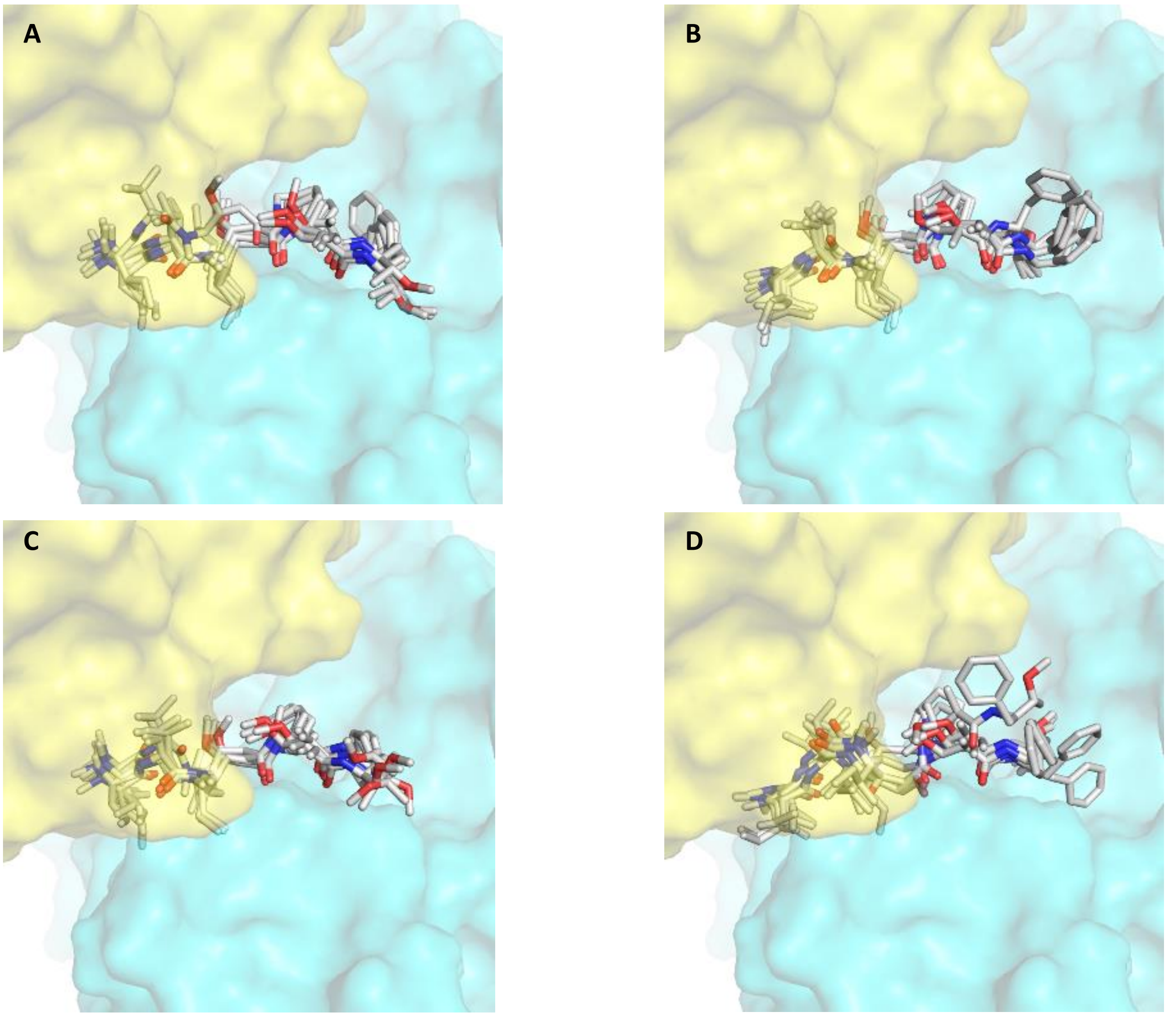


**Fig. S47.** Frames extracted from the MD simulations performed with AEST in different stereochemical configurations (**A**: *SR*, **B**: *RS*, **C**: *SS*, **D**: *RR*).

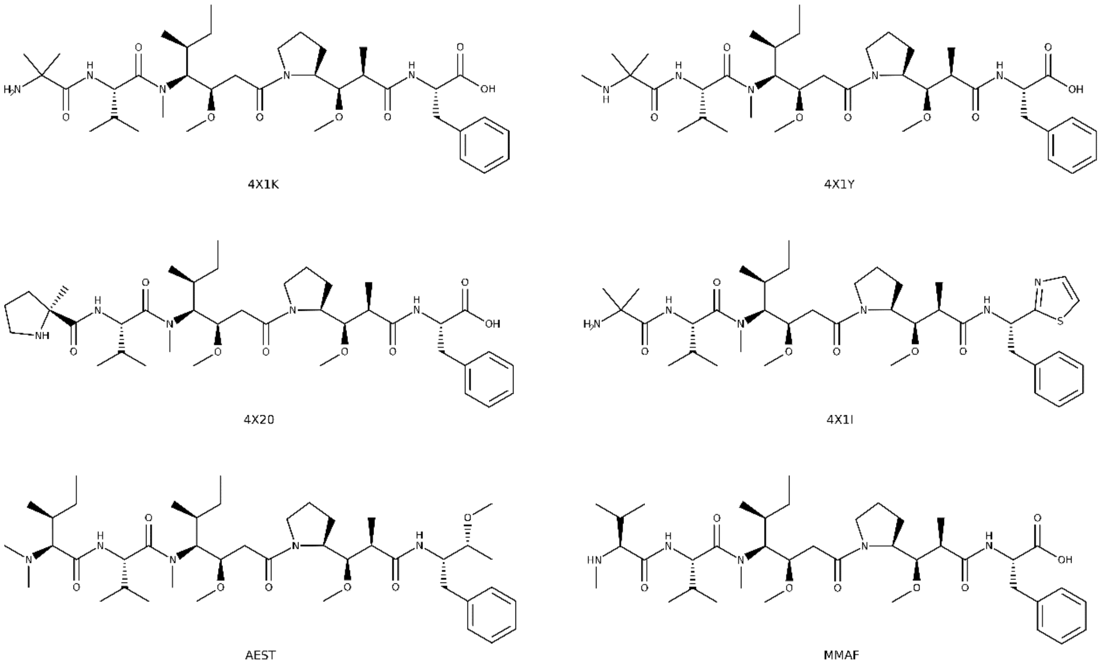


**Fig. S48.** Chemical structures of the cocrystallized inhibitors (PDB ID given) as well as of AEST (SR configuration) and MMAF for comparison.

### SUPPLEMENTARY FIGURES – Biosynthesis

**A**

*aesH*

CAGGAAAACAAAAGCGTGGGCGCGAGCCAGCTGACCGATCCGGTGAAAGATACCGTGAAAACCTTTTATAAC
AGCATTAACGATCATCTGAACACCAGCGGCTTTGGCGAATATAGCACCTTTCTGAACTGGGGCTATGTGAGCG
ATGGCAGCCCGGATTTTAGCAAAGTGGAACTGCCGGAACGCTGCCTGAACAAAAACTGCCTGAAACTGATTCT
GGAAGTGGTGGGCGATTGCGATCTGACCGGCCGCGATATTCTGGAAGTGGGCTGCGGTCGCGGCGGCAACAT
TCAGACCCTGGATAAATTTTTTAAACCGAAAGGCCTGACCGGCATTGATATTACCCCGGCGAGCATTGCGTTTT
GCAAAAAATATCTGGAAACCGAACGCCGCCGCTTTCTGGAAGGCGATGCGGAAGCGCTGCCGTTTGAAAACG
AAACCTTTGATGTGGTGATTAACGTGGAAAGCAGCAACGCGTATCCGAACCTGTTTAACTTTTATAGCCATGTG
AGCCGCGTGCTGAAACCGGGCGGCTATTTTCTGTATACCGATCTGATTAAAACCGAACTGATTGAAGATTGCAT
TCAGTATCTGGCGAAATGCGGCCTGGTGCTGGAAATTAACCGCGATATTACCAGCAACGTGCTGCTGAGCCGC
CAGGAAACCGCGGCGACCCAGATGAAAAGTTTTGGCTTTCAGAAAAGCGAAGTGGCGATGAGCAACGAAGA
AAAAAAAATTTTTGATTTTTTTCTGGCGCCGGAAGGCACCACCTTTTTTGAACAGATTCAGGATGGCCGCTTTA
GCTATAACATTTTTCGCTTTCGCAAAAAACGCGAATAA

**B**

*aesI*

GTGGAAAAAAGCGTGCCGGAAACCGCGCCGCAGGAACTGAAAAACCGCATTCAGCGCTTTTATGAATTTCTGA
ACACCCGCCTGAACGCGAGCATTTATGCGGAATATGCGATGTTTATGAACTGGGGCTATGAACATGATGAAAA
CCCGAGCTTTAGCCAGATTAAACTGCCGAACCATTATCTGAACAAATGCAGCACCCAGCTGGTGCTGGAACTG
ATTGGCGATTGCGATCTGACCGGCGCGACCGTGCTGGATGTGGGCTGCGGTCGCGGCGGCACCATTTATACCC
TGAACCATTTTTTTAACCCGAAACAGCTGATTGGCCTGGATATTACCGCGGCGAACATTGAATATTGCCGCAAA
AACGTGATTGGCGATCGCATTCAGTTTGTGCGCGGCGATGCGGAAAACCTGCCGTTTGCGAACGATGAATTTA
ACGTGGTGGTGAACATTGAAAGCAGCCATGGCTATCCGAACCGCCTGAAATTTTATCAGGAAGTGTATCGCGT
GCTGAAACCGGGCGGCAGCTTTCTGTATACCGATATTTTTCTGAGCCAGGAACTGGAAGGCTGCATTAACAGC
ATTCAGGAAATTGGCTTTGTGCTGGAAATTGATCGCGATATTACCAACAACGTGCTGCTGAGCCGCGAAAAAG
ATGCGGAATATCAGATTAACGCGTTTACCACCGAAGAAGATAACTGCGTGGATAACACCGAACTGGCGGAAG
ATATTGAAATGAGCGAATATGTGATGGCGCTGCCGAACAGCAACCTGTATGAAGGCATGCGCAAAAGCGAAT
ATTTTTATCGCATTTATCGCTTTAAAAAAAGCGAAGAAAACGTGGATAGCCTGAACCTGCAGCAGGAAAAAAA
CGAACCGGATGTGATTGAAAACGTGAAACTGCGCAGCCAGAAACAGAAAGAAGCGATTCGCCGCATTAACAG
CCGCAACAACAACCTGCATCAGTAA

**C**

*aesK*

GTGAACCTGGAAAGCGCGAGCAAATTTGATAGCTTTATTCTGCCGCCGGTGATGCGCGAACTGTTTGATCAGA
CCGGCTTTTTTAACTTTGGCTATTGGCTGAGCGATACCAAAAGCGCGCGCGAAGCGAGCGAAAACCTGATGGA
AAAACTGCTGGCGTTTATTCCGGAAAAAAAAGGCACCATTCTGGATGTGGCGTGCGGCCTGGGCGGCACCAC
CAACTATCTGCTGAAATATTATAGCCCGCTGGATATTGTGGGCATTAACATTAGCGCGAAACAGCTGGAAAAA
TGCCGCGATAACGCGCCGGGCTGCAAATTTATTGCGATGGATGCGGCGCAGATGGAATTTGAAGATAACAGC
TTTGATAACATTATTTGCGTGGAAGCGGCGTTTCATTTTTATACCCGCGAAAACTTTCTGCAGGAAGCGTGGCG
CGTGCTGAAACCGGGCGGCTATCTGGTGCTGAGCGATATTATTTTTCCGAACATGGAAGCGCCGAACGATTGG
ATGATTCCGAGCCAGAACGATGTGAAAGATATTGATGAATATAAAGCGCTGTATCTGAAAGCGGGCTTTGAAG
AAGTGGAACTGCTGGATAGCACCCATCAGAGCTGGATTGAATTTCATCGCCATGTGAAAACCTGGAACCTGCA
GAACTATCCGGATCAGGTGACCGAAAGCCATGTGCAGGAACTGGATGAACGCCTGAACTGGGGCGTGCTGTA
TCCGCTGGTGAGCGCGCGCAAAATTTAA

**Fig. S49.** Sequences of methyltransferase genes *aesH* (A), *aes I* (B) and *aes K* (C) optimized for heterologous expression in *E. coli*. Codon optimization was performed using the Geneious 9.1 software.

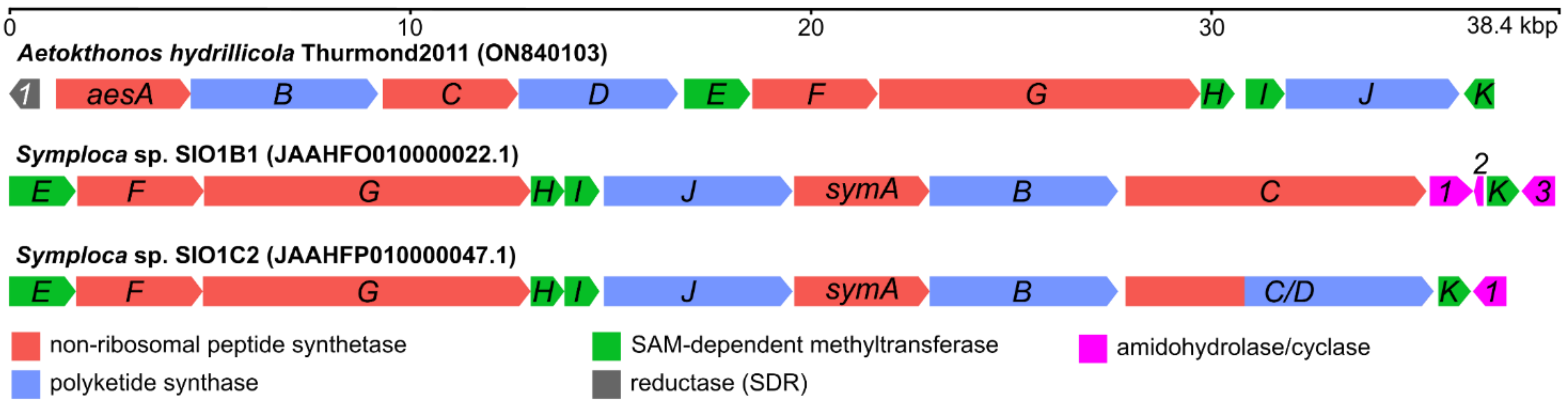


**Fig. S50.** Gene map of the aetokthonostatin biosynthetic gene cluster compared to homologous putative biosynthetic gene clusters of dolastatin family compounds found in metagenome-assembled genomes of environmental *Symploca* sp. derived from the NCBI nr database using BLAST search. SAM – S-adenosylmethionine; SDR – short-chain dehydrogenase/reductase. The genes identified in *Symploca* sp. are tentatively labelled *symA*–*K* as they presumably do not code for aetokthonostatins but rather symplostatins or their analogs (based on preliminary bioinformatic analysis).

**SUPPLEMENTARY FIGURES – *In vitro* enzyme assays**

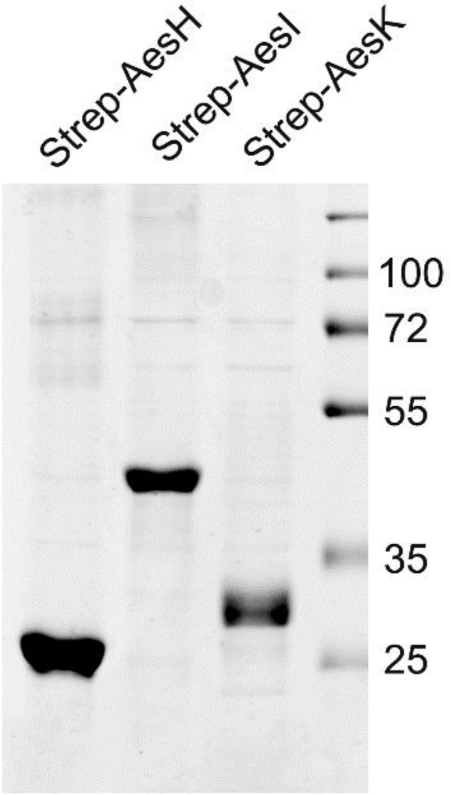


**Fig. S51.** SDS-PAGE of purified Strep-tagged AesH, AesI and AesK proteins. All the proteins were expressed in *E. coli* BL21(DE3) and purified using Strep-Tactin affinity chromatography. From the total elution volume (500 µL), 10 µL were separated on a 10% SDS-PAGE gel and stained with Coomassie blue.

*In vitro* reactions were carried out using the purified methyl transferases Strep-AesH, -AesI, and -AesK. The following figures show the results of these experiments. Heat-inactivated enzyme (left column) and a blank reaction without enzyme addition (center column) were compared with a reaction containing freshly purified enzyme (right column). The enzymes were incubated with the potential substrates MMAEST (upper row) and MMAF (lower row). In case these compounds are substrates of the respective methyl transferase, the respective methylated reaction products AEST and AF should be detectable by HPLC-MS analysis. Thus, the extracted ion chromatograms show the ions for MMAEST ($[M+H]^+$ 760.57 Da, shown in black) and AEST ($[M+H]^+$774.58 Da, shown in red) in the upper row, and MMAF ($[M+H]^+$732.49 Da, shown in black) and AF ($[M+H]^+$746.50 Da, shown in red) in the lower row. It can clearly be seen that neither Strep-AesH (Fig. S49) nor Strep-AesI (Fig. S50) are able to methylate MMAEST or MMAF. However, Strep-AesK can methylate both compounds (Fig. S51). The position of the methylation of MMAF could be confirmed to be located at the N-terminal amino acid (Fig. S52). However, Strep-AesK is not able to methylate the respective single N-terminal amino acids *N*-MeIle or *N*-MeVal (Fig. S53).

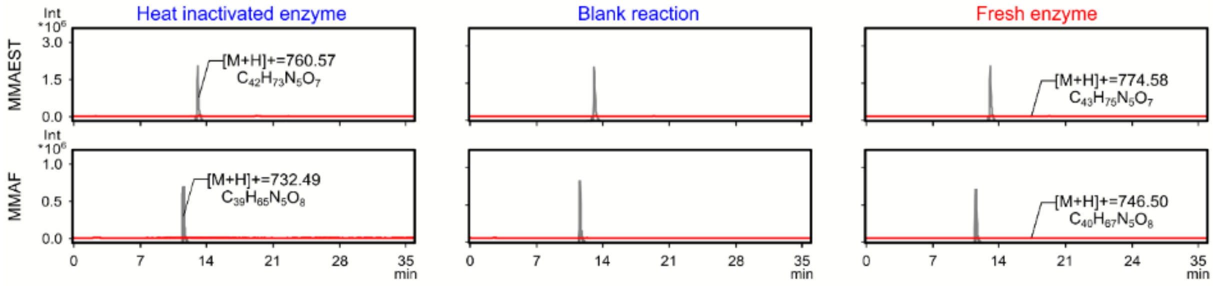


**Fig. S52**. EICs of *in vitro* reactions with Strep-AesH. No reaction products can be detected.

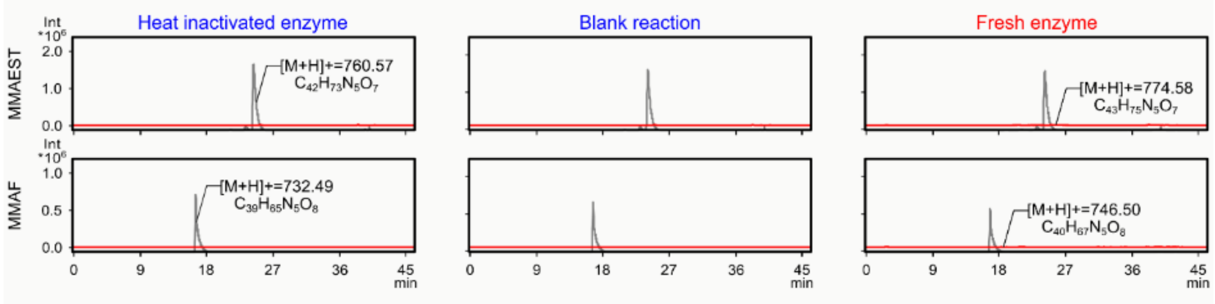


**Fig. S53**. EICs of *in vitro* reactions with Strep-AesI. No reaction products can be detected.

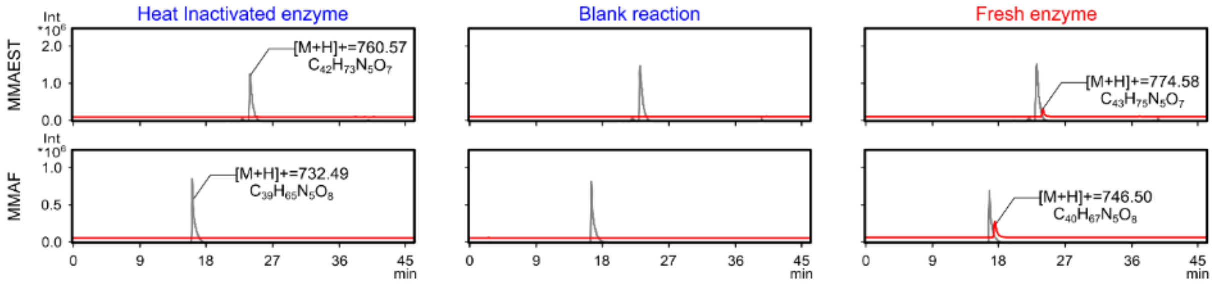


**Fig. S54.** EICs of *in vitro* reactions with Strep-AesK. Reaction products can be detected.

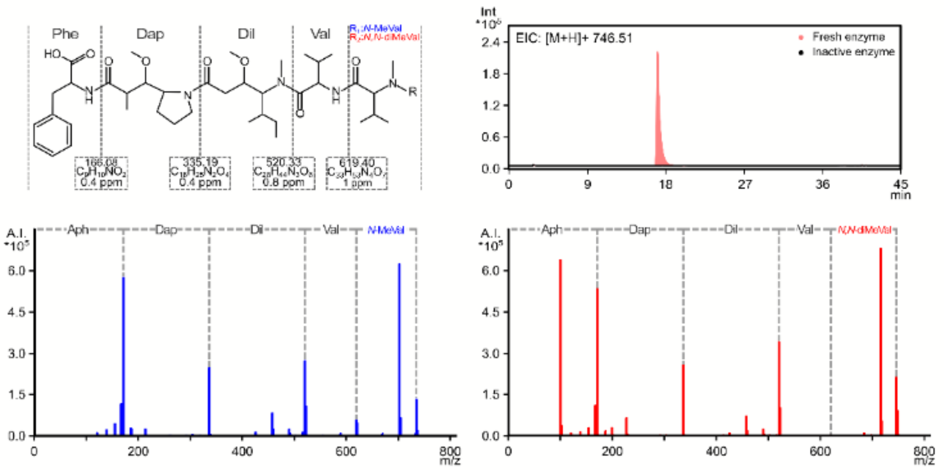


**Fig. S55**. Comparison of the structures of MMAF and AF (top left) showing that they only differ by methylation of their N-terminal amino acid (*N,N*-diMeVal for AF and *N*-MeVal for MMAF). HPLC-HRMS/MS analysis (extracted ion chromatograms) of AF content in reaction mixtures of MMAF incubated with heat-inactivated or fresh Strep-AesK (top right) show that the inactivated enzyme (black) had no effect on the substrate while the fresh Strep-AesK methylated MMAF (red). Comparison of the MS/MS spectra of MMAF (bottom left) versus the product of the fresh enzyme reaction (bottom right) shows that AesK methylated the N-terminus of MMAF, producing AF. Abbreviations: Dap, Dolaproine; Dil, Dolaisoleuine; *N*-MeVal, *N*-methylvaline; *N,N*-diMeVal, *N,N*-dimethylvaline.

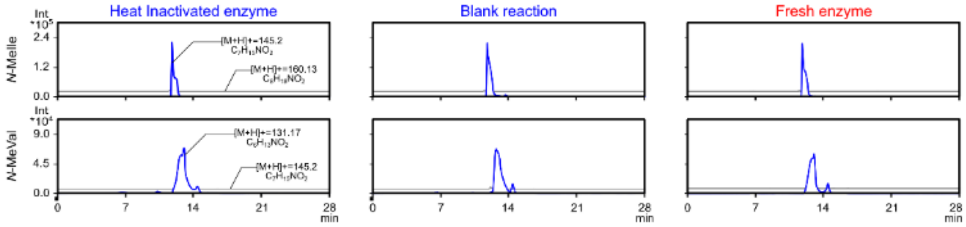


**Fig. S56.** EICs of *in vitro* reactions with Strep-AesK. Columns as described above; upper row: substrate *N*-MeIle, lower row substrate *N*-MeVal. No reaction products can be detected.
